# High-resolution yeast quiescence profiling in human-like media reveals interacting influences of auxotrophy and nutrient availability

**DOI:** 10.1101/2020.05.25.114801

**Authors:** Sean M. Santos, Samantha Laflin, Audrie Broadway, Cosby Burnett, Joline Hartheimer, John Rodgers, Daniel L. Smith, John L. Hartman

## Abstract

Yeast cells survive in stationary phase culture by entering quiescence, which is measured by colony forming capacity upon nutrient re-exposure. Yeast chronological lifespan (CLS) studies, employing the comprehensive collection of gene knockout strains, have correlated weakly between independent laboratories, which is hypothesized to reflect differential interaction between the deleted genes, auxotrophy, media composition and other assay conditions influencing quiescence. This hypothesis was investigated by high-throughput quiescence profiling of the parental prototrophic strain, from which the gene deletion strain libraries were constructed, and all possible auxotrophic allele combinations in that background. Defined media resembling human cell culture media promoted long-term quiescence, and was used to assess effects of glucose, ammonium sulfate, auxotrophic nutrient availability, Target of Rapamycin signaling, and replication stress. Frequent, high-replicate measurements of colony forming capacity from cultures aged past 60 days provided profiles of quiescence phenomena such as gasping and hormesis. Media acidification was assayed in parallel to assess correlation. Influences of leucine, methionine, glucose, and ammonium sulfate metabolism were clarified, and a role for lysine metabolism newly characterized, while histidine and uracil perturbations had less impact. Interactions occurred between glucose, ammonium sulfate, auxotrophy, auxotrophic nutrient limitation, aeration, TOR signaling, and/or replication stress. Weak correlation existed between media acidification and maintenance of quiescence. In summary, experimental factors, uncontrolled across previous genome-wide yeast CLS studies, influence quiescence and interact extensively, revealing quiescence as a complex metabolic and developmental process that should be studied in a prototrophic context, omitting ammonium sulfate from defined media, and employing highly replicable protocols.

## Introduction

Cell quiescence is defined by adaptive cell cycle exit (G0), prior to the G1-S transition, which preserves viability as evidenced by later cell cycle re-entry (Hartwell 1974). Quiescence occurs in yeast cultures when media nutrients are depleted, and is assayed by colony-forming capacity upon replenishment of nutrients (Lillie and Pringle 1980). Quiescence underlies yeast chronological lifespan (Gray et al. 2004; Longo et al. 2012; MacLean et al. 2001). Additional stimuli elicit quiescence in multicellular organisms (Mohammad et al. 2019). However, much remains to be learned about cellular processes that enable quiescence in yeast and humans, which quiescence mechanisms are conserved through evolution, and how quiescence contributes to longevity (Dhawan and Laxman 2015). Understanding the influence of quiescence mechanisms on organismal function is fundamental to aging biology, for example in stem cell function, and the developmental interplay with senescence, apoptosis and necrosis (Cho et al. 2019). In these regards, yeast is a useful genetic model for unbiased discovery and hypothesis generation about human conditions (Hartman IV et al. 2001; Hartman IV et al. 2015; Hartwell 2002). It is well positioned to inform eukaryotic quiescence mechanisms linking nutrient signaling, quiescence, and longevity in multicellular organisms (Dhawan and Laxman 2015; Templeman and Murphy 2018). Although systems level analysis of quiescence, uniquely obtainable through yeast genetics, could lead to a more global understanding of how the functional integration of genetic and metabolic networks is conserved through evolution, different genome-wide analyses of CLS in yeast knockout (**YKO**) strain libraries, consisting of ~5000 genes/strains lacked consensus (Fabrizio et al. 2010; Matecic et al. 2010; Powers et al. 2006; Smith et al. 2016). To help explain this complex outcome, we investigated plausible genetic and nutrient causes for weak correlation, using a quantitative high throughput colony forming capacity assay and highly controlled experimental designs to characterize effects on quiescence of interaction between auxotrophic alleles, media composition, aeration, media acidification and other perturbations known to influence yeast CLS.

The YKO collections were engineered on the FY4 genetic background with *leu2*Δ0, *ura3*Δ0, and *his3*Δ1 (Brachmann et al. 1998; Winston et al. 1995) deletion alleles in both the haploid *MATα* (BY4741) and *MATa* (BY4742) YKO library backgrounds. Library-specific *met17*Δ0 and *lys2*Δ0 mutations were introduced in to BY4741 and BY4742, respectively (Giaever et al. 2002; Winzeler et al. 1999). Limitation of leucine and other auxotrophic nutrients has been reported to reduce survival in stationary phase (Alvers et al. 2009; Aris et al. 2013; Boer et al. 2008; Gomes et al. 2007; Lillie and Pringle 1980). In contrast to other amino acids, methionine limitation in the context of *met17*Δ0 auxotrophy increases chronological survival in yeast and other eukaryotes (Johnson and Johnson 2014; Lee et al. 2016; Orentreich et al. 1993; Ruckenstuhl et al. 2014; Unger and Hartwell 1976). However, effects of combinations of auxotrophic mutations have not been systematically characterized.

Glucose and nitrogen availability affect longevity (Kapahi et al. 2017), and auxotrophy can influence utilization of glucose and other nutrients (Brauer et al. 2008; Gomes et al. 2007). Ammonium sulfate is a nitrogen source in yeast media that has been characterized as toxic for both proliferation and stationary phase survival (Hess et al. 2006; Santos et al. 2013; Santos et al. 2012). Preferred nitrogen sources, such as ammonium and glutamine, activate target of rapamycin (TOR) complex 1 (TORC1) activity (Stracka et al. 2014), a lifespan modulator across all eukaryotic model organism species (Harrison et al. 2009; Powers et al. 2006; Wanke et al. 2008; Wei et al. 2008). TORC1 promotes ribosome biogenesis, protein synthesis, and cell cycle progression (Barbet et al. 1996; Loewith and Hall 2011; Nakashima et al. 2008). Conversely, nutrient depletion inhibits TORC1, inducing cell cycle arrest, activation of stress response pathways, and quiescence. Genetic loss of ribosomal protein S6 kinase, *SCH9*, a downstream signal of active TORC1 (Urban et al. 2007) mimics the effects of TORC1 inhibition (Fabrizio et al. 2001). Thus, it appears that toxic levels of ammonium sulfate impede quiescence by driving cell cycle progression when other nutrients required to complete cell division are depleted, whereby inhibition of TOR signaling alleviates this dysfunction.

Replication stress is another modulator of lifespan (Weinberger et al. 2007; Weinberger et al. 2013), where it has been shown that genetic or nutrient interventions that promote rate-limiting substrates for DNA synthesis, allow for completion of “the final S-phase” that is needed for development of quiescence (Hartwell 1974). *SCH9* deletion or reduced glucose in the media alleviates disruption of quiescence by replication stress (Weinberger et al. 2013). Induction of quiescence by replication stress (hormesis) has also been reported (Ross and Maxwell 2018).

To promote systems level analysis of yeast quiescence, we sought to characterize auxotrophy and media composition influences broadly, in order to build a better foundation for reproducible studies by helping explain whether the weak correlation between previous genome-wide studies of CLS in the YKO libraries might be attributable to the use of different media, auxotrophic backgrounds, and/or phenotypic assays (Smith et al. 2016). The studies described here used quantitative high throughput cell array phenotyping (**Q-HTCP**), a robotic method for obtaining tens of thousands of growth curves per assay, and human-like media (HL), which was designed to eliminate ammonium toxicity and with greater resemblance to human tissue culture media (Hartman IV et al. 2015). Using this approach in highly controlled experiments, all possible auxotrophic allele combinations, availability of corresponding nutrients, glucose, or ammonium sulfate, as well as perturbation of TORC1 signaling or replication stress were examined for effects on quiescence development and maintenance by frequent monitoring of colony forming capacity extending past 60 days to characterize different stages of quiescence development. Thus, quiescence profiles rigorously characterize primary and interacting effects of perturbations with many replicates for each culture condition.

Results from this studies were useful for integrating and clarifying prior CLS literature and suggested strategies that could improve the consistency of interpretation moving forward. For example, TORC1 signaling appeared to affect CLS only in the context of ammonium sulfate addition to HL media. Reduction of CLS, associated with leucine auxotrophy and restriction, was enhanced by lysine auxotrophy and alleviated by methionine auxotrophy. The influence of replication stress on CLS depended on the intensity of the perturbation and also interacted with *MET17/met17* allele status and available concentrations of methionine and cysteine in the media. Increased glucose concentration reduced quiescence and CLS for most auxotrophic cultures, but did not, by itself (in the absence of ammonium sulfate), have much effect on fully prototrophic cultures. Ammonium sulfate enhanced loss of quiescence with increasing glucose in the prototroph, which was further modulated by auxotrophic alleles. For example, ammonium sulfate strongly induced loss of quiescence at high glucose concentration in the BY4741 background, but had less effect on the BY4742 strain, which was more sensitive to increased glucose than BY4741 in the absence of ammonium sulfate. Auxotrophic alleles and aeration were found to alter media acidification in complex ways that were not simply correlated with quiescence, suggesting auxotrophy modulates media acidification and quiescence somewhat independently.

In characterizing individual and combinations of auxotrophy and nutrient perturbations affecting quiescence in the genetic background of the YKO library, we conclude that these influences are numerous enough to confound prior studies, leading to weak correlation. YKO library mutations could have differential influence through interaction with factors that vary between studies, but haven’t been appreciated for their influence. A prototrophic reference strain, together with media optimized for quiescence that omits ammonium sulfate, could increase consensus from large-scale genetic and/or metabolic studies of yeast quiescence.

## Materials and methods

### Yeast strains and media conditions

BY4741 (*MAT**a** his3*Δ*1 leu2*Δ*0 met17*Δ*0 ura3*Δ*0*) and BY4742 (*MATαhis3*Δ*1 leu2*Δ*0 lys2*Δ*0 ura3*Δ*0*) were obtained from Research Genetics (Huntsville, AL). Additional ‘BY’ strains from the Boeke laboratory were shared by Jeff Smith (**Online Resource 2-Table S1**). Construction of additional auxotrophic strains by tetrad dissection of crosses between FY4 and BY4741/ BY4742 is described in **Online Resource 1**, and the strains are listed in **Online Resource 2-Table S2**. The BY X FY4 diploids that were dissected were heterozygous for all auxotrophic loci (*his3*Δ*1/HIS3 leu2*Δ*0/LEU2 lys2*Δ*0/LYS2 met17*Δ*0/MET17 ura3*Δ*0/URA3*) used in the YKO libraries. Results from tetrad analysis revealed reduced spore viability, which increased in a second backcross **Online Resource 2-Table S3**. Human-like (HL) yeast media was used for all experiments (Hartman IV et al. 2015); its composition is detailed in **Online Resource 2-Table S4**. When supplemented, the ammonium sulfate concentration was 0.5 g/L. The YNB for HL media was purchased from Sunrise Science (“Hartman Custom YNB”, https://sunrisescience.com/). Quiescence profiling assays were performed in 384 well plates from Evergreen Scientific (222-8210-01I) at 30°C, inverted to aerate the cells, and covered with gas permeable membranes (USA Scientific 9123-6100) after the first assay time (3-4 days after inoculation) to prevent contamination and evaporation. In 384-well format, the surface tension between the media and well hold it in place when the plate is inverted, so that cells settle at the air-liquid interface. For experiments involving pH measurement by media transfer, high-profile, square-well 384-well plates were used (USA Scientific #1843-8410). Additional Q-HTCP procedures are described elsewhere (Rodgers et al. 2014). Culture age is reported from time of inoculation. A doxycycline-repressible *TOR1* allele was constructed in the Y15578-1.2b strain (*MATα his3Δ1 leu2Δ0 ura3Δ0 can1Δ0::PGAL1-TADH1-PMFA1-his5+ lyp1Δ0 hmrΔ0::URA3ca*), as previously described (Hartman IV 2007; Singh et al. 2009).

### Quantitative high throughput cell array phenotyping (Q-HTCP) and data analysis

The Q-HTCP method was used to obtain cell proliferation parameters (CPPs) from growth curve analysis of yeast cultures spotted on to agar media (Hartman IV and Tippery 2004; Louie et al. 2012; Rodgers et al. 2014; Santos and Hartman IV 2019; Shah et al. 2007), which were used to characterize colony forming capacity and thus construct quiescence profiles. A custom imaging robot (Hartman laboratory) integrated with Cytomat 6001 (Thermo Fisher Scientific, Asheville, NC, USA) was used for agar cell array imaging at the time of transfer from liquid aging arrays to fresh HLD (2% dextrose) agar media, and ~every two hours until carrying capacity was surpassed. Image analysis and growth curve fitting was performed to obtain cell proliferation parameters (CPPs), using custom Matlab programs (Rodgers et al. 2014) to fit time series image data to the equation, G(t) = K/(1 + e^-r(t-l)^), assuming G(0) < K, where G(t) is the image intensity of a spotted culture vs. time, *K* is the carrying capacity, *r* is the maximum specific growth rate, and *L* is the time at which maximum absolute growth rate occurs, corresponding to G(t) = K/2 (the time to reach half carry capacity) (Shah et al. 2007). A MediaClave (Integra Bioscience) was used for consistent media preparation. A Masterflex L/S peristaltic pump (Cole Parmer) was used for consistent dispensing of agar media in imaging-type monowell plates (Nunc #267060). A Caliper Sciclone ALH 3000 or Biotek MultiFlo FX instrument was used for liquid dispensing into microwell plates.

### Quiescence profiles

Quiescence profiles report on establishment and maintenance of quiescence, in terms of colony-forming capacity during stationary phase, and are plotted as *L* (hours to half carrying capacity) *vs. age* (days) in the main figures. Note: *r* is the intrinsic exponential growth rate and is related to the doubling time (DT) during exponential growth by DT = ln2/*r*. Thus, change in L is convertible to change in viability (assuming K and r remain constant) by 0.5^(*Ln-L0* /DT)^ (100%). However, K and/or r can change with age, thus results are reported in terms of raw CPPs for clarity and accuracy. Qualitative differences in colony forming capacity are also evident in quiescence profiles for the cell proliferation parameter, K (carrying capacity), which complement L profiles and are assembled in **Online Resource 3**. Results are reported from the time of inoculation (i.e., ‘Day 0’). In summary, quiescence profiles consist of Q-HTCP outgrowth data vs. age. L is the primary estimate of CFU capacity (**Figs. 1, 2, and S1**), where increase in L indicates loss of quiescence. Quiescence can be further informed by CPP profiles for *K* or *r*. At low colony forming capacity, when individual colonies are observed instead of a lawn, r and K parameters are depressed, because G(t) is the average pixel intensity across the entire lawn of cells. For cultures where no cell proliferation is detected, the maximum L value is set to 50, corresponding to *K* = 0.

**Fig. 1.**
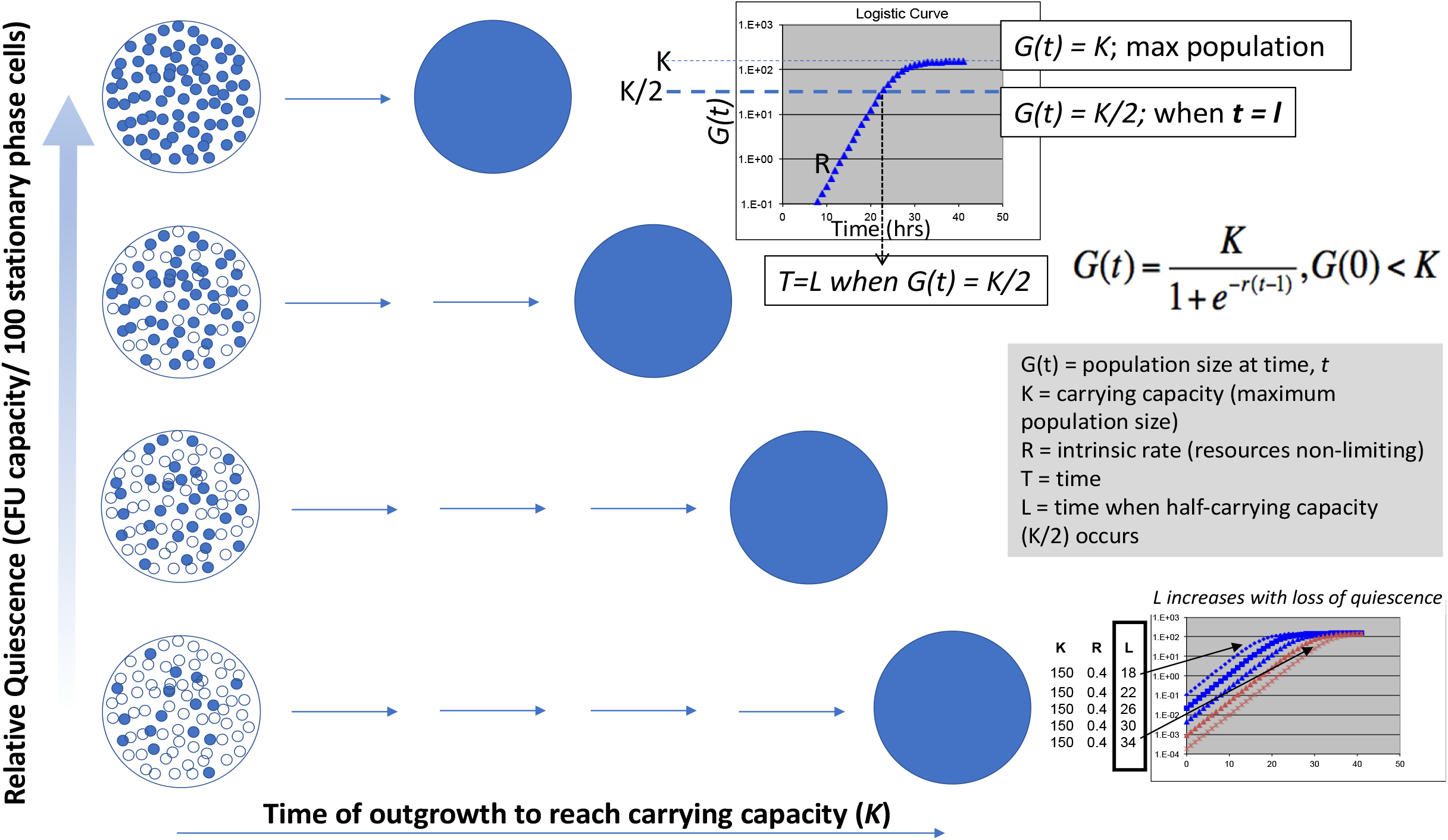
The cell proliferation parameter, L, reports on colony forming capacity and quiescence. Quiescence is the maintenance of colony forming capacity with age. As colony forming capacity is reduced by a lower percentage of cells able to reenter the cell division cycle, a spot culture must proliferate for additional generation times to reach the final carrying capacity (K). Q-HTCP captures time series image data reporting on colony forming capacity of stationary phase cultures upon transfer of a fixed volume to fresh agar media. Fitting the quantitative image data to the logistic function, *G(t) = K/(1 + e^-r(t-l)^)*, yields the parameter ‘L’, which is the time required for the culture to reach half of carrying capacity (‘K/2’), thus reporting on colony forming capacity. L increases with loss of quiescence. *For more detail, see Online Resource 4 – Fig. S1*.

**Fig. 2.**
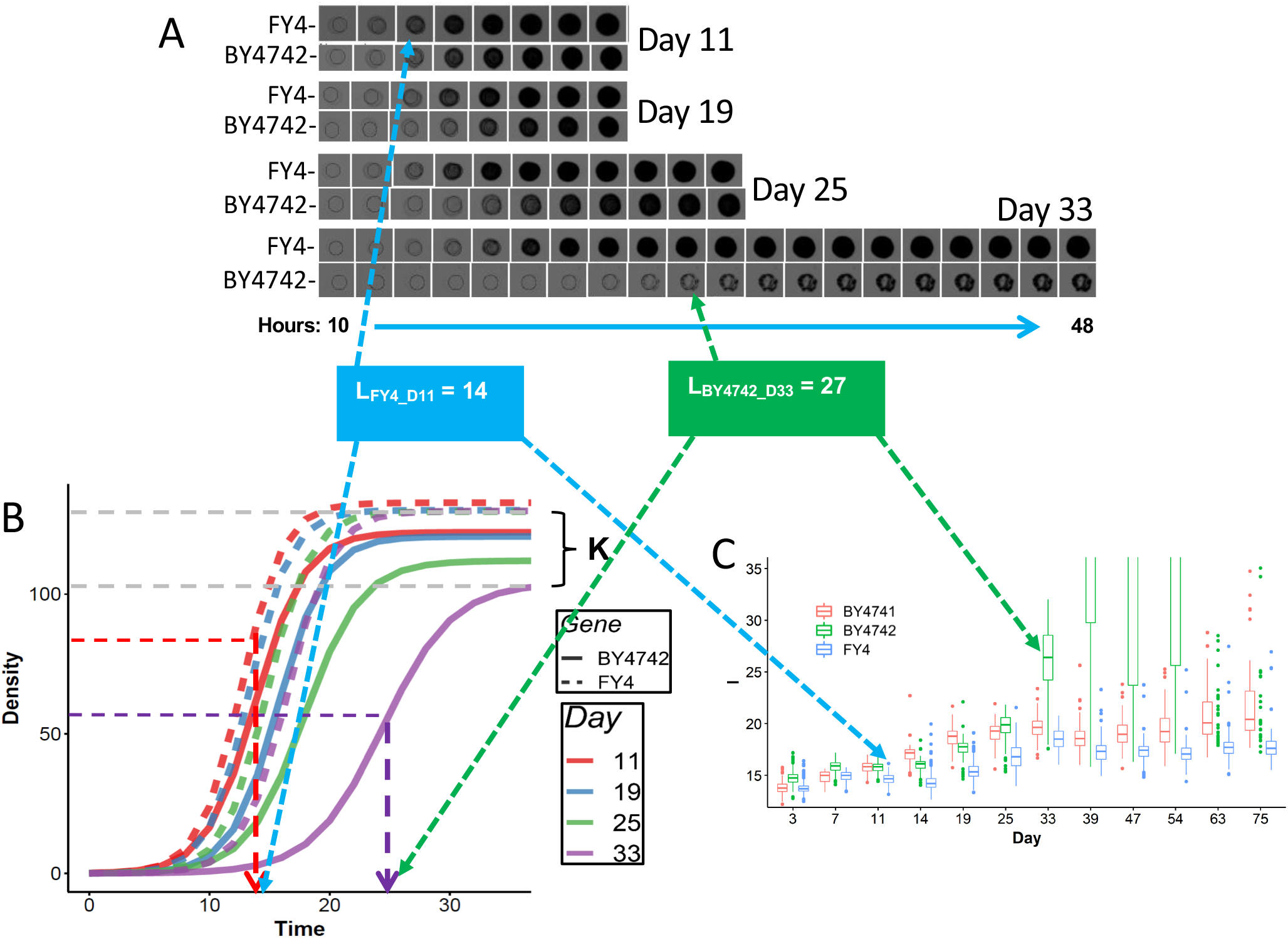
Examples of quiescence profiles. (**A**) Q-HTCP image data are quantified by image analysis and (**B**) fit to a growth curve model, from which the parameter, *‘L’* (blue and green boxes), is determined (*see Fig. 1*) and **(C)** plotted for replicate cultures (N=96 in this example), providing profiles of mean quiescence and variance with age. Arrows indicate (**A**) individual images and (**B**) growth curves representative of (**C**) distributions of replicate cultures indicating relatively high quiescence (FY4 at Day 11, blue) or low quiescence (BY4742 at Day 33, green). BY4741 (not shown in panels A and B due to space limitation) exhibits a quiescence profile distinct from FY4 and BY4742. All cultures were aged in HLD media with 2% glucose and without ammonium sulfate. Strain genotypes are listed in **Online Resource 2 - Table S1**.

### High throughput measurement of media pH

A 384-well format pH assay was developed with bromophenol blue (**BPB**) as a dye indicator. A standard curve was established using conditioned media from stationary phase yeast cultures exhibiting a range of pH between 3 and 5.4, which were obtained from BY4741 cultures grown in flask cultures in HLD media with a range of dextrose (0.1 – 1.0%) or HU (0-120 mM) concentrations, collected at different culture ages, and measured by pH electrode. After centrifugation to clear cells, 10uL of the stationary phase conditioned media was transferred from 384 culture aging arrays (in parallel with standards of known pH) to wells of a plate pre-loaded with 30 uL BPB reagent (stock concentration of 1 mg/mL BPB stock in 50% ethanol is diluted 1:20 in water on the day of use) and read at 594 nM with a Biotek Powerwave HT plate spectrophotometer. pH standard curves were generated using a linear regression model. The pH of each culture was inferred from its absorbance plotted against the standard curve (see **Online Resource 1** for additional details).

### Data processing and visualization

R (http://www.r-project.org/) and the ggplot2 package (Wickham 2016) was used for processing and visualizing Q-HTCP data. In box plots, the median is represented by the center bar, the 25^th^ and 75^th^ percentile by the lower and upper hinges, and the lesser of either 1.5X interquartile range or extreme value by the whiskers. Density plots of pH data were generated using the geom_density() function of ggplot2 to create kernel density estimates (smoothed histograms).

## Results

### Quiescence profiles obtained by Q-HTCP

Quiescence is inferred by sustained colony forming capacity with age, and is measured by outgrowth from stationary phase cultures (Murakami et al. 2008). Q-HTCP, which has capability to obtain tens of thousands of high-resolution growth curves simultaneously (Louie et al. 2012; Santos and Hartman IV 2019; Santos et al. 2019), was applied for high throughput analysis of colony forming capacity. Estimation of colony forming capacity is illustrated in **Figure 1**. Assuming constant stationary phase cell density, culture droplets contain approximately the same number of cells so that viability within the population is inversely proportional to the time required for the droplet to reach half of carrying capacity, a cell proliferation parameter referred to as ‘L’. Thus, loss of quiescence is detected by increases in L with age (**Fig. 1**).

The increased throughput of Q-HTCP derives from its basis in imaging and image analysis of cultures spotted onto agar (Hartman IV and Tippery 2004). Differential quiescence can be directly visualized from raw images of individual cultures (**Fig. 2A**), but fitting image analysis results to a logistic growth function (**Fig. 2B**) enables extraction of cell proliferation parameters (Shah et al. 2007) and automated analysis. The resolution of quiescence profiles derives from how frequently the assay is deployed with age, the frequency with which colony forming capacity is assessed during each outgrowth trial, and how variance (*i.e*., the quantity of replicate cultures) is measured. Each aspect contributes accuracy to discriminate differential quiescence between culture populations (**Fig. 2C**). For example, from comparison of raw data from representative cultures (1 of 96 replicates) for the prototrophic parent of the yeast gene knockout (**YKO**) library (FY4) vs. the auxotrophic genetic background common to the *MATa* YKO library (BY4742, *MATa his3*Δ*1 leu2*Δ*0 lys2*Δ*0 ura3*Δ*0*), one can appreciate, qualitatively, a loss of quiescence between Days 11 and 33 from either the images (**Fig. 2A**), or growth curves derived from them (**Fig. 2B**). However, quiescence profiles presented as box plots can summarize much more data and convey quantitative information about differences in phenotypic distributions (e.g., **Fig 2C** represents 3456 growth curves; 3 profiles/ experiment X 12 timepoints/ profile X 96 growth curves/ timepoint). Quiescence profiles for BY4741, BY4742 and FY4 reveal that FY4 establishes and maintains quiescence most efficiently, while BY4741 and BY4742 each differ in their colony forming capacity profile with age (**Fig. 2**). Box plots for the cell proliferation parameters (K), corresponding to main L, which is presented in the figures, are provided in **Online Resource 3.** Additional detail about the use of L to estimate colony forming capacity and quiescence (**Online Resource 4-Fig. S1A**), an example of whole Q-HTCP array images (**Online Resource 4-Fig. S1B**), and comparison of the L-based estimate of colony forming capacity with traditional serial dilution spot tests (**Online Resource 4-Fig. S1C**) are presented in **Online Resource 4**.

### Interactions between auxotrophy and glucose or nitrogen availability modulate quiescence

Dietary restriction is a hallmark of longevity and/or healthspan across many eukaryotes (Kapahi et al. 2017; Mair and Dillin 2008). As such, yeast quiescence (induced by nutrient limitation) could potentially serve as a biologically informative systems level model. Previous work has demonstrated that yeast quiescence responses depend upon distinct nutrient limitations (Boer et al. 2008; Gresham et al. 2011; Lillie and Pringle 1980; Petti et al. 2011). Others have noted that the ammonium sulfate present in standard media is toxic to yeast and reduces chronological lifespan (Hess et al. 2006; Santos et al. 2013). Media recipes and/or autoclaving amino acids can have additional effects (Alvers et al. 2009; Smith et al. 2016). Glucose concentration has been used as a paradigm of caloric restriction by decreasing the initial standard glucose concentration of 2% to 0.5%, which shifts metabolism from glycolysis to respiration during logarithmic phase growth and reduces media acidification. However, the relative importance and context-dependence of different glucose restriction mechanisms remain to be further clarified (Arlia-Ciommo et al. 2018a; Arlia-Ciommo et al. 2018b; Choi et al. 2011; Fabrizio and Wei 2011; Maruyama et al. 2016; Mirisola and Longo 2012; Murakami et al. 2011; Weinberger et al. 2013; Wierman et al. 2017).

We developed HL media to eliminate the effect of ammonium toxicity (Hess et al. 2006; Santos et al. 2013; Santos et al. 2012) and improve translational relevance between yeast and human cells by increasing similarity between yeast and human cell culture media (Hartman IV et al. 2015). Influences of glucose, ammonium sulfate, and selected auxotrophic backgrounds on quiescence in HL media were evidenced by quiescence profiles for BY4741, BY4742, FY4 (prototrophic parent) and BY4712 (*MAT**a** leu2*Δ*0*), using dextrose concentrations of 0.4, 2, and 5%, each with and without 0.5 gm ammonium sulfate (**Fig. 3**). Increasing glucose resulted in loss of quiescence beginning between Days 7 and 11, with BY4741 and BY4742 being more sensitive to this effect than the prototroph (FY4) or the *leu2*Δ*0* single auxotroph (BY4712) (**Fig. 3**, compare Days 11 and 14 across panels). BY4741 lost quiescence (between Days 7 and 14) the most rapidly at 5% glucose, but also had a gasping-like (Fabrizio et al. 2004) response (recovery of increased CFU capacity between Days 14 and 51). BY4742 did not achieve the same initial colony forming capacity (Days 4 and 7), and was less impacted than other genotypes by ammonium sulfate. In the absence of ammonium sulfate, BY4712 resembled FY4, until Day 28, after which it exhibited relative loss of quiescence in 2% and 5% glucose. Ammonium sulfate addition reduced quiescence, most notably at higher glucose and in the FY4 strain. In the absence (but not in the presence) of ammonium sulfate, the quiescence profile of the prototroph was relatively unaffected by increasing glucose (**Fig. 3**). Additionally, low glucose concentration alleviated much of the differential quiescence observed due to auxotrophy and ammonium sulfate at the higher glucose concentrations (**Fig. 3A**).

**Fig. 3.**
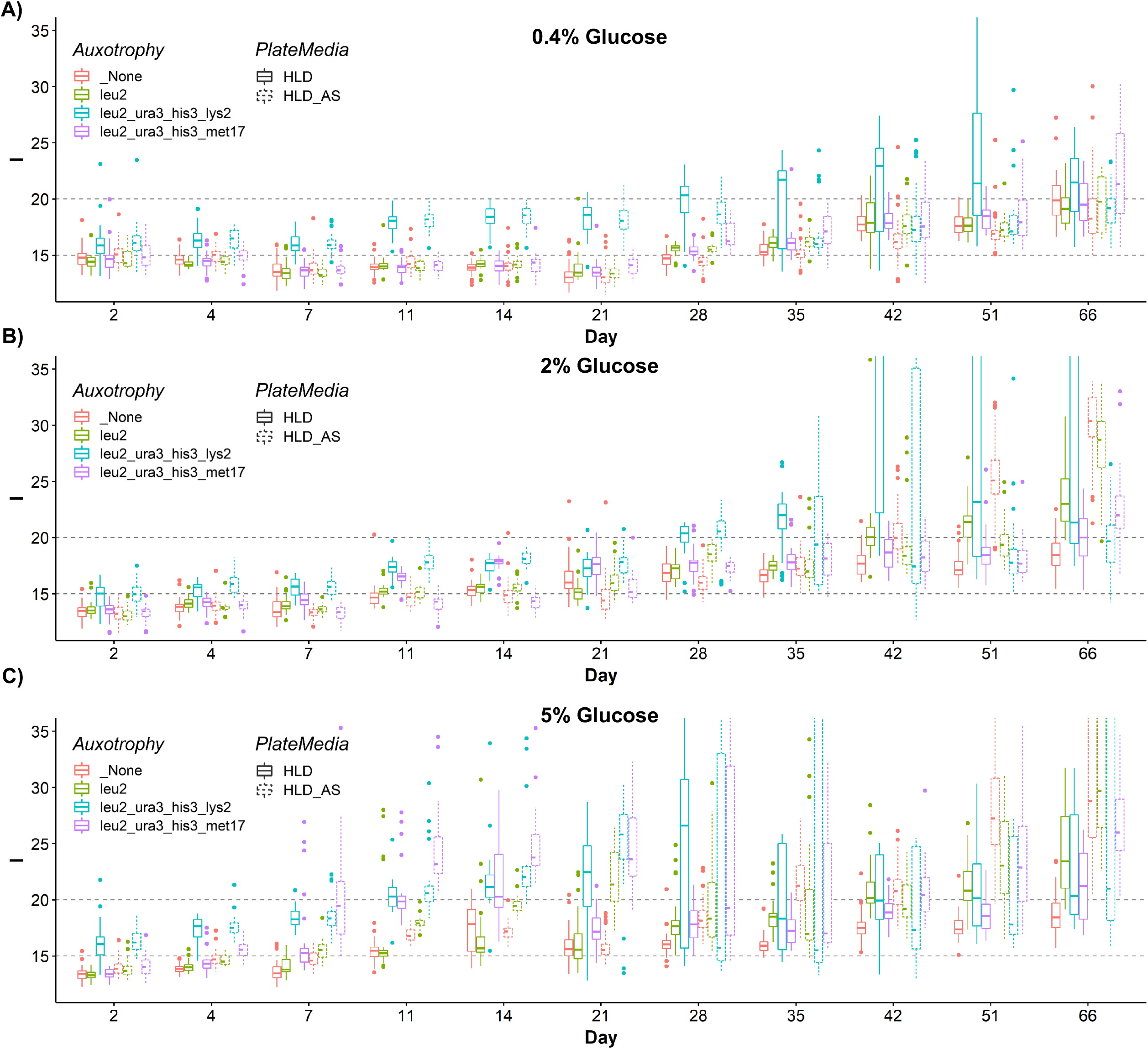
Concentrations of glucose and ammonium sulfate have interacting influences on quiescence in HLD media. Quiescence profiles for FY4 (prototroph) BY4741 (*MAT**a** his3 leu2 met17 ura3*), BY4742 (*MATa his3 leu2 lys2 ura3*), and BY4712 (*leu2* single auxotroph) are compared in aerated HLD media for 3 glucose concentrations: (**A**) 0.4%, (**B**) 2.0%, and (**C**) 5%, each with or without .5 g/L ammonium sulfate.

To summarize above, quiescence profiles of BY4741 and BY4742 (which share *his3, leu2*, and *ura3* auxotrophy, but differ by *met17* vs. *lys2* auxotrophy), FY4 (prototrophic) and BY4712 (*leu2* auxotrophy only) showed that relative viability can depend on the age at which cultures are compared, the glucose concentration, and/or inclusion of ammonium sulfate in the media (**Fig. 3**). Additionally, BY4730 (*MAT**a** leu2*Δ*0 ura3*Δ*0 met17*Δ*0*) had a similar CLS profile to BY4741 (*leu2*Δ*0 ura3*Δ*0 met17*Δ*0 his3*Δ*1*), suggesting the *his3*Δ*1* allele did not influence quiescence in this context (**Online Resource 4-Fig. S2**). BY4700 (*MAT**a** ura3*Δ*0*) and BY4706 (*MAT**a** met17*Δ*0*) exhibited quiescence profiles similar to FY4 until Day 42, at which time *met17*Δ*0* conferred prolonged quiescence relative to *MET17*, but only in HL media with 2% or 5% glucose with ammonium sulfate added (**Online Resource 4-Fig. S3**). Thus, in HL media, glucose and ammonium sulfate availability, auxotrophic alleles of *leu2*Δ*0, lys2*Δ*0*, and *met17*Δ*0*, and combinations of these modulators influence quiescence in specific developmental timeframes. In contrast, *ura3*Δ*0* and *his3*Δ*0* have relatively minor influences. Quiescence profiling was next applied to investigate influences of auxotrophic nutrient availability.

### Nutrient availability reveals additional influences of auxotrophic mutations

Auxotrophic nutrient limitation, and leucine deficiency in particular, reduces viability in stationary phase (Aris et al. 2013; Boer et al. 2008; Gomes et al. 2007; Santos et al. 2013), and thus it was characterized under conditions of this study. Auxotrophic nutrients were varied at 1/3- or 3-times recipe amount in the HLD media with 2% glucose (**Fig. 4**). Leucine availability had differential effects in *leu2*Δ*0* auxotrophs, depending on the context of other auxotrophic alleles. 1/3X leucine availability had the strongest phenotypic effect on BY4741, which underwent rapid loss of quiescence between Days 3 and 11. However, the rapid loss in BY4741 colony forming capacity was reversed in a gasping-like response from Days 11 to 19, followed by recurring loss of CFU capacity after Day 19 (**Fig. 4A**). BY4730, which is isogenic to BY4741 except that it is prototrophic for *HIS3*, exhibited the same profile, indicating his3/*HIS3* allele status has little influence (**Online Resource 4-Fig. S4**). In contrast to the BY4741 phenotype, the *leu2* single auxotroph (BY4712) lost quiescence in response to leucine restriction more gradually between Days 14 and 25, after which the cultures died without gasping (**Fig. 4**), suggesting interaction between *leu2* and other auxotrophic loci.

**Fig. 4.**
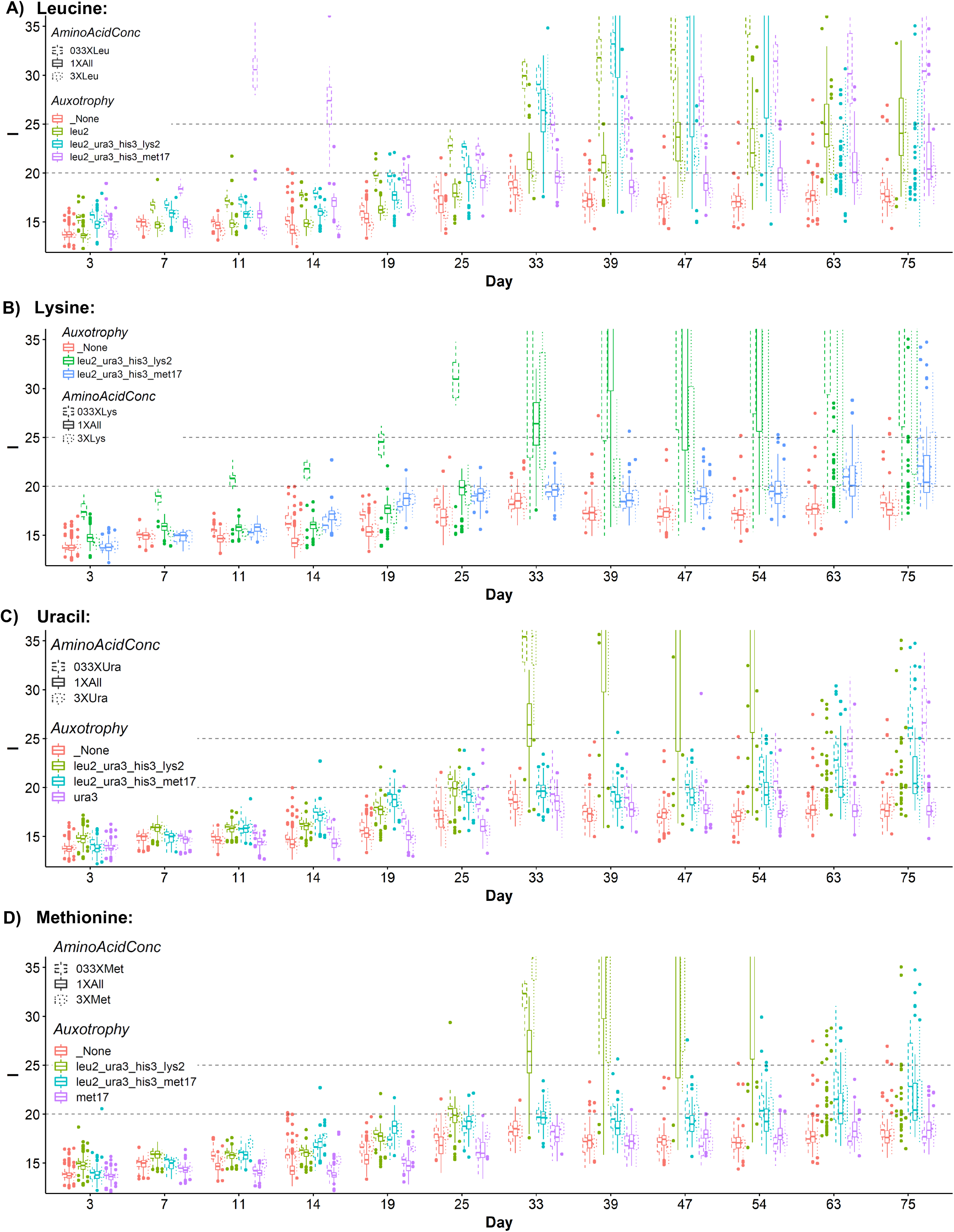
Auxotrophic nutrient limitations variably influene quiescence. In each panel, FY4 (prototroph; orange), BY4742, and BY4741 were aged in parallel, using media with 1/3X, 1X, or 3X of (**A**) leucine, (**B**) lysine, (**C**) uracil, or (**D**) methionine. (**A-D**) FY4 loses some quiescence between days 14 and 39, but maintains a largely similar profile under most conditions tested. Single auxotrophs relevant to each amino acid were included (if available), including (**A**) BY4712 (*leu2*), (**C**) BY4700 (*ura3*), and (**D**) BY4706 (*met17)*. (**A**) BY4712 (*leu2* single mutant) loses quiescence relative to FY4 and BY4741, but preserves quiescence relative to BY4742. Like BY4741 and BY4742, BY4712 loss of quiescence is exacerbated by 1/3X leucine, but relatively unaffected by 3X leucine. BY4741 (met17 autotroph) rapidly loses quiescence with 1/3X leucine in media, but recovers quiescence (gasping), evidenced by greater colony forming capacity after Day 33. (**B**) BY4742 (harboring *lys2*) and FY4, to a lesser degree, but not BY4741 (harboring *met17*), lose quiescence with 1/3X lysine. BY4742 loses quiescence beginning at Day 33, which is partially mitigated by 3X lysine. (**C**) Loss of quiescence is evidenced after Day 47 with 1/3X uracil for BY4700 (*ura3* single auxotroph) and BY4741, but not FY4 or BY4742. (**D**) 3X methionine causes a transient (Days 11-25) loss of quiescence for BY4706, but BY4706 exhibits more stable quiescence in the presence of ammonium sulfate (**Online Resource 4-Fig. S3**).

Initial colony forming capacity of BY4742 in HLD media was slightly reduced relative to FY4, BY4741 or BY4712 (**Fig. 4**), and quiescence was more stable than BY4741 against perturbation with 1/3X leucine between Days 7 and 14. BY4742 also had a gasping response to 3X leucine after Day 33 (**Fig. 4A**). 1/3X leucine limitation impeded quiescence slightly in the FY4 prototroph between Days 15-25, but not the *ura3*Δ*0* or *met17*Δ*0* single mutant strains (**Online Resource 4-Fig. S5A**), indicating effects of leucine independent of leucine auxotrophy. Between Days 25-54, 3X leucine had little effect relative to 1X leucine on quiescence of FY4, but slightly promoted quiescence in the *met17* or *ura3* auxotrophic context (**Online Resource 4-Fig. S5A**). Taken together, these results suggest that *leu2* auxotrophy and leucine availability interact with *met17*Δ*0* and *lys2*Δ*0* auxotrophy in altering quiescence profiles. The quiescence profile of the FY4 prototroph was only slightly affected by depletion of leucine and unaffected by its supplementation in HLD media with 2% glucose (**Fig. 4A**).

Perturbation of lysine concentrations in HL media primarily affected BY4742 (the *lys2* auxotroph), as 1/3X lysine reduced the initial colony forming capacity evident at Day 3 and also accelerated quiescence between Days 11 and 33 (after which BY4742 cultures lose essentially all quiescence, regardless of lysine availability) (**Fig. 4B**). 1/3X lysine was associated with a slightly accelerated loss of quiescence between Days 11 and 33 in the prototrophic FY4 context, BY4700 (*ura3*Δ*0*), and BY4706 (*met17*Δ*0*), which was not observed for BY4712 (*leu2*Δ*0*) or BY4741 (**Online Resource 4-Fig. S5B**). 3X lysine effects were few, but included transient gasping of BY4742 at Day 39 (**Fig. 4B**) and loss of quiescence in BY4712 (*leu2*Δ*0*), also at Day 39 (**Online Resource 4-Fig. S5B**).

1/3X uracil gradually reduced quiescence in the *ura3*Δ*0* single auxotroph (BY4700) between Day 11 and Day 47, but not for BY4741. However, quiescence was similarly compromised by 1/3X uracil in both strains after Day 47 (**Fig. 4C**). Effects of 1/3X uracil were not seen for BY4742, which lost quiescence before Day 47. Uracil availability did not impact quiescence in FY4 or the *met17* (BY4706) or *leu2* (BY4712) single auxotrophs (**Online Resource 4-Fig. S5C**).

Methionine restriction, an anti-aging paradigm across eukaryotic species (Lee et al. 2016), had only slight effects in the context of HLD media. The strongest effect of methionine in HL media was 3X methionine supplementation, which accelerated quiescence of BY4706 (*met17*Δ*0*) between Days 11 and 33 (**Fig. 4D**). Either 1/3X or 3X methionine perturbation was associated with small reductions in quiescence in FY4 or BY4700 (*ura3*Δ*0*) (**Online Resource 4-Fig. S5D**). Other effects were subtle. A hypothetical explanation for the influence of methionine limitation in some but not other contexts is that restriction can be beneficial under suboptimal nutrient conditions, but that HL media supports quiescence in a way that somewhat precludes such benefits. Consistent with this idea, BY4706 exhibited stable quiescence relative to FY4, beginning around Day 42 of age, when challenged with ammonium sulfate in HL media with 2% dextrose (**Online Resource 4-Fig. S3B**).

The results above indicate that auxotrophic mutations and nutrient factors exert dynamic (*e.g*., non-linear and/or bi-directional) influence across different stages of quiescence establishment and maintenance, evidenced by age-specific differential quiescence; the stages roughly consisting of: (1) initial colony forming capacity in early stationary phase, up to ~Day 10-14 (quiescence formation), (2) between ten days and a month (transition to long term quiescence), and (3) after 4-5 weeks (maintenance of stable quiescence).

### Systematic analysis of auxotrophic loci and their combinatorial effects on quiescence profiles

The studies here were focused on auxotrophic combinations present in the YKO libraries (*i.e*., the BY4741 and BY4742 backgrounds), comparing selected parental strains from which they were derived (Brachmann et al. 1998) (**Online Resource 2, Table S1**), and thus revealing potential sources of non-concordance between previously published genome-wide CLS studies (Smith et al. 2016). As noted above, the findings are generally consistent with prior literature about auxotrophy and media composition. To further characterize how these auxotrophic alleles differentially influence CLS or quiescence studies, 384 progeny comprising an assortment of all 32 auxotrophic allele combinations were obtained by tetrad dissection of compound heterozygous FY4/BY diploids (*HIS3/his3*Δ*1, LEU2/leu2*Δ*0, LYS2/lys2*Δ*0, MET17/met17*Δ*0, URA3/ura3*Δ*0*). The resulting strains (**Online Resource 2, Tables S2A-S2B**) were profiled for quiescence, varying culture aeration, and correlating quiescence with media acidification.

Quiescence profiles for the panel of 384 auxotrophic strains were obtained in triplicate in HLD with 2% glucose. Single loci were assessed by comparing all cultures having an individual prototrophic vs. auxotrophic allele (**Figs. 5A-E**). Consistent with results obtained from profiling the BY strains, *LYS2* and *LEU2* prototrophy, along with *met17* auxotrophy tended to support quiescence establishment and/or maintenance, while *URA3/ura3* and *HIS3/his3* allele status had little influence on quiescence profiles (**Figs. 5A-E**). The *URA3* and *HIS3* loci have been shown to affect survival in stationary phase in other experimental contexts (Boer et al. 2008).

**Fig. 5.**
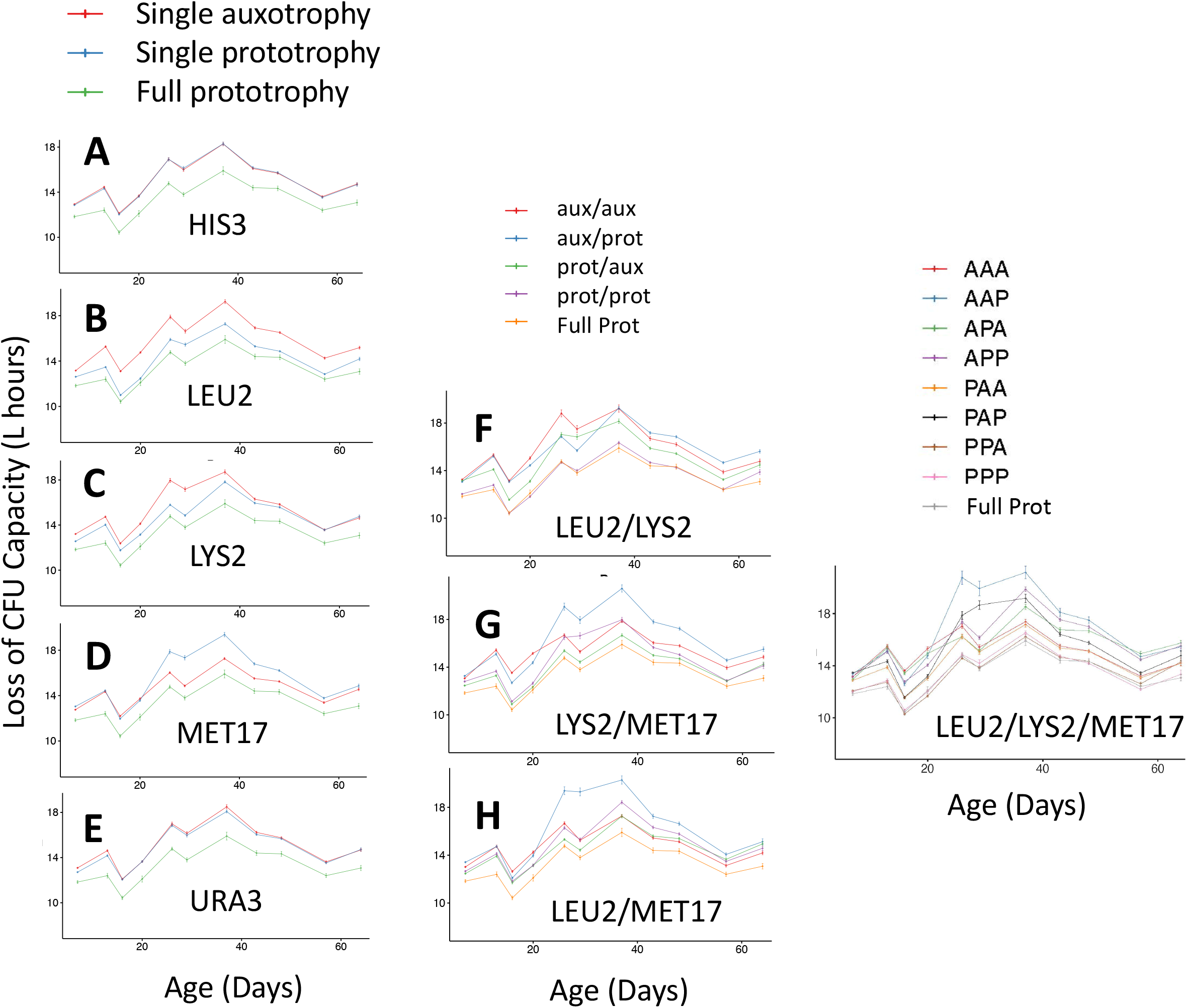
Combinatorial influences of *LEU2, LYS2*, and *MET17* allele status on quiescence. All possible combinations of prototrophy / auxotrophy for *HIS3/his3, LEU2/leu2, LYS2/lys2, MET17/met17*, and *URA3/ura3* were obtained by tetrad dissection of compound heterozygous diploid yeast containing all FY4 and BY alleles. 384 haploid progeny were studied in replicate (**Online Resource 2-Table S2A**). Progeny were grouped by indicated loci for comparisons. (**A-I**) Quiescence profile of the full prototroph is plotted on each graph for reference. (**A-E**) Across all genotypes, comparison of auxotrophy (red) vs. prototrophy (blue) reveals that (**B**) *leu2* or (**C**) *lys2* auxotrophy, and (**D**) *MET17* prototrophy reduce quiescence, while (**A**) *HIS3/his3* or (**E**) *URA3/ura3* allele status do not strongly influence quiescence in HL media. (**F-H**) Quiescence profiles for the four possible allele combinations of (**F**) *LEU2/leu2* and *LYS2/lys2*, (**G**) *LYS2/lys2* and *MET17/met17*, and (**H**) *LEU2/leu2* and *MET17/met17* reveal combinatorial influences on quiescence. (**I**) Three-locus analysis reveals the *leu2 lys2 MET17* (‘AAP’, blue) genotype to be the most compromised for quiescence, and much alleviated by *met17* auxotrophy (‘AAA’, red). In each panel legend the allele status is indicated by ‘A’ or ‘aux’ (auxotrophic) and ‘P’ or ‘prot’ (prototrophic), with the locus order alphabetic and indicated on the graph (*e.g*., for Panel F, locus order = *LEU2/LYS2*).

Pairwise combinations of the auxotrophic loci having single-allele effects revealed compounding of quiescence defects resulting from *lys2*Δ*0 MET17* (**Fig. 5F**), *leu2*Δ*0 MET17* (**Fig. 5G**), and *leu2*Δ*0 lys2*Δ*0* (**Fig. 5H**). In considering three-locus allele combinations, quiescence was most disrupted by the *leu2 lys2 MET17* genotypes, along with *leu2 LYS2 MET17, leu2 LYS2 met17*, and *LEU2 lys2 MET17*, while the *LEU2 LYS2* prototrophic strains, with either *MET17* or *met17* were comparable to the fully prototrophic strain (**Fig. 5I**). The *leu2 lys2 met17* genotype had lower initial CFU capacity, but established and maintained quiescence in sharp contrast to *leu2 lys2 MET17* strains (**Fig. 5I**), demonstrating that *met17*Δ*0* auxotrophy suppresses the quiescence defects of *leu2*Δ*0* and *lys2Δ0*.

### Interaction between auxotrophy and aeration status influences media acidification

Media acidification has been implicated in yeast CLS and other aging models (Fabrizio and Wei 2011; Maruyama et al. 2016; Mirisola and Longo 2012; Murakami et al. 2011). Glucose concentration has been used to model caloric restriction in yeast CLS, but also positively correlates with alcohol production and media acidification after the diauxic shift. Acetic acid has been associated with chronological aging (Burtner et al. 2009; Mirisola and Longo 2012), but mechanisms of chronological longevity independent of media acidification have been demonstrated (Arlia-Ciommo et al. 2018a; Cao et al. 2016; Kapahi et al. 2017; Laporte et al. 2018; Lefevre et al. 2015; Lillie and Pringle 1980; Quan et al. 2015; Shi et al. 2010; Wierman et al. 2017; Wierman et al. 2015). Considering that other forms of dietary restriction, besides glucose and calories, are of interest in yeast and animals (Lee et al. 2016; Mair and Dillin 2008), it would be helpful to further clarify the impact of media acidification relative to other aging mechanisms (Fabrizio and Wei 2011). We sought the correlation of media acidification with quiescence in the context of auxotrophy, but controlling for media influences, as an alternative means to test its influence (Gresham et al. 2011; Longo et al. 2012).

A pH indicator dye-based assay was developed to measure culture pH in parallel with colony forming capacity for the panel of strains harboring all auxotrophic allele combinations. Building on the observation that the *LEU2, LYS2*, and *MET17* loci influence quiescence (**Fig. 5**), the correlation between quiescence and media acidification for strains harboring different allele combinations at these loci, was ascertained in a single media composition (**Fig. 6**). pH was measured from stationary phase cultures at Day 31, and colony forming capacity was assessed at Day 42, when media acidification and quiescence were stable (data not shown). Respiration, and its role in producing organic acids from glycolytic substrates, was perturbed by differentially aerating the cultures (see methods) to further assess the potential correlations between auxotrophy, media acidification and quiescence (**Fig. 6**). Aeration resulted in greater media acidification, with stationary phase media pH ranging from 3.5 to 4.1 under non-aerated conditions (**Fig. 6A**) and from 3.0 to 3.5 under aerated conditions (**Fig. 6B**). Interestingly, however, auxotrophy exerted differential influences on media pH and quiescence under non-aerated (**Fig. 6C**) vs. aerated (**Fig. 6D**) conditions, indicative of interactions between auxotrophic loci and aeration status with regard to media acidification (**Fig. 6E**).

**Fig. 6.**
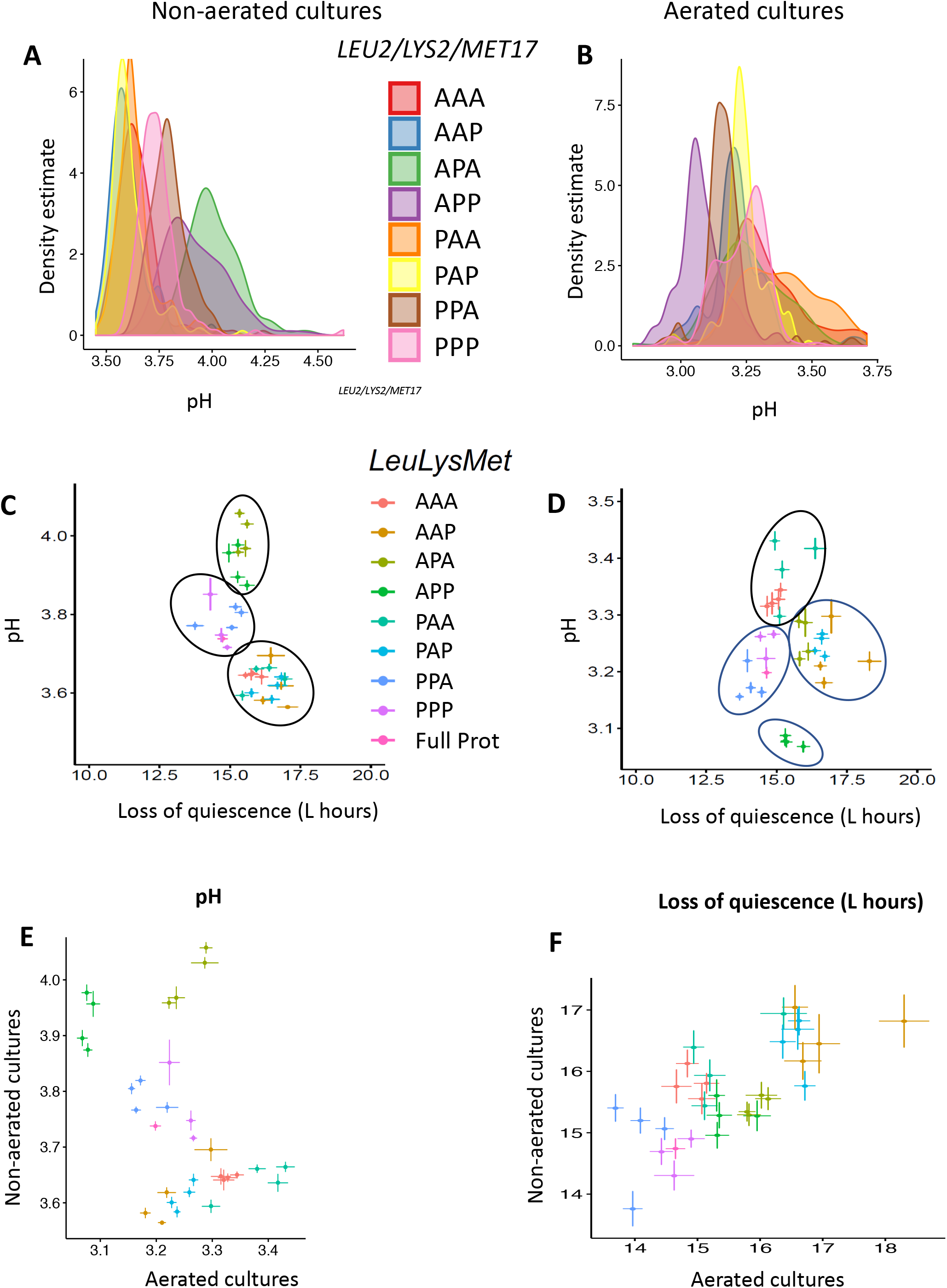
Auxotrophic effects on media acidification and quiescence are weakly correlated in HL media. Like Fig. 5I, the 3-locus genotype refers to the allele status (indicated by ‘A’ (auxotrophic) or ‘P’ (prototrophic) for the locus order, *LEU2/LYS2/MET17*. **(A-B)** Stationary phase pH at Day 31 from replicate strains with the designated genotypes. Density distributions are shown for (**A**) non-aerated and (**B**) aerated cultures. **(C-F)** Four data points are shown for each of the 3-locus genotypes (representing combinations of *his3/HIS3* and *ura3/URA3* in each background; all values and genotypes are presented in **Online Resource 2-Table S5**). **(C-D)** Quiescence (L) vs media acidification (pH) for all auxotrophic genotypes (mean and standard error are indicated) under **(C)** non-aerated and **(D)** aerated conditions. (**E-F**) Aeration interacts with auxotrophy to influence (**E**) media acidification, but less so for (**F**) quiescence.

### Independent effects of auxotrophy on acidification and quiescence in HL media

Correlations between media acidification (pH taken at Day 31) and quiescence (estimated by the mean of L at Day 42) were examined for all 32 auxotrophic combinations (**Figs. 6C-D**). Non-aerated cultures clustered into three groups with regard to pH, but were overlapping in their quiescence (**Fig. 6C**): *(1)* genotypes with the most acidified media included all of the *lys2* auxotrophs; *(2)* the cluster with intermediate media acidification included the *LEU2 LYS2* double prototrophs; and *(3)* the cultures with the highest conditioned media pH were the *leu2 LYS2* genotypes. Aerated cultures, by contrast, clustered into four groups, depending on the allele status at the *MET17* locus in addition to *LEU2* and *LYS2*. Clusters with the highest and lowest quiescence had intermediate pH, and the clusters with the highest and lowest pH had intermediate quiescence (**Fig. 6D**): *(1)* the *LEU2 LYS2* prototrophs were associated with the highest quiescence but intermediate media acidification, and with a slight trend within the group for *met17* auxotrophs to have both increased quiescence and greater acidification; (*2*) the *lys2 met17* double auxotrophs had the least acidified media, and intermediate quiescence; *(3)* both *lys2 MET17* genotypes displayed intermediate acidification and the lowest quiescence, while the *leu2 LYS2 met17* genotypes were clustered with them, maintaining slightly greater quiescence; *(4)* the *leu2 LYS2 MET17* genotypes clustered distinctly with intermediate quiescence and the most acidified media. Of note, influences of *HIS3/his3* and *URA3/ura3* allele status on quiescence (and media acidification) could be observed within selected auxotrophic contexts, such as *LEU2 lys2 met17* and*leu2 lys2 MET17*, by the spread of mean measures across the four genotypes within the respective groups (**Fig. 6D, Online Resource 2-Table S5**).

These results indicate that auxotrophy and combinations of auxotrophic loci can differentially affect media acidification and quiescence, which is further subject to aeration, a variable not always controlled between CLS assays. Multi-locus influences of auxotrophy on both media acidification and quiescence can occur without varying media composition, in the absence of ammonium sulfate, and with amino acid concentrations that fully complement the respective auxotrophies (**Fig. 4**). Thus, associations between media acidification and CLS (Burtner et al. 2009; Burtner et al. 2011) may depend on auxotrophic and other metabolic context, as they do not appear causally associated, using a prototrophic strain and aerated HL media as a reference (**Fig. 6D**). Interaction between auxotrophic loci and aeration tended to impact media acidification (**Fig. 6E**) more than quiescence (**Fig. 6F**). Evolutionarily conserved genetic influences on quiescence may be characterized in a more tractable manner using prototrophic yeast, with media optimized for quiescence, as the initial reference condition.

### Auxotrophy and media composition modulate influences of TORC1 and replication stress

TORC1 activity influences lifespan across many eukaryotic species (Loewith et al. 2002; Powers et al. 2006; Wanke et al. 2008; Wei et al. 2008). *TOR1* deletion can alleviate loss of survival among yeast starved for leucine (Boer et al. 2008). Rapamycin, a specific inhibitor of TORC1, was examined for modulation of quiescence profiles of the BY4741 and BY4742 genotypes in HL media with or without ammonium sulfate. In a concentration-dependent manner, rapamycin reduced colony forming capacity upon entry to stationary phase; however, cultures with lower initial CFU capacity had proportionally increased maintenance of viability with age (i.e., increased quiescence), so that the CFU capacity of all cultures equalized around Day 38 (**Online Resource 4-Fig. S6**). In BY4741, there was a loss of quiescence after Day 38 associated with ammonium sulfate in the media, which was independent of rapamycin treatment (**Online Resource 4-Fig. S6A**). In BY4742, post-Day 38, rapamycin was associated with concentration-dependent, increased chronological survival; until, by Day 60, BY4742 cultures maintained residual colony forming capacity only if treated with high concentrations (2.5 or 5 nM) of rapamycin. Ammonium sulfate had a greater influence on the quiescence profile of BY4741 than BY4742, however the response to rapamycin was little altered by ammonium sulfate in either strain (**Online Resource 4-Fig. S6A-B**). The observation that rapamycin reduces the initial colony forming capacity of stationary phase cultures in HL media suggests induction of quiescence prior to nutrients being fully exhausted. Conversely, progression to quiescence in the absence of rapamycin results in a higher initial CFU capacity in stationary phase, but with less stable establishment and maintenance of quiescence over the ensuing 35-40 days (**Online Resource 4-Fig. S6A-B**). Interestingly, rapamycin exerts a concentration effect between 0.25 and 5 nM in HL media in both BY4741 and BY4742 prior to Day 38. However, after Day 38, BY4741 reached a stable quiescent state that was reduced by addition of ammonium sulfate to the media, but not further affected by rapamycin exposure. In contrast to BY4741, quiescence of BY4742 was relatively unaffected by ammonium sulfate; however, loss of quiescence for BY4742 after Day 38 was delayed in a rapamycin concentrationdependent manner (**Online Resource 4-Fig. S6A-B**).

Lack of response of BY4741 to rapamycin after Day 32, relative to BY4742, suggested the hypothesis that *met17* auxotrophy (present in BY4741 only) may counteract TORC1 signaling to promote quiescence in a manner hypostatic to rapamycin. To test this possibility, we hypothesized that ammonium sulfate challenge, together with forced activity of TORC1, could induce loss of quiescence. Constitutive expression of *TOR1*, repressible by the addition of doxycycline in HLD media with 2% glucose with or without ammonium sulfate, revealed ammonium sulfate-dependent loss of quiescence around Day 32 (**Online Resource 4-Fig. S6C**). Addition of 5X (~0.35 gm/L) glutamine (a preferred nitrogen source like ammonium sulfate) with forced overexpression of *TOR1* in BY4741 resulted in weak loss of quiescence intermediate between HL media with and without ammonium sulfate (data now shown). The age (~30 days) at which quiescence is reduced by the combination of TOR1 overexpression and ammonium sulfate availability (**Online Resource 4-Fig. S6A-B**) roughly corresponds to when BY4741 establishes stable quiescence (**Figs. 3, 4**), and when the quiescence profiles of rapamycin treated BY4741 and BY4742 cultures converge or diverge (**Online Resource 4-Fig. S6A-B**). Forced expression of *TOR1* did not have this effect in the absence of ammonium sulfate addition to the media (**Online Resource 4-Fig. S6C**). The results suggest that combined effects of ammonium sulfate and TORC1-signalling on quiescence further depend on auxotrophic mutations that differ between BY4741 and BY4742.

Replication stress is implicated in yeast and human aging (Burhans and Weinberger 2012; Flach et al. 2014), alternatively reducing yeast CLS (Weinberger et al. 2007), or increasing it through hormesis (Ross and Maxwell 2018). Methionine restriction (Lee et al. 2016; Petti et al. 2011) and threonine metabolism (Weinberger et al. 2013) have been implicated with respect to mechanisms of replication stress and aging (Liu et al. 2019). The potential influence of methionine auxotrophy on replication stress was investigated by aging BY4741 and BY4742 in HL media with 2% dextrose, in the presence of 0, 30 or 60 mM hydroxyurea (HU) to inhibit ribonucleotide reductase (RNR). Additionally, methionine and cysteine levels, which are convertible by transulfuration of homocysteine and cystathionine (Thomas and Surdin-Kerjan 1997), were perturbed by varying their amounts at .25X or 2X the normal media concentrations. Observations from these quiescence profiles included (**Online Resource 4-Fig. S7**): (1) HU treatment was associated with a dose-dependent reduction of CFU capacity for BY4741 and BY4742 at Days 4 and 6 (**Online Resource 4-Fig. S7A-B**); (2) 1/3X methionine and cysteine was associated with a reduction in CFU capacity (elevation of L) at Day 4 for BY4741 under treatment with 30 mM or 60 mM HU-treatment (**Online Resource 4-Fig. S7A**), followed by gasping (reduction of L) and hormesis-like development of stable quiescence (stable L), which persisted for 96 days; and (3) a consistent trend for auxotrophic genotype and methionine-cysteine limitation to promote quiescence in response to HU-induced replication stress, evidenced by (a) 1/3X methionine-cysteine promoting quiescence in BY4741 at 30 and 60 mM HU (**Online Resource 4-Fig. S7A**), but only at 30 mM HU in BY4742 (which lost colony forming capacity by Day 11 with 60 mM HU) (**Online Resource 4-Fig. S7B**), (b) BY4741 could tolerate 30 mM HU with 1X methionine-cysteine, but lost quiescence at Day 11 when challenged with 60 mM HU, while BY4742 lost quiescence at either 30 or 60 mM HU (**Online Resource 4-Fig. S7A-B**), (c) BY4741 was able to maintain its normal quiescence profile with 2X methionine-cysteine, but in that context could not tolerate either HU concentration, while BY4742 was, regardless of HU challenge, unable to maintain quiescence beyond Day 11 (**Online Resource 4-Fig. S7A-B**). This consistent pattern suggests reduced methionine metabolism can buffer quiescence against replication stress (Hartman IV et al. 2001).

The impact of threonine metabolism, which can buffer replication stress and extend CLS (Burhans and Weinberger 2012; Hartman IV 2007; Weinberger et al. 2013), was tested for potential modulation of quiescence in HL media. 10X threonine supplementation increased CLS of BY4741 through Day 24 (**Online Resource 4-Fig. S7C**), but afterward quiescence was stably maintained independent of threonine. In contrast, the impact of threonine supplementation on BY4742 quiescence profiles occurred only after Day 34, when 5X and 10X threonine promoted maintenance of quiescence (**Online Resource 4-Fig. S7D**).

## Discussion

### A search for strategies to increase reproducibility in CLS studies

This work aimed to understand reasons for disagreement between genome-wide CLS studies reported across independent laboratories. Suspected causes were investigated systematically, including differences in auxotrophic mutations, media composition, aeration, age at which colony forming capacity is assayed, and combinations of factors (Smith et al. 2016). Consideration of auxotrophy was focused on the five loci in the genetic backgrounds for the *MAT**a*** (BY4741) and *MATα* (BY4742) YKO libraries (**Online Resource 2-Table S1**) (Brachmann et al. 1998). HL defined media was developed that increases quiescence, which omits ammonium sulfate and slightly adjusts other nutrients (**Online Resource 2-Table S4**) (Hartman IV et al. 2015). Using a 384-well format, uniform aeration was achieved by inverting the culture plates (Santos and Hartman IV 2019). QHTCP accommodated quiescence profiling at high temporal resolution, using the cell proliferation parameter, L, for sensitive detection of changes in colony forming capacity (**Figs. 1, 2, Online Resource 4-Fig. S1**). Profiles revealed influences of auxotrophy and nutrients at distinct stages of quiescence spanning over 60 days. Aging-relevant perturbations, such as TORC1 signaling and replication stress were also characterized (**Online Resource 4-Figs. S6-7**), and a high throughput pH assay was used to correlate media acidification with auxotrophic status and quiescence (**Fig. 6**).

### Q-HTCP profiles characterize regulation of quiescence establishment and maintenance

The genes and biological processes that enable yeast cells to undergo quiescence for chronological survival are complex, challenging to study, and not fully understood as an integrated system. Analogous issues of complexity have been described for *C. elegans* (Banse et al. 2019; Lucanic et al. 2017). Q-HTCP-based quiescence profiling provides high throughput and phenotypic resolution to control for and survey potential interacting factors that may underlie differential quiescence. The 384-array format and automated imaging system accommodates outgrowth measures to assess colony forming capacity for over 50,000 cultures every 3-4 days. High resolution growth curves report on colony forming capacity in terms of cell proliferation parameters that serve as quantitative phenotypes. Increased temporal resolution reports more dynamically on the establishment and maintenance of stable quiescence, while increased replication of results within each experiment enable characterization of technical and biological variance. Q-HTCP profiling of quiescence reveals a complex developmental process, evidenced by quiescence phenotypes occurring at different ages in response to interacting genetic and metabolic perturbations.

### HL media provides a more optimal defined context for quiescence

Quiescence hinges on cell cycle arrest prior to ‘start’ (Hartwell 1974). One of the early studies about G0 utilized prototrophic strains and documented nutrient depletion elicits efficient and long-term survival of approximately 100 days, noting depletion of auxotrophic nutrients as an exception (Lillie and Pringle 1980). Although appreciated for decades, yeast quiescence remains a complex quantitative trait that is not well understood from a systems perspective (Gresham et al. 2011), perhaps in part because large-scale CLS studies using the YKO libraries have used compound auxotrophic backgrounds and media conditions that are sub-optimal for G0 / quiescence entry. Often the entire lifespan is considered to be less than a 3-4 weeks, and establishment of stable quiescence is not reported. CLS studies, as such, may be thought of to reflect enhancers and suppressors of defective quiescence (Gresham et al. 2011). Building from this perspective, this current study demonstrates that *leu2* auxotrophy alone disrupts the normal quiescence of a prototrophic strain, even with fully complementing leucine in the media. *lys2* auxotrophy is similar in this regard, and the combination of *lys2* and *leu2* further disrupts quiescence, which can be mitigated by *met17* autotrophy.

HL media was developed to optimize quiescence studies by removal of ammonium sulfate (Hess et al. 2006; Santos et al. 2015) and complementation of auxotrophic nutrients (**Fig. 4**). HL media does not support quiescence quite as efficiently as YPD, thus further improvement should be possible in the future. Nevertheless, a consistent percentage of prototrophic cells robustly established and maintained quiescence in HL media. HL media is not so different from human tissue culture media (Hartman IV et al. 2015), and thus genetically disrupting biosynthesis in yeast of nutrients that are essential in humans could be used to model evolutionarily conserved influences of dietary deficiency or supplementation on quiescence.

Genetic analysis of Q-cell characterization is limited, relative to CLS, due to issues of throughput for molecular and cellular characterization of Q-cell qualities in the YKO libraries (Aragon et al. 2008; Werner-Washburne et al. 2012). Undefined media (YPD) is often used to enrich the induction of Q-cells, which are routinely isolated by after 7 days for molecular characterization, including gene expression, glycogen or trehalose content, dye exclusion, and associated cellular characteristics such as stress tolerance, replicative age, or survival in water (Aragon et al. 2008; Werner-Washburne et al. 2012). Recent studies have proposed that mitochondrial network morphology is predictive of colony forming capacity, independent of classical Q-cell characteristics (Laporte et al. 2018). Quiescence profiling in HL media provides a new opportunity to address, using the YKO libraries, the longstanding observation that prototrophic yeast can maintain stable quiescence for months (Lillie and Pringle 1980). The SGA method could be used to construct a prototrophic YKO library (Tong and Boone 2006). Conducting studies in defined HL media would enable systems level identification of nutrient factors and gene-nutrient interaction that influence quiescence. Q-HTCP provides the necessary throughput for quiescence profiling of the YKO library, providing richer phenotypic information to more clearly define and pinpoint, for molecular characterization, complex biological processes that underlie quiescence phenotypically.

### Quiescence profiling temporally resolves complex gene and environment interactions

Most interventions reported here reduced quiescence relative to the prototrophic strain in HL media. The prototrophic strain quiesced similarly if only glucose concentration was varied, but quiescence was lost in the presence of ammonium sulfate at 2% or 5% glucose. BY4742, being auxotrophic for *lys2* and prototrophic for *MET17*, was quiescence-deficient relative to BY4741. However, BY4742, achieved colony forming capacity comparable to the prototroph after Day 35 in 0.4% or 2% glucose *with* ammonium sulfate (**Fig. 3A, 3B**). Quiescence profiles for BY4741, B4742, and BY4712 (auxotrophic only for *leu2*) were each unique, even though quiescence was generally disrupted by increasing glucose and/or ammonium sulfate (**Fig. 3**).

Starvation for many components of media efficiently induces quiescence, while starvation for certain auxotrophic nutrients has been noted to reduce survival in stationary phase (Boer et al. 2008; Lillie and Pringle 1980). In HL media, 1/3X reductions and 3X additions was tested in BY4741, BY4742, single auxotrophs, and the prototroph. In general, restriction of leucine, lysine, or uracil reduced quiescence in the context of a corresponding auxotrophic mutation, which was in some cases modified by the presence of additional auxotrophic loci. Colony forming capacity of prototrophic cells was only slightly, if at all, affected by restriction or supplementation. Loss of quiescence due to *leu2* or *lys2* auxotrophy was only partially restored by auxotrophic nutrient supplementation, suggesting an intrinsic deficiency in adaptive cell cycle exit due to auxotrophy even in the absence of starvation for the corresponding nutrient.

Furthermore, magnitude of effect and the age at which effects of auxotrophic nutrient availability occurred varied between different loci and combinations of loci. HL media fully complemented all auxotrophies in that additional supplementation of the corresponding nutrient had little effect (**Figs. 4, Online Resource 4-Fig. S4-S5**). Systematic analysis of the 32 combinations of the five alleles present in BY4741 and BY4742 were consistent with the phenotypes of the BY strains: quiescence was promoted by *LEU2* and *LYS2* prototrophy, and by *met17* auxotrophy (**Fig. 5A-E**). Effects of *URA3* and *HIS3* loci were negligible if considered in the broader context of *LEU2, LYS2, and MET17*, however smaller influences were observable in specific auxotrophic contexts, *e.g., LEU2 lys2 met17* or *leu2 lys2 MET17* (**Fig. 6 C-D** and **Online Resource 2-Table S5**). Interactions between loci were consistent: *leu2* and *lys2* enhanced loss of quiescence, while *met17* partially suppressed either or the combination (**Figs. 5F-I**). A list of quiescence factors characterized in this study, along with their directions of effect and effects of combination, is summarized in **Fig. 7** and **Online Resource 2-Table S6**.

**Fig. 7.**
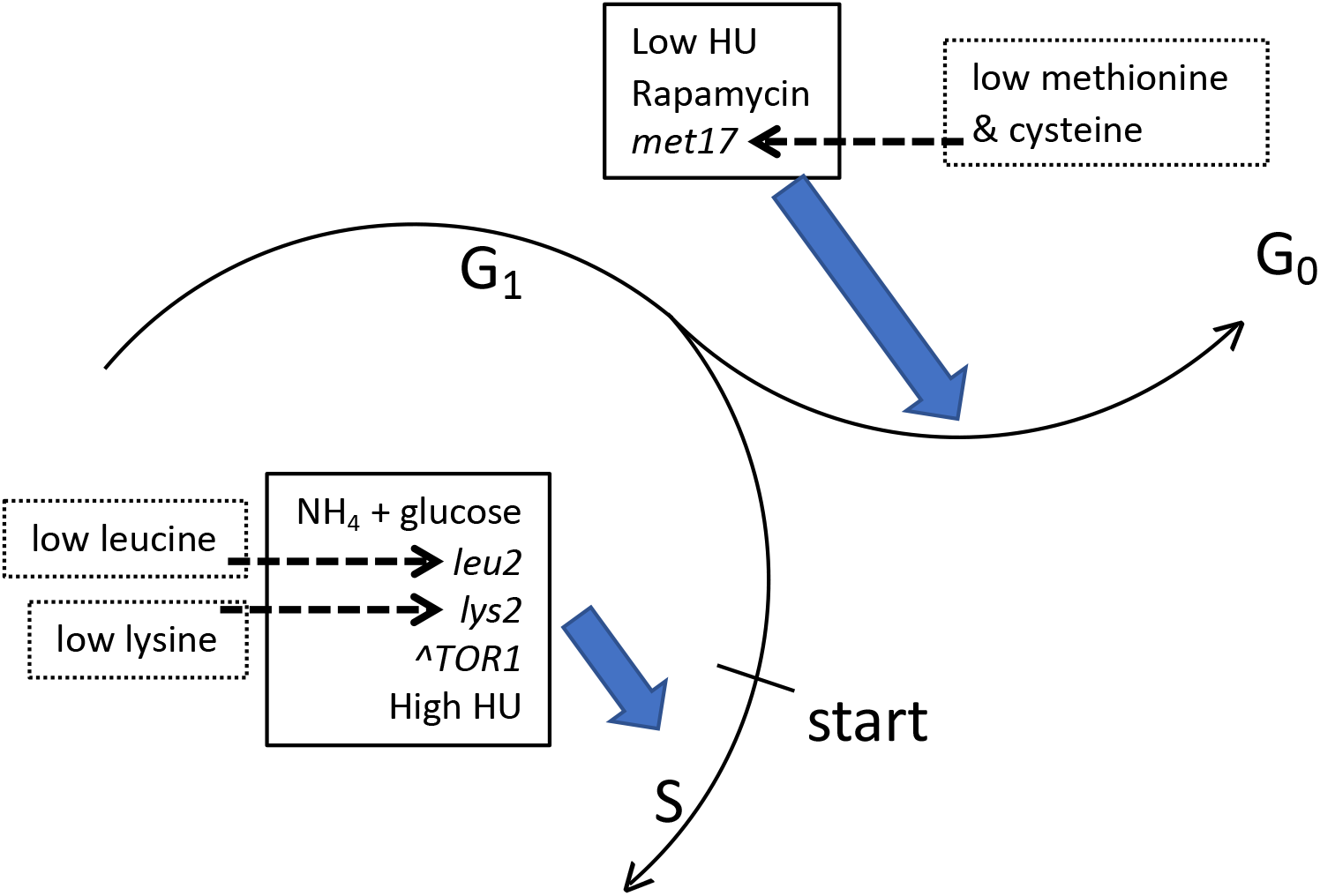
Summary of factors tested and their general impact on quiescence in HL media. Factors that impede or disrupt quiescence under conditions of nutrient depletion are indicated to promote passage of start and entry to S phase. Additional factors and combinatorial influences are summarized in **Online Resource 2-Table S6**. HU - hydroxyurea; NH_4_ - Ammonium sulfate.

### Media acidification and quiescence are not strongly correlated in HL media

Media acidification is generally associated with reduced CLS (Burtner et al. 2009; Fabrizio and Wei 2011; Murakami et al. 2012; Murakami et al. 2011). However, they did not correlate strongly in the context of studies conducted here. In HL media, auxotrophy influences the pH of stationary phase cultures, and these effects further depend on aeration (**Fig. 6E**). Quiescence was less affected by aeration (**Fig. 6F**), indicating effects of auxotrophy on media acidification and quiescence (at least in HL media) are independent and not causally related. For example, *LEU2 LYS2* prototrophic strains exhibited relatively low pH, yet quiesced well; within this group, *met17* auxotrophs exhibited both lower pH and greater quiescence than *MET17* prototrophs (**Fig. 6D**). In development of the pH assay, we observed that HU treatment results in an increase in pH, but lower quiescence (**Online Resource 1; Online Resource 4-Fig. S7**). From these observations, we conclude that media acidification does not strongly correlate with quiescence in all contexts, and thus further studies are needed to clarify causality.

### Characterization of dietary restriction, TORC1 inhibition, and replication stress in HL media

Dietary restriction (DR) and inhibition of nutrient signaling through TORC1, while related aging mechanisms, are also recognized as distinct (Unnikrishnan et al. 2020). DR involves, in addition to TORC1 signaling, other nutrient signaling pathways and the macronutrient composition of the diet (Kapahi et al. 2017; Piper et al. 2011; Soultoukis and Partridge 2016). The yeast quiescence model could serve to map connections between different aging mechanisms. For example, methionine restriction can increase lifespan in different species (Johnson and Johnson 2014) (Hine et al. 2015) (Lee et al. 2016), and numerous effects of *MET17* allele status were observed in these studies. *met17* auxotrophy suppressed loss of quiescence from *leu2* and/or *lys2* auxotrophy (**Fig. 5G-I**). FY4, when challenged with ammonium sulfate and 2% glucose, lost quiescence after Day 42, which was suppressed by *met17* auxotrophy (**Online Resource4-Fig. S3B**). BY4742 quiescence was increased by rapamycin after Day 38, when no effect was seen on BY4741 (**Online Resource4-Fig. S6A-B**). These observations suggest that TORC1 signaling is involved in the reduced quiescence of BY4742, as previously reported (Boer et al. 2008), but that *met17* can be epistatic to the effect. In further support of this hypothesis, forced expression of *TOR1*, together with ammonium sulfate, disrupted quiescence of BY4741 (**Online Resource4-Fig. S6C**). Additional forms of dietary restriction, their interactions, and modulation by evolutionarily conserved nutrient signaling pathways can thus be modeled at a systems level by yeast quiescence profiling.

In a dose-responsive way, rapamycin treatment induced low initial colony forming capacity with proportional increase in quiescence stability in the first 30 days (**Online Resource 4-Fig. S6A-B**). These results, together with differential responses of BY4741 and BY4742 after Day 38, are indicative of distinct developmental stages of quiescence (nutrient depletion, establishment of stable quiescence, and long term maintenance. The “nutrient depletion” response to rapamycin could be related to the prior observation that yeast cells initiate glycogen and trehalose production when half of the initial glucose concentration is consumed, rather than at an absolute level of depletion (Lillie and Pringle 1980). Rapamycin reducing the initial nutrient levels that are sensed, leading to premature but more robust quiescence, may provide a clue to the mechanism, and a molecular strategy to further study the phenomenon.

Replication stress is another biological mechanism of aging (Billard and Poncet 2019; Hämäläinen et al. 2019) across eukaryotes, where yeast CLS serves as a model (Burhans and Weinberger 2012; Ross and Maxwell 2018). We observed effects of replication stress on quiescence to be dependent upon auxotrophic and nutrient status. Limiting methionine and cysteine proportionately increased quiescence in the context of replication stress induced by HU, and this effect was stronger in BY4741 than in BY4742 (**Online Resource 4-Fig. S7A-B**), seemingly due to cooperation with *met17* auxotrophy. Increased resistance to replication stress by BY4741, relative to BY4742, implicates reduced methionine metabolism as a buffering mechanism relatively early (before Day 21) during the development of quiescence. Loss of the capacity to maintain long-term quiescence (beyond Day 30) for BY4742, relative to BY4741 (**Fig. 3A-B**), suggests a later physiological stage, which is revealed in BY4741 by disruption of quiescence after Day 26 by perturbations of ammonium sulfate and forced overexpression of *TOR1* (**Online Resource 4-Fig. S6C**).

### Implications of auxotrophy and media composition for yeast CLS and the biology of aging

This report clarifies in new detail that auxotrophic and nutrient context is critical for yeast quiescence and thus interpretation of CLS effects. To distinguish whether a gene or nutrient has a primary impact on quiescence (rather than modulating an auxotrophic or metabolic influence) the factor should be assessed in a prototrophic context, under optimal and carefully controlled media conditions. Regulators of quiescence and longevity, such as TORC1 signaling or replication stress, may also function in a metabolic context-dependent manner and thus should be studied with auxotrophy and nutrient factors under consideration. The use of a prototrophic strain, highly controlled environmental conditions, and optimal defined media should reveal quiescence factors that have been otherwise identifiable. Likewise, regulators that enhance or suppress auxotrophic or metabolic quiescence factors should be clarified by experiments specifically designed for that purpose. Quiescence profiling would be informative from a systems perspective by perturbing yeast quiescence factors of evolutionary significance, such as *MET17* allele status or TORC1 activity, in combination other quiescence factors.

### Application of other cellular and molecular characterizations to quiescence profiles

The original motivation for quiescence profiling was to functionally characterize factors that could account for non-concordance between CLS studies employing the YKO library (Smith et al. 2016), and thus provide a better foundation for future systems level genetic studies of human relevance (Hartman IV et al. 2015). However, quiescence profiling also shifts emphasis, from identification of genes for which loss of function (YKO) alleviates defective quiescence, to identification of genes primarily required for yeast quiescence. High resolution understanding of the networks of genetic and environmental interactions that regulate the initiation, robust establishment, and long-term maintenance of quiescence will lead to new opportunities for integration of genetics with molecular and cellular studies (Klosinska et al. 2011; Laporte et al. 2018; Miles et al. 2019; Werner-Washburne et al. 2012; Young et al. 2017). Phenomena such as hormesis, gasping, and long-term quiescence could then be characterized in terms of reserve carbohydrates and stress protectants (Gancedo and Flores 2004; Lillie and Pringle 1980; Shi et al. 2010; Sillje et al. 1999; Tapia et al. 2015), replicative age, mitochondrial network morphology (Laporte et al. 2018), or other factors. Interestingly in this regard, Q-cell characteristics such as density, bud scars, and stress resistance (Aragon et al. 2008; Davidson et al. 2011; Young et al. 2017) did not correlate with colony forming capacity examined at the single cell level; rather, cell cycle re-entry was associated with mitochondrial network morphology (Laporte et al. 2018). Q-HTCP-derived quiescence profiles, obtained under conditions controlled for prototrophy, media composition and aeration, could provide novel contextual phenotypes for guiding mechanistic studies that relate to molecular and cellular markers of quiescence.

### An improved approach to identify conserved quiescence factors and their roles in aging

Differential quiescence resulting from compound influences of auxotrophy, media composition and YKO alleles seems to plausibly explain inconsistent results obtained across different genome-wide CLS studies (Gresham et al. 2011; Smith et al. 2016). Using optimal defined media and a prototrophic background for reference, discovery of factors required for quiescence could give rise to genetically tractable characterization of enhancer and suppressor networks modulating quiescence. Yeast is a valuable model in this regard, both for its powerful genetics and the uniformity with which eukaryotic quiescent cell populations can be induced. The genetic rationale for a yeast model of human cell quiescence is rooted in the evolutionary conservation of cell cycle and metabolic regulation. Such a model could be leveraged ultimately to understand relationships between quiescence, aging, and disease in animal models. In animals, the impact of quiescence on aging phenotypes is less obvious than it is for yeast CLS. However, systems level genetic principles of quiescence / G0 from yeast should provide a foundation for clarifying evolutionary conservation of quiescence mechanisms, and ultimately investigating the role of quiescence functions in human aging.

## Conclusions

Quiescence profiling, obtained through high throughput growth curve analysis, suggest that broad inconsistencies between large scale CLS studies with the YKO libraries can be attributed to the large number of possible interactions involving auxotrophy, media composition and environmental conditions. Factors, which have not been controlled between different large scale studies, create a sufficiently complex context for differential genetic interaction that could confound results, making them appear inconsistent. Notwithstanding important biology revealed by CLS studies and informative for aging in animals, we propose a related, but alternate approach that will be more directly informative of human cell quiescence, which is to optimize the conditions for quiescence, discover factors that disrupt it, and then characterize specific factors in a highly controlled manner, and from a phenomic perspective, for their enhancer and suppressor networks. From this modified perspective, a major goal would be to clarify G0 in terms of alternative cell fates (quiescence, senescence, programmed cell death, necrotic death, etc.) as a function of age and relevant genetic and molecular markers. In this way, yeast quiescence is envisioned as a more direct model for human cell *quiescence*, to generate genetic and metabolic hypotheses about which relevance to human *aging* would be best conducted in multicellular models.

## Supporting information

Online Resource 1

Online Resource 2

Online Resource 3

Online Resource 4

## Declarations

### Funding (information that explains whether and by whom the research was supported)

The work was funded by National Institutes of Health, National Institute on Aging R01AG043076 and R56AG059590.

### Conflicts of interest/Competing interests (include appropriate disclosures)

The authors declare no competing of conflicting interests.

### Ethics approval (include appropriate approvals or waivers)

Not applicable.

### Consent to participate (include appropriate statements)

Not applicable.

### Consent for publication (include appropriate statements)

The authors consent for publication.

### Availability of data and material (data transparency)

All data and materials are available upon request.

### Code availability (software application or custom code)

All code will be shared upon request.

### Authors’ contributions (optional: please review the submission guidelines from the journal whether statements are mandatory)

SMS, DLS, and JLH4 designed experiments. SMS, SL, AB, CB, JH, and JLH4 conducted experiments. SMS, JH, JR, and JLH4 performed analyses. SMS and JLH4 wrote the manuscript.

## Acknowledgements

The work was funded by National Institutes of Health, National Institute on Aging R01AG043076 and R56AG059590. The authors thank Haley Albright and Michael Fitch for discussions and Jeff Smith for critiques on the manuscript. We thank Jeff Smith for sharing the FY4 and BY strains derived from FY4 (BY4700, BY4706, BY4712, BY4730) by the Boeke laboratory.

