## Supplementary material for "High-resolution yeast quiescence profiling in human-like media reveals interacting influences of auxotrophy and nutrient availability": Online Resource 1

**Online Resource 1. Additional description of experimental procedures, additional figure legends (for OR 3 and 4), and additional table legends (for OR2)**.

*Generation of strains expressing combinations of auxotrophic alleles*

A BY/FY4 diploid, heterozygous for the five auxotrophic markers used in the YKO libraries, was made by mating the met17Δ0 deletion mutant from the *MATα* YKO library (*MATα his3Δ1, leu2Δ0, lys2Δ0, met17Δ0,* and *ura3Δ0*) to the FY4 prototroph (*MAT**a***). A spore obtained from tetrad dissection with the genotype (*MAT**a*** lys2Δ0) was mated to a complementary spore (*MATα his3Δ1 leu2Δ0 LYS2 met17Δ0 ura3Δ0*) obtained from dissection of BY4743, creating a FY4/BY hybrid that was heterozygous at all five auxotrophic loci. It was sporulated and dissected to create Auxotrophic Panel 1, which was used in the analysis presented in **Fig. 5**. Tetrads were arranged on 96-well plates, transferred to 384 well plates, and scored for auxotrophy by Q-HTCP after transfer to dropout media (**Online Resource 2-Table S2A**). Mating type was determined using an *ilv2* auxotroph mutant as a tester (Konopka et al. 1988), scoring growth on B media (Burke et al. 2000). Thus, mating type was undetermined for fully prototrophic strains. A third backcross was performed by selecting a spore from the Auxotrophic Panel 1 with the *MATα lys2Δ0* genotype (arising from tetrad 85 in Table S2A) and backcrossing it to BY4741. Tetrad dissection of this resulting diploid (heterozygous at all five auxotrophic loci) gave rise to Auxotrophic Panel 2 (**Online Resource 2-Table S2B**), which was used for correlation of media acidification and quiescence (**Fig. 6)**.

*HL media recipe*

(1) Custom YNB: (modifications from standard YNB are highlighted in yellow, purchased from Sunrise Science; https://sunrisescience.com/)

Add0.76 gm/L of YNB.

| Ingredient | mg/L |
|---|---|
| Sodium chloride | 100 |
| Potassium phosphate, monobasic anhydrous | 500 |

| | |
|---|---|
| Calcium chloride | 100 |
| D-Biotin | 0.002 |
| Pantothenic acid | 0.40 |
| Folic acid | 0.002 |
| Inositol | 2.00 |
| Nicotinic acid | 0.40 |
| 4-aminobenzoic acid (PABA) | 0.20 |
| Pyridoxine hydrochloride | 0.40 |
| Riboflavin | 0.20 |
| Thiamine hydrochloride | 0.40 |
| Boric acid | 0.500 |
| Copper sulfate (II) | 0.040 |
| Manganese sulfate | 0.400 |
| Sodium molybdate | 0.200 |
| Zinc sulfate monohydrate | 0.400 |
| Potassium iodide | 0.100 |
| Iron chloride (III) | 0.200 |
| Magnesium sulfate, anhydrous | 50 |
| **Total** | **755.84** |

(2) _Amino acid mix_: (modifications from Cold Spring Harbor recipe are highlighted in yellow, and PABA was omitted: Burke D, Dawson D, Stearns T: *Methods in Yeast Genetics.* Plainview, NY: CSHL Press; 2000.)

Add 1.66 gm/L of AA mixture.

| **Amino Acid Mixture** | amount per liter. |
|---|---|
| Adenine | 0.0183 |
| Alanine | 0.0734 |
| Arginine | 0.0734 |
| Asparagine | 0.0734 |
| Aspartic Acid | 0.0734 |
| Cysteine | 0.0734 |
| Glutamic Acid | 0.0734 |
| Glycine | 0.0734 |
| Histidine | 0.0734 |
| Inositol | 0.0250 |
| Isoleucine | 0.0734 |

| | |
|---|---|
| Leucine | 0.1468 |
| Lysine | 0.0734 |
| Methionine | 0.0734 |
| Phenylalanine | 0.0734 |
| Proline | 0.0734 |
| Serine | 0.0734 |
| Threonine | 0.0734 |
| Tryptophan | 0.0734 |
| Tyrosine | 0.0734 |
| Uracil | 0.0734 |
| Valine | 0.0734 |
| Glutamine | 0.0734 |
| | **1.66** |

YNB + glucose + ammonium sulfate (if added) are autoclaved; AA powder added after autoclaving. Liquid media was filtered for aging assays.

Additional recipe details in **Online Resource 2-Table S4**.

*Q-HTCP (quantitative high throughput cell array phenotyping)*

In contrast to L, which is a very sensitive estimate of changes in colony forming unit capacity (main figures and Online Resource 4)), carrying capacity (K) and the maximums specific rate (r) are cruder measure of larger changes in quiescence (Online Resource 3).

*High throughput pH assay with Bromophenol Blue as pH indicator dye*

We noted that buffered standards, such as those prepared by the Carmody Buffer System (https://pubs.acs.org/doi/10.1021/ed038p559#) did not translate well between direct measurement with a pH electrode and estimation of pH for stationary phase cultures by standard curve created with use of pH indicator dyes (data not shown). We concluded that this resulted from differential interaction of the pH indicator dyes with stationary phase media and buffered solutions. Reliable estimates could be obtained using stationary phase media to construct a standard curve, and so conditions to obtain stationary phase media having a range of pH were identified. pH was dynamic over a 10-day period in HL media with glucose concentration ranging0.0.1%-1.0% or with HU ranging 2.5 mM to 60 mM. (see below)

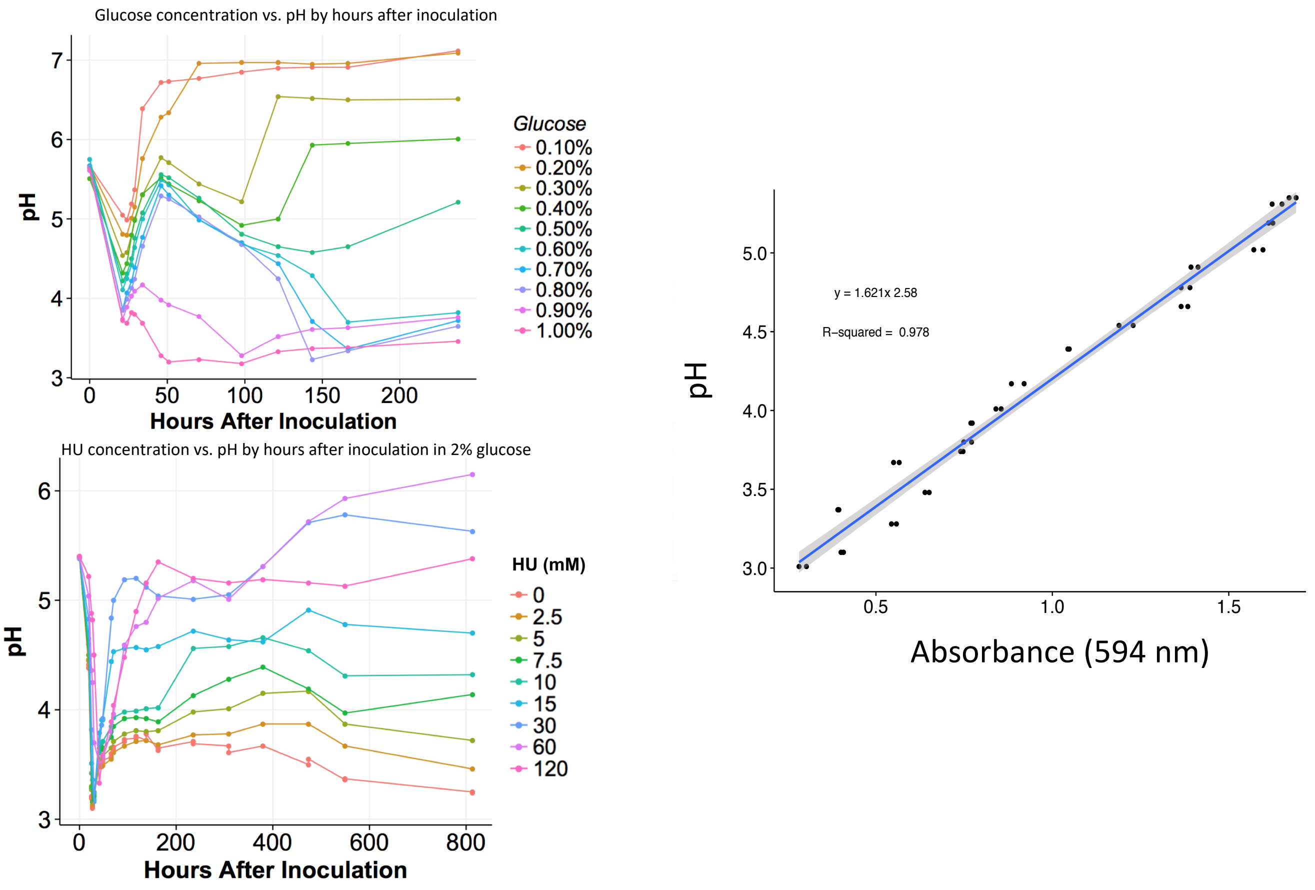


**Figure above:** Large flask cultures were prepared as indicated (left panels). 10 mL of culture was collected at the indicated time points, cells were cleared by centrifugation and pH was measured directly with an electrode. Selected samples, stored at -20 °C, were used to generate a standard curve (right panel), using bromophenol blue as a pH indicator to estimate the pH of 10 uL samples of stationary phase cultures in 384-array format. The assay was validated by strong correlation between pH of unknown samples estimated by the assay and also measured directly by electrode.

*Correlation of media acidification with quiescence*

Aging cell arrays were inoculated from a single source and analyzed in triplicate under non-aerated (upright plates) and aerated (inverted plates) conditions. Standard high-profile plates were used (see methods) in order to obtain enough conditioned media for pH measurements

(**Fig. 6**). At Day 31, 10 uL of conditioned media was removed from each culture for pH measurement (**Fig. 6**). Quiescence was assessed at Day 42.

**Legends for Online Resource 2 (Tables):**

Table S1. 'BY genotypes' - pre-existing strains used in this study (also, see Fig. S1).

Table S2A. 'Aux Panel 1' - Auxotrophic Panel 1 of strains obtained by tetrad dissection of FY4/BY heterozygous diploid (*MAT**a**/MATα HIS3/his3 LEU2/leu2 LYS2/lys2 MET17/met17 URA3/ura3*). Panel used for experiment shown in Fig. 5.

Table S2B. 'Aux Panel 2' - Auxotrophic Panel 2 of strains obtained by backcross of spore (85_1) from Aux Panel 1 with BY4741, followed by repeating tetrad dissection. This strain panel was used for experiments shown in Fig. 6.

Table S3. 'spore viability' - Viability of spores and tetrads from FY4 X BY backcrosses (Aux Panels 1 and 2 in Table 2).

Table S4. 'HL media recipe' – Composition of "human-like" media.

Table S5. 'L_pH correlation' – Data plotted in **Fig. 6C-F**.

Table S6. 'summary'. Overview of factors that, in combination, differentially influence quiescence profiles in HL media. "X" indicates combination-dependent factors; white indicates effects with influences that are not combination-dependent; gray indicates combinations not tested. Black areas indicate redundant matrix.

**Legends for Online Resource 3 (K plots):**

*Quiescence profiles with K values corresponding to L data from main manuscript and **Online Resource 4** (See legends for those figures for more detailed description). BY strain genotypes are listed in **Online Resource 2-Table S1**. Figure labels indicate the corresponding figure for which the K plots were made. For example, Fig. 3_K corresponds to Fig. 3.*

**Fig. 3_K.** Quiescence profiles for BY4741, BY4742, BY4712, and FY4; HLD media with 0.4, 2, and 5% glucose +/- 0.5 g/L ammonium sulfate.

**Fig. S2_K.** Quiescence profiles for BY4730 (*HIS3 leu2 met17 ura3*) vs. BY4741 (*his3 leu2 met17 ura3*) and FY4; HLD media with 0.4, 2, and 5% glucose +/- 0.5 g/L ammonium sulfate.

**Fig. S3_K.** Quiescence profiles for BY4700 (*ura3*) and BY4706 (*met17*) and FY4; HLD media with 0.4, 2, and 5% glucose +/- 0.5 g/L ammonium sulfate.

**Fig. 4_K. (A)** BY4741, BY4742, BY4712 (*leu2*), and FY4; HLD media with 2% glucose with 1/3X, 1X, or 3X leucine; **(B)** BY4741, BY4742 (*lys2*), and FY4; HLD media with 2% glucose with 1/3X, 1X, or 3X lysine; **(C)** BY4741, BY4742, BY4700 (*ura3*), and FY4; HLD media with 2% glucose with 1/3X, 1X, or 3X uracil; **(D)** BY4741, BY4742, BY4706 (*met17*), and FY4; 2% HLD media with glucose with 1/3X, 1X, or 3X methionine.

**Fig. S4_K.** Quiescence profiles for BY4730 (*HIS3 leu2 met17 ura3*) vs. BY4741 (*his3 leu2 met17 ura3*) and FY4; HLD media with 2% glucose with 1/3X, 1X, or 3X leucine.

**Fig. S5_K. (A)** BY4706 (*met17*), BY4700 (*ura3*), and FY4; HLD media with 2% glucose with 1/3X, 1X, or 3X leucine; **(B)** BY4712 (*leu2*), BY4706 (*met17*), BY4700 (*ura3*), and FY4; HLD media with 2% glucose with 1/3X, 1X, or 3X lysine; **(C)** BY4712 (*leu2*), BY4706 (*met17*), and FY4; HLD media with 2% glucose with 1/3X, 1X, or 3X uracil; **(D)** BY4712 (*leu2*), BY4700 (*ura3*), and FY4; HLD media with 2% glucose with 1/3X, 1X, or 3X methionine.

**Fig. S6_K. (A)** BY4741 and **(B)** BY4742; HLD media with 2% glucose +/- 0.5 gm/L ammonium sulfate with indicated rapamycin concentrations. **(C)** BY4741; HLD media with 2% glucose +/- 0.5 gm/L ammonium sulfate with indicated doxycycline concentrations.

**Fig. S7_K. (A)** BY4741 and **(B)** BY4742; HLD media with 2% glucose with 0.25X, 1X or 2X methionine/cysteine concentrations, treated with HU concentrations of 0, 30 and 60 mM. **(C)** BY4741 and **(D)** BY4742; HLD media with 2% glucose, with threonine concentrations varied from 0.25X to 10X the normal concentrations.

**Legends for Online Resource 4 (additional figures):**

**Fig. S1. Additional description and illustration of Q-HTCP-derived quiescence profiles.**
***Fig. S1A***- *Relationship between L and colony forming capacity. (i)* Q-HTCP reports the value of L, which is defined as the time when half of carrying capacity (K/2) occurs. *(ii)* Representative image time series for a culture spotted after serial dilution, with the approximate time that the L value is achieved indicated by the boxes. *(iii)* Raw data and fitted curves from images in panel ii (Shah et al. 2007). *(iv)* Increasing L (corresponding to panels ii and iii) reports reduced colony forming capacity, *e.g.*, loss of quiescence. *(v)* Using a doubling time of 1.5 hours for example, and assuming constant stationary phase cell density, the table illustrates how increasing L reports on colony forming capacity and thus viability and quiescence.
***Fig. S1B*** - *Example array images from a quiescence profiling experiment. (i)* Layout of 7 strains on a quiescence profiling array, corresponding to panels ii and iii. *(ii)* Example Q-HTCP images from outgrowth analysis at Days 11, 19, 25, and 33 (corresponding to **Fig. 2**). Rows reflect increasing age (labeled at left, ordered top to bottom). Columns reflect increasing time of outgrowth (labeled at bottom). *(iii)* Strain genotypes.
***Fig. S1C*** – *Correlation between quiescence profile data (using L parameter from Q-HTCP growth curves) and traditional serial dilution spot test*. *(i)* Quiescence profiles for FY4 in HL media, comparing 4% and 0.4% glucose with and without 0.5 gm/L ammonium sulfate. *(ii)* Expanded view of Day 97 data (inset from panel A). *(iii)* Q-HTCP image, after ~22 hours outgrowth, for 12 replicate cultures of each condition, assayed for quiescence at Day 97. *(iv)* Serial dilution test, performed in parallel from the same stationary phase cultures as panel C, ordered by L values indicated in panel v. *(v)* Relational data for panels iii and iv, *i.e.*, L values (derived from analyses of entire time series of images associated with each of the 48 spot cultures depicted, at 22 hours, in panel iii, and summarized in panel ii) corresponding to serial dilutions (shown in panel iv).

**Fig. S2. Complement to Fig. 3, addressing the effect of his3 auxotrophy** Quiescence profiles for BY4730 (*HIS3 leu2 met17 ura3*) vs. BY4741 (*his3 leu2 met17 ura3*) and FY4; HLD media +/- 0.5 g/L ammonium sulfate, with (**A**) 0.4%, (**B**) 2.0%, or (**C**) 5% glucose. Only the

*HIS3/his3* allele varies between these two strains, thus the similarity of profiles suggests *his3* auxotrophy has no effect in this context.

**Fig. S3. Complement to Fig. 3, addressing the effect of *ura3* or *met17* auxotrophy**. Quiescence profiles for BY4700 (*ura3*) and BY4706 (*met17*) and FY4; HLD media, +/- 0.5 g/L ammonium sulfate, with (**A**) 0.4%, (**B**) 2.0%, or (**C**) 5% glucose.

**Fig. S4. Complement to Fig. 4, addressing nutrient limitations of additional strains.** Quiescence profiles for BY4730 (*HIS3 leu2 met17 ura3*) vs. BY4741 (*his3 leu2 met17 ura3*) and FY4; HLD media with 2% glucose with 1/3X, 1X, or 3X leucine. Only the HIS3/his3 allele varies between these two strains, thus the similarity of profiles suggests *his3* auxotrophy has no effect in this context.

**Fig. S5. Complement to Fig. 4, addressing nutrient limitations of additional strains**. **(A)** BY4706 (*met17*), BY4700 (*ura3*), and FY4; HLD media with 2% glucose with 1/3X, 1X, or 3X leucine; **(B)** BY4712 (*leu2*), BY4706 (*met17*), BY4700 (*ura3*), and FY4; 2% glucose with 1/3X, 1X, or 3X lysine; **(C)** BY4712 (*leu2*), BY4706 (*met17*), and FY4; 2% glucose with 1/3X, 1X, or 3X uracil; **(D)** BY4712 (*leu2*), BY4700 (*ura3*), and FY4; 2% glucose with 1/3X, 1X, or 3X methionine.

**Fig. S6. Effects of TORC1 perturbation on quiescence profiles in HL media.** (**A**) BY4741 (*MAT**a** his3 leu2 met17 ura3*) and (**B**) BY4742 (*MAT*$\alpha$ *his3 leu2 lys2 ura3*) were inoculated into HLD media with 2% glucose with or without 0.5 gm/L ammonium sulfate and colony forming capacity was assessed by Q-HTCP (x-axis) at the indicated ages of cultures (y-axis). (**C**) BY4741, harboring a chromosomal promoter replacement at the *TOR1* locus for doxycycline-repressible gene expression, was aged in HL media with 2% glucose with or without 0.5 gm/L ammonium sulfate and the indicated amounts of doxycycline. Each condition is N=32; comprised of 16 cultures per 384-culture array X 2 arrays.

**Fig. S7. Effects of replication stress on quiescence are modified by auxotrophy, age, and methionine / cysteine availability.** (**A-B**) (A) BY4741 and (B**)** BY4742 were aged in HLD media with 2% glucose, and compared to cultures with methionine/cysteine concentrations increased (2X) or decreased (0.25X) from the normal recipe. Each culture type was treated with HU concentrations of 0, 30 and 60 mM, as indicated. (**C-D**) (C) BY4741 and (D**)** BY4742 were aged in HLD media with 2% glucose, with threonine concentrations decreased (0.25X) or increased (10X) from the normal recipe.
