## Supplementary material for "High-resolution yeast quiescence profiling in human-like media reveals interacting influences of auxotrophy and nutrient availability": Online Resource 2

**Online Resource 2. Tables S1-S6 Legends.**

Table S1. 'BY genotypes' - pre-existing strains used in this study (also, see Fig. S1).

Table S2A. 'Aux Panel 1' - Auxotrophic Panel 1 of strains obtained by tetrad dissection of FY4/BY heterozygous diploid (*MAT* **a**/*MAT* alpha *HIS3/his3 LEU2/leu2 LYS2/lys2 MET17/met17 URA3/ura3* ). Panel used for experiment shown in Fig. 5.

Table S2B. 'Aux Panel 2' - Auxotrophic Panel 2 of strains obtained by backcross of spore (85_1) from Aux Panel 1 with BY4741, repeating tetrad dissection. Panel used for experiment shown in Fig. 6.

Table S3. 'sporeviability' - Viability of spores and tetrads from FY4 X BY backcrosses (Aux Panels 1 and 2 in Table 2).

Table S4. 'HL media recipe' – Composition of "human-like" media.

Table S5. 'L_pH correlation' – Data plotted in Fig. 6C-F.

Table S6. 'summary'. Overview of factors that, in combination, differentially influence quiescence profiles in HL media. "X" indicates combination-dependent factors; white indicates effects with influences that are not combination-dependent; gray indicates combinations not tested. Black areas indicate redundant matrix.

| Table S1. | | | | | | |
|---|---|---|---|---|---|---|
| **Strain** | ***MAT*** | ***HIS3/his3*** | ***LEU2/leu2*** | ***LYS2/lys2*** | ***MET17/met17*** | ***URA3/ura3*** |
| **BY4700** | **a** | *HIS3* | *LEU2* | *LYS2* | *MET17* | *ura3-Δ0* |
| **BY4706** | **a** | *HIS3* | *LEU2* | *LYS2* | *met17-Δ0* | *URA3* |
| **BY4712** | **a** | *HIS3* | *leu2-Δ0* | *LYS2* | *MET17* | *URA3* |
| **BY4730** | **a** | *HIS3* | *leu2-Δ0* | *LYS2* | *met17-Δ0* | *ura3-Δ0* |
| **BY4741** | **a** | *his3-Δ1* | *leu2-Δ0* | *LYS2* | *met17-Δ0* | *ura3-Δ0* |
| **BY4742** | **a** | *his3-Δ1* | *leu2-Δ0* | *lys2--Δ0* | *MET17* | *ura3-Δ0* |
| **FY4** | **a** | *HIS3* | *LEU2* | *LYS2* | *MET17* | *URA3* |

| Table S2B. | | | | | | | | | | |
|---|---|---|---|---|---|---|---|---|---|---|
| Num. | Genotype (M. | Plate_96 | Row_96 | Col_96 | concat96 | Plate_384 | Row_384 | Col_384 | tetrad | spore |
| 1 | APPMPM | 1 | A | 1 | 1_A_1 | 1 | A | 1 | 1 | 1 |
| 2 | BPPMMP | 2 | A | 1 | 2_A_1 | 1 | A | 2 | 25 | 1 |
| 3 | BMMMPP | 1 | A | 2 | 1_A_2 | 1 | A | 3 | 1 | 2 |
| 4 | AMMPPM | 2 | A | 2 | 2_A_2 | 1 | A | 4 | 25 | 2 |
| 5 | APMPMP | 1 | A | 3 | 1_A_3 | 1 | A | 5 | 1 | 3 |
| 6 | BMMMMM | 2 | A | 3 | 2_A_3 | 1 | A | 6 | 25 | 3 |
| 7 | BMPPMM | 1 | A | 4 | 1_A_4 | 1 | A | 7 | 1 | 4 |
| 8 | IPPPPP | 2 | A | 4 | 2_A_4 | 1 | A | 8 | 25 | 4 |
| 9 | AMPMMM | 1 | A | 5 | 1_A_5 | 1 | A | 9 | 2 | 1 |
| 10 | AMMMMM | 2 | A | 5 | 2_A_5 | 1 | A | 10 | 26 | 1 |
| 11 | BPMMPP | 1 | A | 6 | 1_A_6 | 1 | A | 11 | 2 | 2 |
| 12 | BMMPPP | 2 | A | 6 | 2_A_6 | 1 | A | 12 | 26 | 2 |
| 13 | BPPMMM | 1 | A | 7 | 1_A_7 | 1 | A | 13 | 2 | 3 |
| 14 | AMPMPM | 2 | A | 7 | 2_A_7 | 1 | A | 14 | 26 | 3 |
| 15 | AMMPPP | 1 | A | 8 | 1_A_8 | 1 | A | 15 | 2 | 4 |
| 16 | BPPMMM | 2 | A | 8 | 2_A_8 | 1 | A | 16 | 26 | 4 |
| 17 | BPMMPM | 1 | A | 9 | 1_A_9 | 1 | A | 17 | 3 | 1 |
| 18 | BPPMPM | 2 | A | 9 | 2_A_9 | 1 | A | 18 | 27 | 1 |
| 19 | AMPPMP | 1 | A | 10 | 1_A_10 | 1 | A | 19 | 3 | 2 |
| 20 | BMMPPP | 2 | A | 10 | 2_A_10 | 1 | A | 20 | 27 | 2 |
| 21 | BPPMMP | 1 | A | 11 | 1_A_11 | 1 | A | 21 | 3 | 3 |
| 22 | APMMMP | 2 | A | 11 | 2_A_11 | 1 | A | 22 | 27 | 3 |
| 23 | AMMPPM | 1 | A | 12 | 1_A_12 | 1 | A | 23 | 3 | 4 |
| 24 | AMPPMM | 2 | A | 12 | 2_A_12 | 1 | A | 24 | 27 | 4 |
| 25 | BMMPPM | 4 | A | 1 | 4_A_1 | 1 | B | 1 | 73 | 1 |
| 26 | BPPPMM | 3 | A | 1 | 3_A_1 | 1 | B | 2 | 49 | 1 |
| 27 | BMPMMP | 4 | A | 2 | 4_A_2 | 1 | B | 3 | 73 | 2 |
| 28 | AMMPMP | 3 | A | 2 | 3_A_2 | 1 | B | 4 | 49 | 2 |
| 29 | APPMPP | 4 | A | 3 | 4_A_3 | 1 | B | 5 | 73 | 3 |
| 30 | BMPMPM | 3 | A | 3 | 3_A_3 | 1 | B | 6 | 49 | 3 |
| 31 | APMPMM | 4 | A | 4 | 4_A_4 | 1 | B | 7 | 73 | 4 |
| 32 | APMMPP | 3 | A | 4 | 3_A_4 | 1 | B | 8 | 49 | 4 |
| 33 | AMMMPP | 4 | A | 5 | 4_A_5 | 1 | B | 9 | 74 | 1 |
| 34 | AMPPMM | 3 | A | 5 | 3_A_5 | 1 | B | 10 | 50 | 1 |
| 35 | BPPMPM | 4 | A | 6 | 4_A_6 | 1 | B | 11 | 74 | 2 |
| 36 | AMMMPP | 3 | A | 6 | 3_A_6 | 1 | B | 12 | 50 | 2 |
| 37 | APPPMM | 4 | A | 7 | 4_A_7 | 1 | B | 13 | 74 | 3 |
| 38 | BPPMMM | 3 | A | 7 | 3_A_7 | 1 | B | 14 | 50 | 3 |
| 39 | BMMPMP | 4 | A | 8 | 4_A_8 | 1 | B | 15 | 74 | 4 |
| 40 | BPMPPP | 3 | A | 8 | 3_A_8 | 1 | B | 16 | 50 | 4 |
| 41 | BMPMPP | 4 | A | 9 | 4_A_9 | 1 | B | 17 | 75 | 1 |
| 42 | BMPMMM | 3 | A | 9 | 3_A_9 | 1 | B | 18 | 51 | 1 |
| 43 | APMPMM | 4 | A | 10 | 4_A_10 | 1 | B | 19 | 75 | 2 |

| | | | | | | | | | | |
|---|---|---|---|---|---|---|---|---|---|---|
| 44 | BPMPMP | 3 | A | 10 | 3_A_10 | 1 | B | 20 | 51 | 2 |
| 45 | APMPPM | 4 | A | 11 | 4_A_11 | 1 | B | 21 | 75 | 3 |
| 46 | AMPMPP | 3 | A | 11 | 3_A_11 | 1 | B | 22 | 51 | 3 |
| 47 | BMPMMP | 4 | A | 12 | 4_A_12 | 1 | B | 23 | 75 | 4 |
| 48 | APMPPM | 3 | A | 12 | 3_A_12 | 1 | B | 24 | 51 | 4 |
| 49 | BMPMPP | 1 | B | 1 | 1_B_1 | 1 | C | 1 | 4 | 1 |
| 50 | AMMPPM | 2 | B | 1 | 2_B_1 | 1 | C | 2 | 28 | 1 |
| 51 | BPMPMM | 1 | B | 2 | 1_B_2 | 1 | C | 3 | 4 | 2 |
| 52 | AMPMMP | 2 | B | 2 | 2_B_2 | 1 | C | 4 | 28 | 2 |
| 53 | APMPMM | 1 | B | 3 | 1_B_3 | 1 | C | 5 | 4 | 3 |
| 54 | BPMPPM | 2 | B | 3 | 2_B_3 | 1 | C | 6 | 28 | 3 |
| 55 | IPPMPP | 1 | B | 4 | 1_B_4 | 1 | C | 7 | 4 | 4 |
| 56 | BPPMMP | 2 | B | 4 | 2_B_4 | 1 | C | 8 | 28 | 4 |
| 57 | AMPMPP | 1 | B | 5 | 1_B_5 | 1 | C | 9 | 5 | 1 |
| 58 | BPPMPP | 2 | B | 5 | 2_B_5 | 1 | C | 10 | 29 | 1 |
| 59 | BPPPMP | 1 | B | 6 | 1_B_6 | 1 | C | 11 | 5 | 2 |
| 60 | AMMMMM | 2 | B | 6 | 2_B_6 | 1 | C | 12 | 29 | 2 |
| 61 | BMMPMM | 1 | B | 7 | 1_B_7 | 1 | C | 13 | 5 | 3 |
| 62 | APMPPM | 2 | B | 7 | 2_B_7 | 1 | C | 14 | 29 | 3 |
| 63 | APMMPM | 1 | B | 8 | 1_B_8 | 1 | C | 15 | 5 | 4 |
| 64 | BMPPMP | 2 | B | 8 | 2_B_8 | 1 | C | 16 | 29 | 4 |
| 65 | BPMPMP | 1 | B | 9 | 1_B_9 | 1 | C | 17 | 6 | 1 |
| 66 | BPMMMM | 2 | B | 9 | 2_B_9 | 1 | C | 18 | 30 | 1 |
| 67 | APMPPP | 1 | B | 10 | 1_B_10 | 1 | C | 19 | 6 | 2 |
| 68 | IPPPPP | 2 | B | 10 | 2_B_10 | 1 | C | 20 | 30 | 2 |
| 69 | AMPMPM | 1 | B | 11 | 1_B_11 | 1 | C | 21 | 6 | 3 |
| 70 | AMMMMM | 2 | B | 11 | 2_B_11 | 1 | C | 22 | 30 | 3 |
| 71 | BMPMMM | 1 | B | 12 | 1_B_12 | 1 | C | 23 | 6 | 4 |
| 72 | AMPPPP | 2 | B | 12 | 2_B_12 | 1 | C | 24 | 30 | 4 |
| 73 | BMPPMP | 4 | B | 1 | 4_B_1 | 1 | D | 1 | 76 | 1 |
| 74 | BPPMPM | 3 | B | 1 | 3_B_1 | 1 | D | 2 | 52 | 1 |
| 75 | APMMMM | 4 | B | 2 | 4_B_2 | 1 | D | 3 | 76 | 2 |
| 76 | APMPMP | 3 | B | 2 | 3_B_2 | 1 | D | 4 | 52 | 2 |
| 77 | AMMPPM | 4 | B | 3 | 4_B_3 | 1 | D | 5 | 76 | 3 |
| 78 | BMMPMM | 3 | B | 3 | 3_B_3 | 1 | D | 6 | 52 | 3 |
| 79 | BPPMPP | 4 | B | 4 | 4_B_4 | 1 | D | 7 | 76 | 4 |
| 80 | AMPMPM | 3 | B | 4 | 3_B_4 | 1 | D | 8 | 52 | 4 |
| 81 | BPMMPM | 4 | B | 5 | 4_B_5 | 1 | D | 9 | 77 | 1 |
| 82 | AMPMMP | 3 | B | 5 | 3_B_5 | 1 | D | 10 | 53 | 1 |
| 83 | AMMPPP | 4 | B | 6 | 4_B_6 | 1 | D | 11 | 77 | 2 |
| 84 | APMPPM | 3 | B | 6 | 3_B_6 | 1 | D | 12 | 53 | 2 |
| 85 | BMPPMP | 4 | B | 7 | 4_B_7 | 1 | D | 13 | 77 | 3 |
| 86 | BMMMMM | 3 | B | 7 | 3_B_7 | 1 | D | 14 | 53 | 3 |
| 87 | APPMMM | 4 | B | 8 | 4_B_8 | 1 | D | 15 | 77 | 4 |
| 88 | IPPPPP | 3 | B | 8 | 3_B_8 | 1 | D | 16 | 53 | 4 |

| | | | | | | | | | | |
|---|---|---|---|---|---|---|---|---|---|---|
| 89 | BPMMMM | 4 | B | 9 | 4_B_9 | 1 | D | 17 | 78 | 1 |
| 90 | APPPMP | 3 | B | 9 | 3_B_9 | 1 | D | 18 | 54 | 1 |
| 91 | APMPPM | 4 | B | 10 | 4_B_10 | 1 | D | 19 | 78 | 2 |
| 92 | BMPPMP | 3 | B | 10 | 3_B_10 | 1 | D | 20 | 54 | 2 |
| 93 | BMPPMP | 4 | B | 11 | 4_B_11 | 1 | D | 21 | 78 | 3 |
| 94 | AMMMPM | 3 | B | 11 | 3_B_11 | 1 | D | 22 | 54 | 3 |
| 95 | AMPMPP | 4 | B | 12 | 4_B_12 | 1 | D | 23 | 78 | 4 |
| 96 | BPMMPM | 3 | B | 12 | 3_B_12 | 1 | D | 24 | 54 | 4 |
| 97 | AMMPPP | 1 | C | 1 | 1_C_1 | 1 | E | 1 | 7 | 1 |
| 98 | APPPPM | 2 | C | 1 | 2_C_1 | 1 | E | 2 | 31 | 1 |
| 99 | APPPMM | 1 | C | 2 | 1_C_2 | 1 | E | 3 | 7 | 2 |
| 100 | APMPMP | 2 | C | 2 | 2_C_2 | 1 | E | 4 | 31 | 2 |
| 101 | BMMMMP | 1 | C | 3 | 1_C_3 | 1 | E | 5 | 7 | 3 |
| 102 | BMPMPP | 2 | C | 3 | 2_C_3 | 1 | E | 6 | 31 | 3 |
| 103 | BPMMPM | 1 | C | 4 | 1_C_4 | 1 | E | 7 | 7 | 4 |
| 104 | BMMMMM | 2 | C | 4 | 2_C_4 | 1 | E | 8 | 31 | 4 |
| 105 | APMMMM | 1 | C | 5 | 1_C_5 | 1 | E | 9 | 8 | 1 |
| 106 | BPPMMM | 2 | C | 5 | 2_C_5 | 1 | E | 10 | 32 | 1 |
| 107 | AMMPPM | 1 | C | 6 | 1_C_6 | 1 | E | 11 | 8 | 2 |
| 108 | BMMPMP | 2 | C | 6 | 2_C_6 | 1 | E | 12 | 32 | 2 |
| 109 | BMPMPP | 1 | C | 7 | 1_C_7 | 1 | E | 13 | 8 | 3 |
| 110 | AMMPPP | 2 | C | 7 | 2_C_7 | 1 | E | 14 | 32 | 3 |
| 111 | BPPPMP | 1 | C | 8 | 1_C_8 | 1 | E | 15 | 8 | 4 |
| 112 | APPMPM | 2 | C | 8 | 2_C_8 | 1 | E | 16 | 32 | 4 |
| 113 | BPPPMP | 1 | C | 9 | 1_C_9 | 1 | E | 17 | 9 | 1 |
| 114 | BMPPMP | 2 | C | 9 | 2_C_9 | 1 | E | 18 | 33 | 1 |
| 115 | AMPPMP | 1 | C | 10 | 1_C_10 | 1 | E | 19 | 9 | 2 |
| 116 | BPMPPM | 2 | C | 10 | 2_C_10 | 1 | E | 20 | 33 | 2 |
| 117 | AMMMPM | 1 | C | 11 | 1_C_11 | 1 | E | 21 | 9 | 3 |
| 118 | APPMPP | 2 | C | 11 | 2_C_11 | 1 | E | 22 | 33 | 3 |
| 119 | BPMMPM | 1 | C | 12 | 1_C_12 | 1 | E | 23 | 9 | 4 |
| 120 | AMMMMM | 2 | C | 12 | 2_C_12 | 1 | E | 24 | 33 | 4 |
| 121 | APPMPM | 4 | C | 1 | 4_C_1 | 1 | F | 1 | 79 | 1 |
| 122 | APPPPM | 3 | C | 1 | 3_C_1 | 1 | F | 2 | 55 | 1 |
| 123 | BMMMPP | 4 | C | 2 | 4_C_2 | 1 | F | 3 | 79 | 2 |
| 124 | BMMMMP | 3 | C | 2 | 3_C_2 | 1 | F | 4 | 55 | 2 |
| 125 | APMPMP | 4 | C | 3 | 4_C_3 | 1 | F | 5 | 79 | 3 |
| 126 | APMPPM | 3 | C | 3 | 3_C_3 | 1 | F | 6 | 55 | 3 |
| 127 | BMPPMM | 4 | C | 4 | 4_C_4 | 1 | F | 7 | 79 | 4 |
| 128 | BMPMMP | 3 | C | 4 | 3_C_4 | 1 | F | 8 | 55 | 4 |
| 129 | AMPMMM | 4 | C | 5 | 4_C_5 | 1 | F | 9 | 80 | 1 |
| 130 | BMPPMP | 3 | C | 5 | 3_C_5 | 1 | F | 10 | 56 | 1 |
| 131 | BPMMPP | 4 | C | 6 | 4_C_6 | 1 | F | 11 | 80 | 2 |
| 132 | AMMMPM | 3 | C | 6 | 3_C_6 | 1 | F | 12 | 56 | 2 |
| 133 | BPPPMM | 4 | C | 7 | 4_C_7 | 1 | F | 13 | 80 | 3 |

| | | | | | | | | | | |
|---|---|---|---|---|---|---|---|---|---|---|
| 134 | IPPPPP | 3 | C | 7 | 3_C_7 | 1 | F | 14 | 56 | 3 |
| 135 | AMMPPP | 4 | C | 8 | 4_C_8 | 1 | F | 15 | 80 | 4 |
| 136 | APMMMM | 3 | C | 8 | 3_C_8 | 1 | F | 16 | 56 | 4 |
| 137 | BPMMPM | 4 | C | 9 | 4_C_9 | 1 | F | 17 | 81 | 1 |
| 138 | BPPMPP | 3 | C | 9 | 3_C_9 | 1 | F | 18 | 57 | 1 |
| 139 | AMPPMP | 4 | C | 10 | 4_C_10 | 1 | F | 19 | 81 | 2 |
| 140 | APPPMP | 3 | C | 10 | 3_C_10 | 1 | F | 20 | 57 | 2 |
| 141 | BPPMMP | 4 | C | 11 | 4_C_11 | 1 | F | 21 | 81 | 3 |
| 142 | APMPPP | 3 | C | 11 | 3_C_11 | 1 | F | 22 | 57 | 3 |
| 143 | AMMPPM | 4 | C | 12 | 4_C_12 | 1 | F | 23 | 81 | 4 |
| 144 | BMPPPM | 3 | C | 12 | 3_C_12 | 1 | F | 24 | 57 | 4 |
| 145 | BMPPMP | 1 | D | 1 | 1_D_1 | 1 | G | 1 | 10 | 1 |
| 146 | APPMPM | 2 | D | 1 | 2_D_1 | 1 | G | 2 | 34 | 1 |
| 147 | AMMMPM | 1 | D | 2 | 1_D_2 | 1 | G | 3 | 10 | 2 |
| 148 | BMMMMP | 2 | D | 2 | 2_D_2 | 1 | G | 4 | 34 | 2 |
| 149 | BPMPPM | 1 | D | 3 | 1_D_3 | 1 | G | 5 | 10 | 3 |
| 150 | BMPPPM | 2 | D | 3 | 2_D_3 | 1 | G | 6 | 34 | 3 |
| 151 | APPMMP | 1 | D | 4 | 1_D_4 | 1 | G | 7 | 10 | 4 |
| 152 | APMPMP | 2 | D | 4 | 2_D_4 | 1 | G | 8 | 34 | 4 |
| 153 | APMPPM | 1 | D | 5 | 1_D_5 | 1 | G | 9 | 11 | 1 |
| 154 | BMMPPP | 2 | D | 5 | 2_D_5 | 1 | G | 10 | 35 | 1 |
| 155 | AMPMPP | 1 | D | 6 | 1_D_6 | 1 | G | 11 | 11 | 2 |
| 156 | BPPMPM | 2 | D | 6 | 2_D_6 | 1 | G | 12 | 35 | 2 |
| 157 | BPPPMP | 1 | D | 7 | 1_D_7 | 1 | G | 13 | 11 | 3 |
| 158 | AMPPMM | 2 | D | 7 | 2_D_7 | 1 | G | 14 | 35 | 3 |
| 159 | BMMMMM | 1 | D | 8 | 1_D_8 | 1 | G | 15 | 11 | 4 |
| 160 | APMMMP | 2 | D | 8 | 2_D_8 | 1 | G | 16 | 35 | 4 |
| 161 | BPMPMP | 1 | D | 9 | 1_D_9 | 1 | G | 17 | 12 | 1 |
| 162 | BMMMPP | 2 | D | 9 | 2_D_9 | 1 | G | 18 | 36 | 1 |
| 163 | BPPPPM | 1 | D | 10 | 1_D_10 | 1 | G | 19 | 12 | 2 |
| 164 | BPPPMM | 2 | D | 10 | 2_D_10 | 1 | G | 20 | 36 | 2 |
| 165 | AMPMPM | 1 | D | 11 | 1_D_11 | 1 | G | 21 | 12 | 3 |
| 166 | APMPMP | 2 | D | 11 | 2_D_11 | 1 | G | 22 | 36 | 3 |
| 167 | AMMMMP | 1 | D | 12 | 1_D_12 | 1 | G | 23 | 12 | 4 |
| 168 | APPMPM | 2 | D | 12 | 2_D_12 | 1 | G | 24 | 36 | 4 |
| 169 | BMPMPP | 4 | D | 1 | 4_D_1 | 1 | H | 1 | 82 | 1 |
| 170 | BPPPPM | 3 | D | 1 | 3_D_1 | 1 | H | 2 | 58 | 1 |
| 171 | BPMPMM | 4 | D | 2 | 4_D_2 | 1 | H | 3 | 82 | 2 |
| 172 | BPMMPP | 3 | D | 2 | 3_D_2 | 1 | H | 4 | 58 | 2 |
| 173 | APMPMM | 4 | D | 3 | 4_D_3 | 1 | H | 5 | 82 | 3 |
| 174 | AMMPMP | 3 | D | 3 | 3_D_3 | 1 | H | 6 | 58 | 3 |
| 175 | AMPMPP | 4 | D | 4 | 4_D_4 | 1 | H | 7 | 82 | 4 |
| 176 | AMPMMM | 3 | D | 4 | 3_D_4 | 1 | H | 8 | 58 | 4 |
| 177 | AMPMPP | 4 | D | 5 | 4_D_5 | 1 | H | 9 | 83 | 1 |
| 178 | AMPMMP | 3 | D | 5 | 3_D_5 | 1 | H | 10 | 59 | 1 |

| | | | | | | | | | | |
|---|---|---|---|---|---|---|---|---|---|---|
| 179 | BPPPMP | 4 | D | 6 | 4_D_6 | 1 | H | 11 | 83 | 2 |
| 180 | BPMMPM | 3 | D | 6 | 3_D_6 | 1 | H | 12 | 59 | 2 |
| 181 | BMMPMM | 4 | D | 7 | 4_D_7 | 1 | H | 13 | 83 | 3 |
| 182 | AMMPPM | 3 | D | 7 | 3_D_7 | 1 | H | 14 | 59 | 3 |
| 183 | APMMPM | 4 | D | 8 | 4_D_8 | 1 | H | 15 | 83 | 4 |
| 184 | BPPPMP | 3 | D | 8 | 3_D_8 | 1 | H | 16 | 59 | 4 |
| 185 | BPMPMP | 4 | D | 9 | 4_D_9 | 1 | H | 17 | 84 | 1 |
| 186 | AMPPPM | 3 | D | 9 | 3_D_9 | 1 | H | 18 | 60 | 1 |
| 187 | APMPPP | 4 | D | 10 | 4_D_10 | 1 | H | 19 | 84 | 2 |
| 188 | AMMMMP | 3 | D | 10 | 3_D_10 | 1 | H | 20 | 60 | 2 |
| 189 | AMPMPM | 4 | D | 11 | 4_D_11 | 1 | H | 21 | 84 | 3 |
| 190 | BPPMMM | 3 | D | 11 | 3_D_11 | 1 | H | 22 | 60 | 3 |
| 191 | BMPMMM | 4 | D | 12 | 4_D_12 | 1 | H | 23 | 84 | 4 |
| 192 | BPMPPP | 3 | D | 12 | 3_D_12 | 1 | H | 24 | 60 | 4 |
| 193 | BMMPPP | 1 | E | 1 | 1_E_1 | 1 | I | 1 | 13 | 1 |
| 194 | APMMPP | 2 | E | 1 | 2_E_1 | 1 | I | 2 | 37 | 1 |
| 195 | BPPMMM | 1 | E | 2 | 1_E_2 | 1 | I | 3 | 13 | 2 |
| 196 | BMMMMP | 2 | E | 2 | 2_E_2 | 1 | I | 4 | 37 | 2 |
| 197 | AMMMMP | 1 | E | 3 | 1_E_3 | 1 | I | 5 | 13 | 3 |
| 198 | BMPPPM | 2 | E | 3 | 2_E_3 | 1 | I | 6 | 37 | 3 |
| 199 | APPPPM | 1 | E | 4 | 1_E_4 | 1 | I | 7 | 13 | 4 |
| 200 | APPPMM | 2 | E | 4 | 2_E_4 | 1 | I | 8 | 37 | 4 |
| 201 | IPPPPP | 1 | E | 5 | 1_E_5 | 1 | I | 9 | 14 | 1 |
| 202 | AMPMPM | 2 | E | 5 | 2_E_5 | 1 | I | 10 | 38 | 1 |
| 203 | BMMMMM | 1 | E | 6 | 1_E_6 | 1 | I | 11 | 14 | 2 |
| 204 | BPMPMP | 2 | E | 6 | 2_E_6 | 1 | I | 12 | 38 | 2 |
| 205 | AMPMPP | 1 | E | 7 | 1_E_7 | 1 | I | 13 | 14 | 3 |
| 206 | BPPPPM | 2 | E | 7 | 2_E_7 | 1 | I | 14 | 38 | 3 |
| 207 | APMPMM | 1 | E | 8 | 1_E_8 | 1 | I | 15 | 14 | 4 |
| 208 | AMMMMP | 2 | E | 8 | 2_E_8 | 1 | I | 16 | 38 | 4 |
| 209 | APMPMP | 1 | E | 9 | 1_E_9 | 1 | I | 17 | 15 | 1 |
| 210 | IPPPPP | 2 | E | 9 | 2_E_9 | 1 | I | 18 | 39 | 1 |
| 211 | BPPMMP | 1 | E | 10 | 1_E_10 | 1 | I | 19 | 15 | 2 |
| 212 | AMMMPM | 2 | E | 10 | 2_E_10 | 1 | I | 20 | 39 | 2 |
| 213 | BMMMPM | 1 | E | 11 | 1_E_11 | 1 | I | 21 | 15 | 3 |
| 214 | BPPPMP | 2 | E | 11 | 2_E_11 | 1 | I | 22 | 39 | 3 |
| 215 | AMPPPP | 1 | E | 12 | 1_E_12 | 1 | I | 23 | 15 | 4 |
| 216 | BMMMMM | 2 | E | 12 | 2_E_12 | 1 | I | 24 | 39 | 4 |
| 217 | AMMPMP | 4 | E | 1 | 4_E_1 | 1 | J | 1 | 85 | 1 |
| 218 | BMPPMP | 3 | E | 1 | 3_E_1 | 1 | J | 2 | 61 | 1 |
| 219 | BPPMPM | 4 | E | 2 | 4_E_2 | 1 | J | 3 | 85 | 2 |
| 220 | AMMPPP | 3 | E | 2 | 3_E_2 | 1 | J | 4 | 61 | 2 |
| 221 | BMMPMM | 4 | E | 3 | 4_E_3 | 1 | J | 5 | 86 | 1 |
| 222 | APMMPM | 3 | E | 3 | 3_E_3 | 1 | J | 6 | 61 | 3 |
| 223 | AMMMMM | 4 | E | 4 | 4_E_4 | 1 | J | 7 | 86 | 2 |

| | | | | | | | | | | |
|---|---|---|---|---|---|---|---|---|---|---|
| 224 | BPPMMM | 3 | E | 4 | 3_E_4 | 1 | J | 8 | 61 | 4 |
| 225 | BPMMMP | 4 | E | 5 | 4_E_5 | 1 | J | 9 | 87 | 1 |
| 226 | AMPMMP | 3 | E | 5 | 3_E_5 | 1 | J | 10 | 62 | 1 |
| 227 | BMPMPM | 4 | E | 6 | 4_E_6 | 1 | J | 11 | 87 | 2 |
| 228 | AMMMPM | 3 | E | 6 | 3_E_6 | 1 | J | 12 | 62 | 2 |
| 229 | AMPMMM | 4 | E | 7 | 4_E_7 | 1 | J | 13 | 88 | 1 |
| 230 | BPMPPM | 3 | E | 7 | 3_E_7 | 1 | J | 14 | 62 | 3 |
| 231 | BPPMMM | 4 | E | 8 | 4_E_8 | 1 | J | 15 | 88 | 2 |
| 232 | APPPMP | 3 | E | 8 | 3_E_8 | 1 | J | 16 | 62 | 4 |
| 233 | AMMPMM | 4 | E | 9 | 4_E_9 | 1 | J | 17 | 89 | 1 |
| 234 | BPPPPM | 3 | E | 9 | 3_E_9 | 1 | J | 18 | 63 | 1 |
| 235 | BPPMMP | 4 | E | 10 | 4_E_10 | 1 | J | 19 | 89 | 2 |
| 236 | AMMMMP | 3 | E | 10 | 3_E_10 | 1 | J | 20 | 63 | 2 |
| 237 | APPMPP | 4 | E | 11 | 4_E_11 | 1 | J | 21 | 90 | 1 |
| 238 | APMMMP | 3 | E | 11 | 3_E_11 | 1 | J | 22 | 63 | 3 |
| 239 | BPPMMP | 4 | E | 12 | 4_E_12 | 1 | J | 23 | 90 | 2 |
| 240 | BMPPPM | 3 | E | 12 | 3_E_12 | 1 | J | 24 | 63 | 4 |
| 241 | BMPPPP | 1 | F | 1 | 1_F_1 | 1 | K | 1 | 16 | 1 |
| 242 | BPPPMM | 2 | F | 1 | 2_F_1 | 1 | K | 2 | 40 | 1 |
| 243 | BMMMPM | 1 | F | 2 | 1_F_2 | 1 | K | 3 | 16 | 2 |
| 244 | APMPPP | 2 | F | 2 | 2_F_2 | 1 | K | 4 | 40 | 2 |
| 245 | APPMMP | 1 | F | 3 | 1_F_3 | 1 | K | 5 | 16 | 3 |
| 246 | AMPMPM | 2 | F | 3 | 2_F_3 | 1 | K | 6 | 40 | 3 |
| 247 | APMPMM | 1 | F | 4 | 1_F_4 | 1 | K | 7 | 16 | 4 |
| 248 | BMMMMP | 2 | F | 4 | 2_F_4 | 1 | K | 8 | 40 | 4 |
| 249 | BMPMPM | 1 | F | 5 | 1_F_5 | 1 | K | 9 | 17 | 1 |
| 250 | BPMPMP | 2 | F | 5 | 2_F_5 | 1 | K | 10 | 41 | 1 |
| 251 | BPMPMP | 1 | F | 6 | 1_F_6 | 1 | K | 11 | 17 | 2 |
| 252 | BMPMMM | 2 | F | 6 | 2_F_6 | 1 | K | 12 | 41 | 2 |
| 253 | APPPPM | 1 | F | 7 | 1_F_7 | 1 | K | 13 | 17 | 3 |
| 254 | AMMPPP | 2 | F | 7 | 2_F_7 | 1 | K | 14 | 41 | 3 |
| 255 | AMMMMP | 1 | F | 8 | 1_F_8 | 1 | K | 15 | 17 | 4 |
| 256 | APPMPM | 2 | F | 8 | 2_F_8 | 1 | K | 16 | 41 | 4 |
| 257 | IPPPPP | 1 | F | 9 | 1_F_9 | 1 | K | 17 | 18 | 1 |
| 258 | APMPPP | 2 | F | 9 | 2_F_9 | 1 | K | 18 | 42 | 1 |
| 259 | IPPPPP | 1 | F | 10 | 1_F_10 | 1 | K | 19 | 18 | 2 |
| 260 | BMPMMM | 2 | F | 10 | 2_F_10 | 1 | K | 20 | 42 | 2 |
| 261 | IPPPPP | 1 | F | 11 | 1_F_11 | 1 | K | 21 | 18 | 3 |
| 262 | APPPPM | 2 | F | 11 | 2_F_11 | 1 | K | 22 | 42 | 3 |
| 263 | IPPPPP | 1 | F | 12 | 1_F_12 | 1 | K | 23 | 18 | 4 |
| 264 | BMMMMP | 2 | F | 12 | 2_F_12 | 1 | K | 24 | 42 | 4 |
| 265 | BPPMPP | 4 | F | 1 | 4_F_1 | 1 | L | 1 | 91 | 1 |
| 266 | APPMPM | 3 | F | 1 | 3_F_1 | 1 | L | 2 | 64 | 1 |
| 267 | APMMMM | 4 | F | 2 | 4_F_2 | 1 | L | 3 | 91 | 2 |
| 268 | IPMMMP | 3 | F | 2 | 3_F_2 | 1 | L | 4 | 64 | 2 |

| | | | | | | | | | | |
|---|---|---|---|---|---|---|---|---|---|---|
| 269 | AMMPMM | 4 | F | 3 | 4_F_3 | 1 | L | 5 | 91 | 3 |
| 270 | BPPPMP | 3 | F | 3 | 3_F_3 | 1 | L | 6 | 64 | 3 |
| 271 | IPPPPP | 4 | F | 4 | 4_F_4 | 1 | L | 7 | 92 | 1 |
| 272 | APMPPM | 3 | F | 4 | 3_F_4 | 1 | L | 8 | 64 | 4 |
| 273 | IPPPPP | 4 | F | 5 | 4_F_5 | 1 | L | 9 | 92 | 2 |
| 274 | APPPPM | 3 | F | 5 | 3_F_5 | 1 | L | 10 | 65 | 1 |
| 275 | IPPPPP | 4 | F | 6 | 4_F_6 | 1 | L | 11 | 92 | 3 |
| 276 | BMPMPM | 3 | F | 6 | 3_F_6 | 1 | L | 12 | 65 | 2 |
| 277 | APPMPP | 4 | F | 7 | 4_F_7 | 1 | L | 13 | 93 | 1 |
| 278 | BPMMMP | 3 | F | 7 | 3_F_7 | 1 | L | 14 | 65 | 3 |
| 279 | BMPPMP | 4 | F | 8 | 4_F_8 | 1 | L | 15 | 93 | 2 |
| 280 | AMMPMP | 3 | F | 8 | 3_F_8 | 1 | L | 16 | 65 | 4 |
| 281 | APMPMM | 4 | F | 9 | 4_F_9 | 1 | L | 17 | 93 | 3 |
| 282 | IPPPPP | 3 | F | 9 | 3_F_9 | 1 | L | 18 | 66 | 1 |
| 283 | AMPMPP | 4 | F | 10 | 4_F_10 | 1 | L | 19 | 94 | 1 |
| 284 | AMMMPM | 3 | F | 10 | 3_F_10 | 1 | L | 20 | 66 | 2 |
| 285 | BPPPMM | 4 | F | 11 | 4_F_11 | 1 | L | 21 | 94 | 2 |
| 286 | AMPMMM | 3 | F | 11 | 3_F_11 | 1 | L | 22 | 66 | 3 |
| 287 | APMPMM | 4 | F | 12 | 4_F_12 | 1 | L | 23 | 94 | 3 |
| 288 | BPMPMP | 3 | F | 12 | 3_F_12 | 1 | L | 24 | 66 | 4 |
| 289 | BPMMPM | 1 | G | 1 | 1_G_1 | 1 | M | 1 | 19 | 1 |
| 290 | BPPPMM | 2 | G | 1 | 2_G_1 | 1 | M | 2 | 43 | 1 |
| 291 | APPPMP | 1 | G | 2 | 1_G_2 | 1 | M | 3 | 19 | 2 |
| 292 | APMMPP | 2 | G | 2 | 2_G_2 | 1 | M | 4 | 43 | 2 |
| 293 | BMPMMP | 1 | G | 3 | 1_G_3 | 1 | M | 5 | 19 | 3 |
| 294 | AMPMMM | 2 | G | 3 | 2_G_3 | 1 | M | 6 | 43 | 3 |
| 295 | AMMPPM | 1 | G | 4 | 1_G_4 | 1 | M | 7 | 19 | 4 |
| 296 | BMMPPP | 2 | G | 4 | 2_G_4 | 1 | M | 8 | 43 | 4 |
| 297 | AMMMPM | 1 | G | 5 | 1_G_5 | 1 | M | 9 | 20 | 1 |
| 298 | APPMMP | 2 | G | 5 | 2_G_5 | 1 | M | 10 | 44 | 1 |
| 299 | BPPMMP | 1 | G | 6 | 1_G_6 | 1 | M | 11 | 20 | 2 |
| 300 | BPPMMP | 2 | G | 6 | 2_G_6 | 1 | M | 12 | 44 | 2 |
| 301 | BPMPPM | 1 | G | 7 | 1_G_7 | 1 | M | 13 | 20 | 3 |
| 302 | BMMPPM | 2 | G | 7 | 2_G_7 | 1 | M | 14 | 44 | 3 |
| 303 | AMPPMP | 1 | G | 8 | 1_G_8 | 1 | M | 15 | 20 | 4 |
| 304 | AMMPPM | 2 | G | 8 | 2_G_8 | 1 | M | 16 | 44 | 4 |
| 305 | BMMPMP | 1 | G | 9 | 1_G_9 | 1 | M | 17 | 21 | 1 |
| 306 | APMPMP | 2 | G | 9 | 2_G_9 | 1 | M | 18 | 45 | 1 |
| 307 | AMMMMP | 1 | G | 10 | 1_G_10 | 1 | M | 19 | 21 | 2 |
| 308 | BPPMPM | 2 | G | 10 | 2_G_10 | 1 | M | 20 | 45 | 2 |
| 309 | APPMPM | 1 | G | 11 | 1_G_11 | 1 | M | 21 | 21 | 3 |
| 310 | AMPMPP | 2 | G | 11 | 2_G_11 | 1 | M | 22 | 45 | 3 |
| 311 | BPPPPM | 1 | G | 12 | 1_G_12 | 1 | M | 23 | 21 | 4 |
| 312 | BMMPMM | 2 | G | 12 | 2_G_12 | 1 | M | 24 | 45 | 4 |
| 313 | IPPPPP | 4 | G | 1 | 4_G_1 | 1 | N | 1 | 95 | 1 |

| | | | | | | | | | | |
|---|---|---|---|---|---|---|---|---|---|---|
| 314 | IPPPPP | 3 | G | 1 | 3_G_1 | 1 | N | 2 | 67 | 1 |
| 315 | BPMMPM | 4 | G | 2 | 4_G_2 | 1 | N | 3 | 95 | 2 |
| 316 | IMPPPM | 3 | G | 2 | 3_G_2 | 1 | N | 4 | 67 | 2 |
| 317 | AMMMMM | 4 | G | 3 | 4_G_3 | 1 | N | 5 | 95 | 3 |
| 318 | BPPPMP | 3 | G | 3 | 3_G_3 | 1 | N | 6 | 67 | 3 |
| 319 | AMMPMM | 4 | G | 4 | 4_G_4 | 1 | N | 7 | 96 | 1 |
| 320 | APMPPP | 3 | G | 4 | 3_G_4 | 1 | N | 8 | 67 | 4 |
| 321 | BMMPPM | 4 | G | 5 | 4_G_5 | 1 | N | 9 | 96 | 2 |
| 322 | AMMMMP | 3 | G | 5 | 3_G_5 | 1 | N | 10 | 68 | 1 |
| 323 | BPPMPP | 4 | G | 6 | 4_G_6 | 1 | N | 11 | 96 | 3 |
| 324 | APMPPP | 3 | G | 6 | 3_G_6 | 1 | N | 12 | 68 | 2 |
| 325 | APMMMM | 4 | G | 7 | 4_G_7 | 1 | N | 13 | 97 | 1 |
| 326 | BMPMPM | 3 | G | 7 | 3_G_7 | 1 | N | 14 | 68 | 3 |
| 327 | AMPPPP | 4 | G | 8 | 4_G_8 | 1 | N | 15 | 97 | 2 |
| 328 | BPPPMM | 3 | G | 8 | 3_G_8 | 1 | N | 16 | 68 | 4 |
| 329 | BMMPMM | 4 | G | 9 | 4_G_9 | 1 | N | 17 | 97 | 3 |
| 330 | BMPPMM | 3 | G | 9 | 3_G_9 | 1 | N | 18 | 69 | 1 |
| 331 | BPPPPM | 4 | G | 10 | 4_G_10 | 1 | N | 19 | 98 | 1 |
| 332 | IPPPPP | 3 | G | 10 | 3_G_10 | 1 | N | 20 | 69 | 2 |
| 333 | AMPMPP | 4 | G | 11 | 4_G_11 | 1 | N | 21 | 98 | 2 |
| 334 | APMMMM | 3 | G | 11 | 3_G_11 | 1 | N | 22 | 69 | 3 |
| 335 | APMMMP | 4 | G | 12 | 4_G_12 | 1 | N | 23 | 98 | 3 |
| 336 | AMMMPP | 3 | G | 12 | 3_G_12 | 1 | N | 24 | 69 | 4 |
| 337 | BMPPMM | 1 | H | 1 | 1_H_1 | 1 | O | 1 | 22 | 1 |
| 338 | BMPMMP | 2 | H | 1 | 2_H_1 | 1 | O | 2 | 46 | 1 |
| 339 | APMMMM | 1 | H | 2 | 1_H_2 | 1 | O | 3 | 22 | 2 |
| 340 | APPPPM | 2 | H | 2 | 2_H_2 | 1 | O | 4 | 46 | 2 |
| 341 | APMMPP | 1 | H | 3 | 1_H_3 | 1 | O | 5 | 22 | 3 |
| 342 | BPMPMP | 2 | H | 3 | 2_H_3 | 1 | O | 6 | 46 | 3 |
| 343 | BMPPPP | 1 | H | 4 | 1_H_4 | 1 | O | 7 | 22 | 4 |
| 344 | AMMMPM | 2 | H | 4 | 2_H_4 | 1 | O | 8 | 46 | 4 |
| 345 | AMMMPM | 1 | H | 5 | 1_H_5 | 1 | O | 9 | 23 | 1 |
| 346 | BMPPPM | 2 | H | 5 | 2_H_5 | 1 | O | 10 | 47 | 1 |
| 347 | BPPPMM | 1 | H | 6 | 1_H_6 | 1 | O | 11 | 23 | 2 |
| 348 | BMMPPP | 2 | H | 6 | 2_H_6 | 1 | O | 12 | 47 | 2 |
| 349 | AMPMMP | 1 | H | 7 | 1_H_7 | 1 | O | 13 | 23 | 3 |
| 350 | APPMMM | 2 | H | 7 | 2_H_7 | 1 | O | 14 | 47 | 3 |
| 351 | BPMPPP | 1 | H | 8 | 1_H_8 | 1 | O | 15 | 23 | 4 |
| 352 | APMMMP | 2 | H | 8 | 2_H_8 | 1 | O | 16 | 47 | 4 |
| 353 | BPMMMP | 1 | H | 9 | 1_H_9 | 1 | O | 17 | 24 | 1 |
| 354 | APMMPM | 2 | H | 9 | 2_H_9 | 1 | O | 18 | 48 | 1 |
| 355 | BPPMPM | 1 | H | 10 | 1_H_10 | 1 | O | 19 | 24 | 2 |
| 356 | BMMMPM | 2 | H | 10 | 2_H_10 | 1 | O | 20 | 48 | 2 |
| 357 | AMMPPP | 1 | H | 11 | 1_H_11 | 1 | O | 21 | 24 | 3 |
| 358 | BPPPMP | 2 | H | 11 | 2_H_11 | 1 | O | 22 | 48 | 3 |

| | | | | | | | | | | |
|---|---|---|---|---|---|---|---|---|---|---|
| 359 | AMPPMM | 1 | H | 12 | 1_H_12 | 1 | O | 23 | 24 | 4 |
| 360 | AMPPMP | 2 | H | 12 | 2_H_12 | 1 | O | 24 | 48 | 4 |
| 361 | APMMPM | 4 | H | 1 | 4_H_1 | 1 | P | 1 | 99 | 1 |
| 362 | AMPPMM | 3 | H | 1 | 3_H_1 | 1 | P | 2 | 70 | 1 |
| 363 | BPMPMP | 4 | H | 2 | 4_H_2 | 1 | P | 3 | 99 | 2 |
| 364 | APMMPP | 3 | H | 2 | 3_H_2 | 1 | P | 4 | 70 | 2 |
| 365 | BMPPMP | 4 | H | 3 | 4_H_3 | 1 | P | 5 | 99 | 3 |
| 366 | BMPMMP | 3 | H | 3 | 3_H_3 | 1 | P | 6 | 70 | 3 |
| 367 | BPMPMM | 4 | H | 4 | 4_H_4 | 1 | P | 7 | 100 | 1 |
| 368 | BPMPPM | 3 | H | 4 | 3_H_4 | 1 | P | 8 | 70 | 4 |
| 369 | BMMMPM | 4 | H | 5 | 4_H_5 | 1 | P | 9 | 100 | 2 |
| 370 | BMPMPM | 3 | H | 5 | 3_H_5 | 1 | P | 10 | 71 | 1 |
| 371 | APPPMP | 4 | H | 6 | 4_H_6 | 1 | P | 11 | 100 | 3 |
| 372 | BPPPPM | 3 | H | 6 | 3_H_6 | 1 | P | 12 | 71 | 2 |
| 373 | BPPMPP | 4 | H | 7 | 4_H_7 | 1 | P | 13 | 101 | 1 |
| 374 | APMPMP | 3 | H | 7 | 3_H_7 | 1 | P | 14 | 71 | 3 |
| 375 | IPPPPP | 4 | H | 8 | 4_H_8 | 1 | P | 15 | 101 | 2 |
| 376 | AMMMMP | 3 | H | 8 | 3_H_8 | 1 | P | 16 | 71 | 4 |
| 377 | AMMPMM | 4 | H | 9 | 4_H_9 | 1 | P | 17 | 101 | 3 |
| 378 | BMPMPP | 3 | H | 9 | 3_H_9 | 1 | P | 18 | 72 | 1 |
| 379 | APMMMM | 4 | H | 10 | 4_H_10 | 1 | P | 19 | 102 | 1 |
| 380 | AMPPMM | 3 | H | 10 | 3_H_10 | 1 | P | 20 | 72 | 2 |
| 381 | BMPMMP | 4 | H | 11 | 4_H_11 | 1 | P | 21 | 102 | 2 |
| 382 | APMMMM | 3 | H | 11 | 3_H_11 | 1 | P | 22 | 72 | 3 |
| 383 | IPPPPP | 4 | H | 12 | 4_H_12 | 1 | P | 23 | 102 | 3 |
| 384 | BPMPPP | 3 | H | 12 | 3_H_12 | 1 | P | 24 | 72 | 4 |

| Table S3. | | | | | | |
|---|---|---|---|---|---|---|
| **Aux Panel 1** | **tetrads** | **4 viable** | **3 viable** | **2 viable** | **1 viable** | **total spore viability** |
| | 104 | 76 | 20 | 6 | 2 | |
| | % | 73.1 | 19.2 | 5.8 | 1.9 | 0.908653846 |
| **Aux Panel 2** | 113 | 92 | 15 | 6 | 0 | |
| | % | 81.4 | 13.3 | 5.3 | 0.0 | 0.940265487 |
| **increase** | | **0.1135** | **-0.3073** | | | **0.03478953** |

**Table S4.**

| YNB | (g/L) |
|---|---|
| *Inorganic Salts* | |
| Calcium chloride | 0.1 |
| Magnesium sulfate | 0.05 |
| Sodium chloride | 0.1 |
| PABA | 0.0002 |
| Ferric chloride | 0.0002 |
| Manganese sulfate | 0.0004 |
| Biotin | 0.000002 |
| Sodium molybdate | 0.0002 |
| Pantothenate calcium | 0.0004 |
| Zinc sulfate | 0.0004 |
| Boric acid | 0.0005 |
| Potassium phosphate | 0.5 |
| Copper sulfate | 0.00004 |
| Potassium iodide | 0.0001 |

| | |
|---|---|
| *Vitamins* | |
| Folic acid | 0.000002 |
| Inositol | 0.002 |
| Niacin | 0.0004 |
| Pyridoxine | 0.0004 |
| Riboflavin | 0.0002 |
| Thiamine | 0.0004 |

| Amino Acid Mixture | (g/L) | recipe (gm) |
|---|---|---|
| Adenine | 0.0183 | 0.5 |
| Alanine | 0.0734 | 2 |
| Arginine | 0.0734 | 2 |
| Asparagine | 0.0734 | 2 |
| Aspartic Acid | 0.0734 | 2 |
| Cysteine | 0.0734 | 2 |
| Glutamic Acid | 0.0734 | 2 |
| Glycine | 0.0734 | 2 |
| Histidine | 0.0734 | 2 |
| Inositol | 0.0250 | 0.7 |
| Isoleucine | 0.0734 | 2 |
| Leucine | 0.1468 | 4 |
| Lysine | 0.0734 | 2 |
| Methionine | 0.0734 | 2 |
| Phenylalanine | 0.0734 | 2 |
| Proline | 0.0734 | 2 |
| Serine | 0.0734 | 2 |
| Threonine | 0.0734 | 2 |
| Tryptophan | 0.0734 | 2 |
| Tyrosin | 0.0734 | 2 |
| Uracil | 0.0734 | 2 |
| Valine | 0.0734 | 2 |
| Glutamine | 0.0734 | 2 |

** PABA omitted

| Other (variable) | |
|---|---|
| Glucose | 20 |
| Ammonium Sulfate | 0.5 |

| Table S5. | | | | | | | | | | | |
|---|---|---|---|---|---|---|---|---|---|---|---|
| **Gene** | **L_A** | **L_A_se** | **L_NA** | **L_NA_se** | **pH_A** | **pH_A_se** | **pH_NA** | **pH_NA_se** | **LeuLysMet** | **His** | **Ura** |
| MMMMM | 14.84 | 0.18 | 16.13 | 0.22 | 3.32 | 0.02 | 3.64 | 0.02 | MMM | M | M |
| MMMMP | 15.14 | 0.20 | 15.81 | 0.17 | 3.34 | 0.01 | 3.65 | 0.01 | MMM | M | P |
| PMMMM | 14.66 | 0.24 | 15.76 | 0.27 | 3.32 | 0.02 | 3.65 | 0.01 | MMM | P | M |
| PMMMP | 15.07 | 0.22 | 15.55 | 0.25 | 3.33 | 0.02 | 3.65 | 0.01 | MMM | P | P |
| MMMPM | 16.56 | 0.21 | 17.05 | 0.36 | 3.21 | 0.01 | 3.56 | 0.00 | MMP | M | M |
| MMMPP | 16.94 | 0.34 | 16.45 | 0.48 | 3.30 | 0.03 | 3.70 | 0.02 | MMP | M | P |
| PMMPM | 16.68 | 0.30 | 16.17 | 0.30 | 3.18 | 0.01 | 3.58 | 0.01 | MMP | P | M |
| PMMPP | 18.30 | 0.40 | 16.82 | 0.43 | 3.22 | 0.02 | 3.62 | 0.01 | MMP | P | P |
| MMPMM | 16.13 | 0.21 | 15.56 | 0.18 | 3.24 | 0.02 | 3.97 | 0.02 | MPM | M | M |
| MMPMP | 16.02 | 0.25 | 15.61 | 0.21 | 3.29 | 0.02 | 4.03 | 0.01 | MPM | M | P |
| PMPMM | 15.82 | 0.19 | 15.29 | 0.18 | 3.22 | 0.01 | 3.96 | 0.01 | MPM | P | M |
| PMPMP | 15.79 | 0.13 | 15.34 | 0.16 | 3.29 | 0.01 | 4.06 | 0.01 | MPM | P | P |
| MMPPM | 15.95 | 0.22 | 15.28 | 0.25 | 3.07 | 0.01 | 3.90 | 0.01 | MPP | M | M |
| MMPPP | 15.35 | 0.20 | 15.29 | 0.21 | 3.08 | 0.01 | 3.98 | 0.01 | MPP | M | P |
| PMPPM | 15.31 | 0.13 | 15.61 | 0.26 | 3.08 | 0.01 | 3.87 | 0.01 | MPP | P | M |
| PMPPP | 15.32 | 0.19 | 14.96 | 0.21 | 3.09 | 0.01 | 3.96 | 0.02 | MPP | P | P |
| MPMMM | 16.38 | 0.37 | 16.94 | 0.26 | 3.42 | 0.02 | 3.64 | 0.02 | PMM | M | M |
| MPMMP | 15.20 | 0.25 | 15.93 | 0.26 | 3.38 | 0.02 | 3.66 | 0.01 | PMM | M | P |
| PPMMM | 15.11 | 0.21 | 15.44 | 0.23 | 3.30 | 0.02 | 3.59 | 0.01 | PMM | P | M |
| PPMMP | 14.94 | 0.17 | 16.40 | 0.27 | 3.43 | 0.02 | 3.66 | 0.01 | PMM | P | P |
| MPMPM | 16.36 | 0.21 | 16.48 | 0.28 | 3.24 | 0.01 | 3.58 | 0.01 | PMP | M | M |
| MPMPP | 16.62 | 0.18 | 16.82 | 0.23 | 3.27 | 0.01 | 3.64 | 0.01 | PMP | M | P |
| PPMPM | 16.72 | 0.17 | 15.76 | 0.24 | 3.23 | 0.01 | 3.60 | 0.01 | PMP | P | M |
| PPMPP | 16.60 | 0.26 | 16.69 | 0.33 | 3.26 | 0.01 | 3.62 | 0.01 | PMP | P | P |
| MPPMM | 13.97 | 0.16 | 13.76 | 0.28 | 3.22 | 0.02 | 3.77 | 0.01 | PPM | M | M |
| MPPMP | 14.09 | 0.16 | 15.20 | 0.21 | 3.17 | 0.01 | 3.82 | 0.01 | PPM | M | P |
| PPPMM | 14.48 | 0.15 | 15.07 | 0.18 | 3.16 | 0.01 | 3.77 | 0.01 | PPM | P | M |
| PPPMP | 13.69 | 0.13 | 15.40 | 0.22 | 3.16 | 0.01 | 3.80 | 0.01 | PPM | P | P |
| MPPPM | 14.42 | 0.19 | 14.69 | 0.22 | 3.26 | 0.01 | 3.75 | 0.02 | PPP | M | M |
| MPPPP | 14.63 | 0.32 | 14.30 | 0.24 | 3.22 | 0.02 | 3.85 | 0.04 | PPP | M | P |
| PPPPM | 14.90 | 0.22 | 14.90 | 0.14 | 3.27 | 0.01 | 3.72 | 0.01 | PPP | P | M |
| PPPPP | 14.65 | 0.16 | 14.74 | 0.17 | 3.20 | 0.01 | 3.74 | 0.01 | PrPrPr | Pr | Pr |

| **Table S6.** | | | | | | | | | | | | | | | | | |
|---|---|---|---|---|---|---|---|---|---|---|---|---|---|---|---|---|---|
| | *his3* | *leu2* | *lys2* | *met17* | *ura3* | [leucine] | [lysine] | [methionine] | [uracil] | [ammonium sulfate] | [glutamine] | [glucose] | HU | rapamycin | *TOR1* | [methionine and cysteine] | [threonine] |
| *his3* | | | | | | | | | | | | | | | | | |
| *leu2* | | | | | | | | | | | | | | | | | |
| *lys2* | | x | | | | | | | | | | | | | | | |
| *met17* | | x | x | | | | | | | | | | | | | | |
| *ura3* | | | | | | | | | | | | | | | | | |
| [leucine] | | x | | | | | | | | | | | | | | | |
| [lysine] | | x | x | | | | | | | | | | | | | | |
| [methionine] | | | | x | | | | | | | | | | | | | |
| [uracil] | | | | | x | | | | | | | | | | | | |
| [ammonium sulfate] | | x | | x | | | | | | | | | | | | | |
| [glutamine] | | | | x | | | | | | | | | | | | | |
| [glucose] | | x | x | x | | | | | | x | | | | | | | |
| HU | | | x | x | | | | | | | | | | | | | |
| rapamycin | | | x | x | | | | | | | | | | | | | |
| *TOR1* | | | | x | | | | | | x | x | | | | | | |
| [methionine and cysteine] | | | x | x | | | | | | | | | x | | | | |
| [threonine] | | | x | x | | | | | | | | | | | | | |
