## Supplementary material for "High-resolution yeast quiescence profiling in human-like media reveals interacting influences of auxotrophy and nutrient availability": Online Resource 3

Online Resource 3. Quiescence profiles, with K data, which are complementary ("_K") to main figures that display L data. See Online Resource 1 ("K plots") for legends.

Fig. 3_K

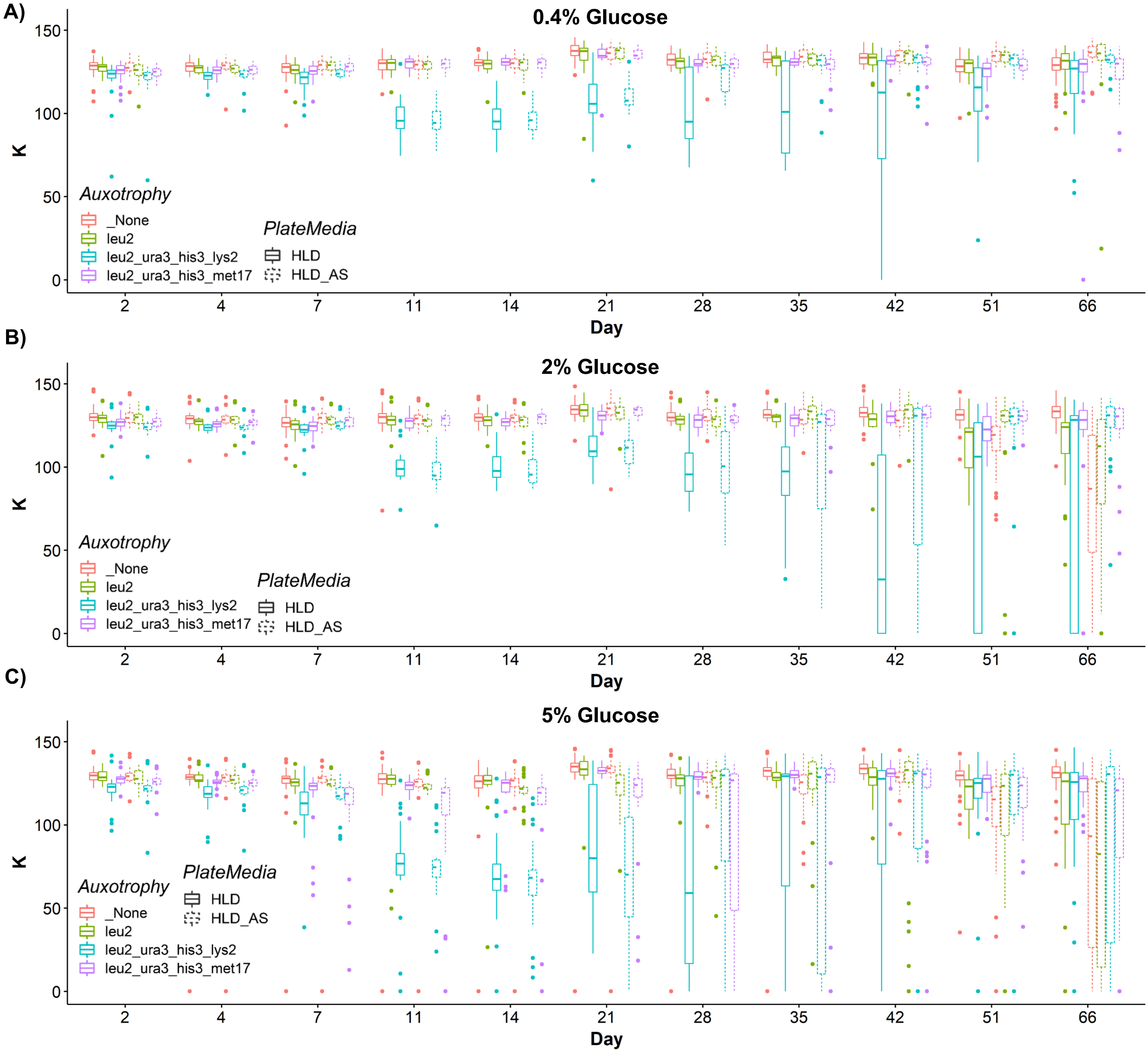
A)
0.4% Glucose
K
150
100
50
0
Auxotrophy
_None
leu2
leu2_ura3_his3_lys2
leu2_ura3_his3_met17
PlateMedia
HLD
HLD_AS
2
4
7
11
14
21
28
35
42
51
66
Day
B)
2% Glucose
C)
5% Glucose

### Fig. S2_K

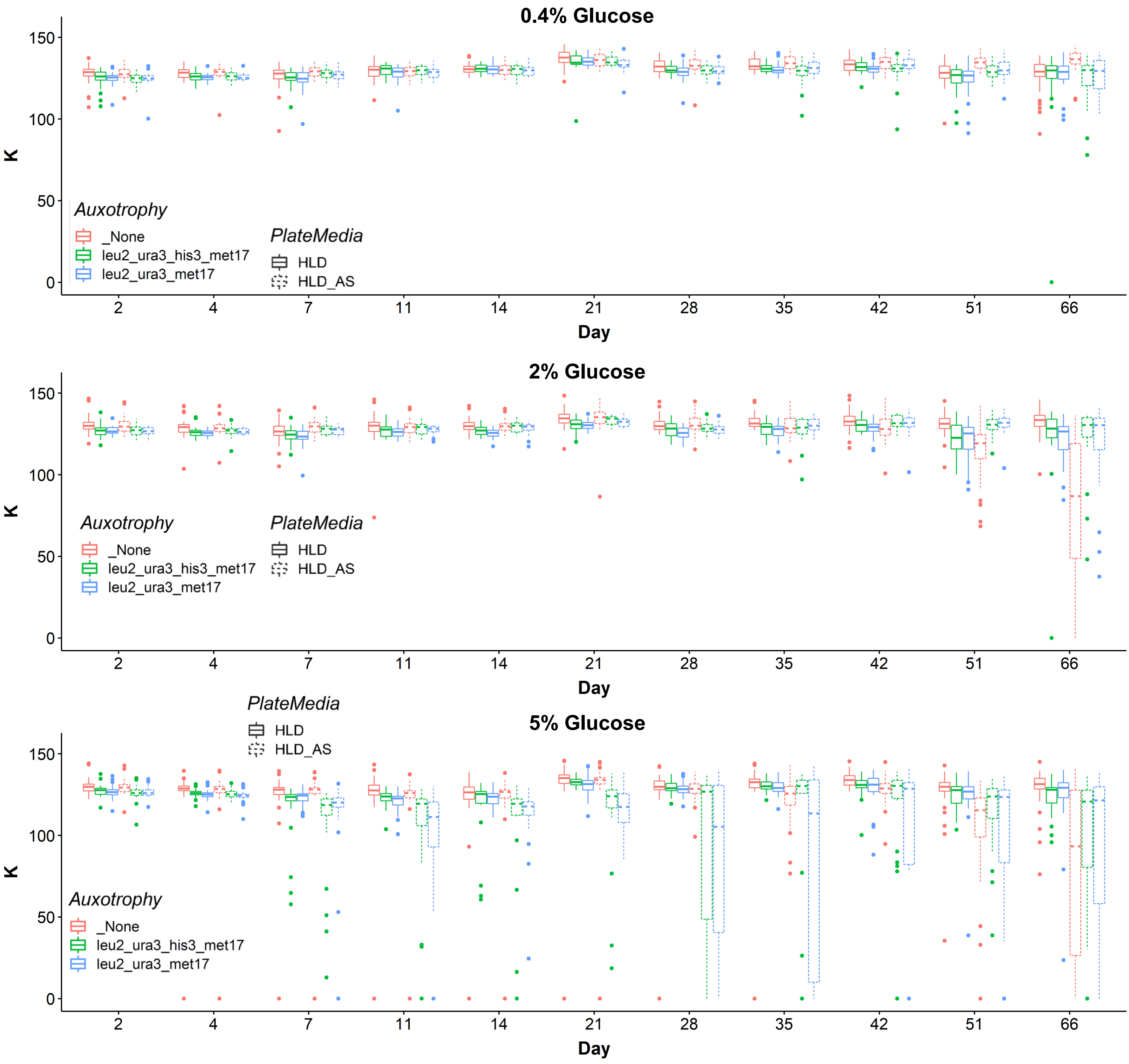
0.4% Glucose
2% Glucose
5% Glucose
K
Day
0
50
100
150
2
4
7
11
14
21
28
35
42
51
66
Auxotrophy
_None
leu2_ura3_his3_met17
leu2_ura3_met17
PlateMedia
HLD
HLD_AS

Fig. S3_K

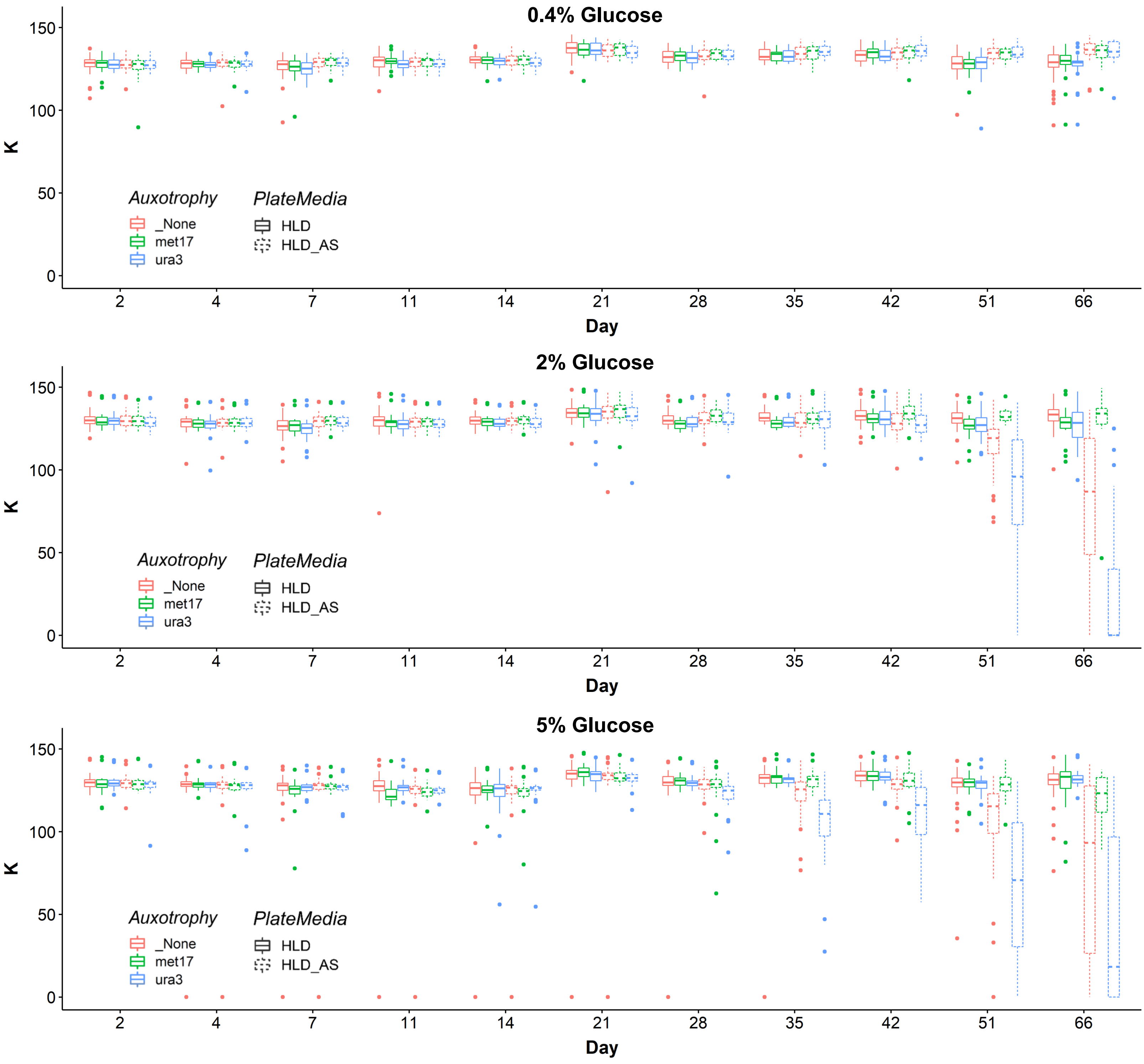
0.4% Glucose
K
150
100
50
0
Auxotrophy
_None
met17
ura3
PlateMedia
HLD
HLD_AS
2
4
7
11
14
21
28
35
42
51
66
Day
2% Glucose
K
150
100
50
0
Auxotrophy
_None
met17
ura3
PlateMedia
HLD
HLD_AS
2
4
7
11
14
21
28
35
42
51
66
Day
5% Glucose
K
150
100
50
0
Auxotrophy
_None
met17
ura3
PlateMedia
HLD
HLD_AS
2
4
7
11
14
21
28
35
42
51
66
Day

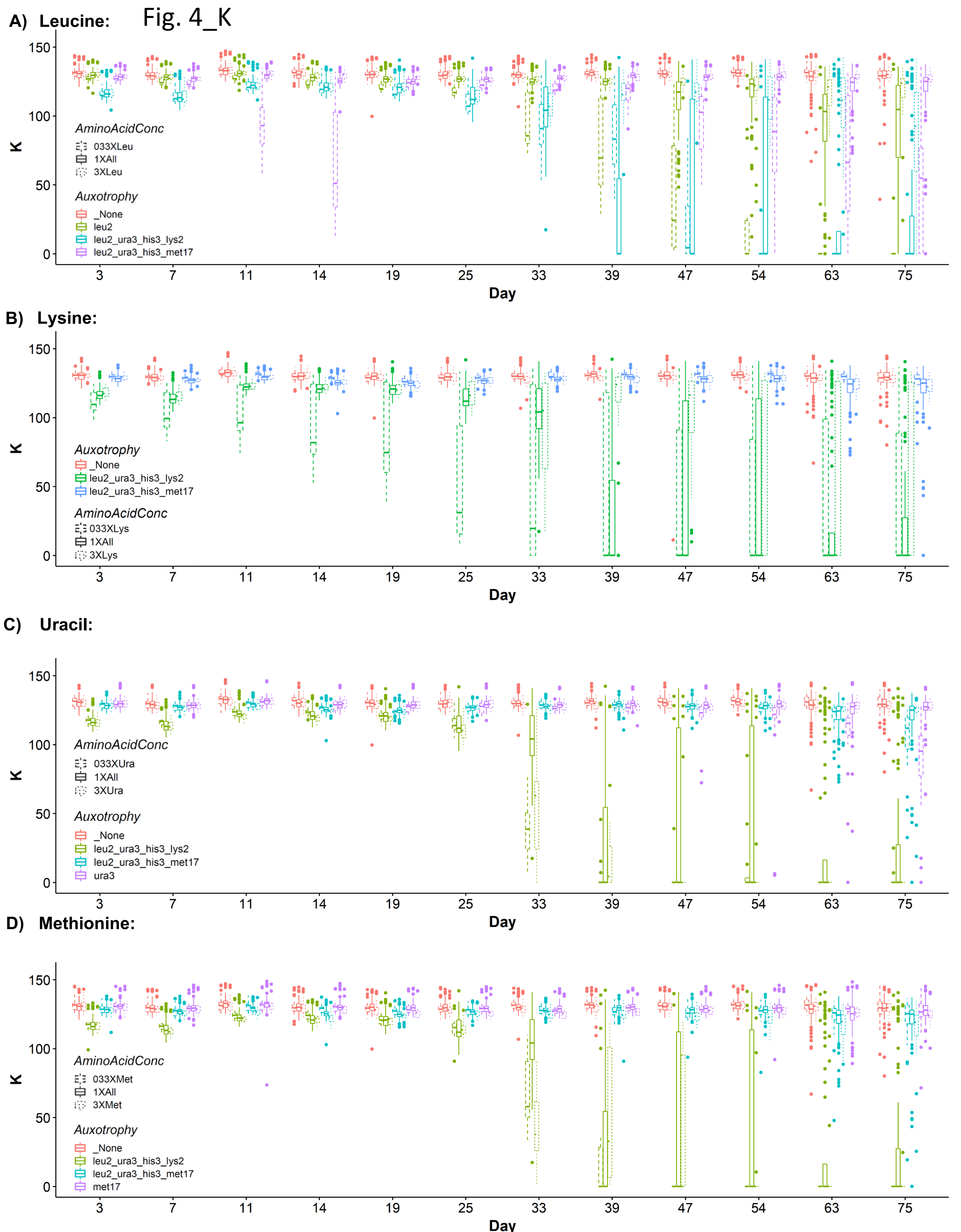
A) Leucine:
Fig. 4_K
K
Day
3
7
11
14
19
25
33
39
47
54
63
75
AminoAcidConc
033XLeu
1XAll
3XLeu
Auxotrophy
_None
leu2
leu2_ura3_his3_lys2
leu2_ura3_his3_met17
B) Lysine:
Auxotrophy
_None
leu2_ura3_his3_lys2
leu2_ura3_his3_met17
AminoAcidConc
033XLys
1XAll
3XLys
C) Uracil:
AminoAcidConc
033XUra
1XAll
3XUra
Auxotrophy
_None
leu2_ura3_his3_lys2
leu2_ura3_his3_met17
ura3
D) Methionine:
AminoAcidConc
033XMet
1XAll
3XMet
Auxotrophy
_None
leu2_ura3_his3_lys2
leu2_ura3_his3_met17
met17

Fig. S4_K

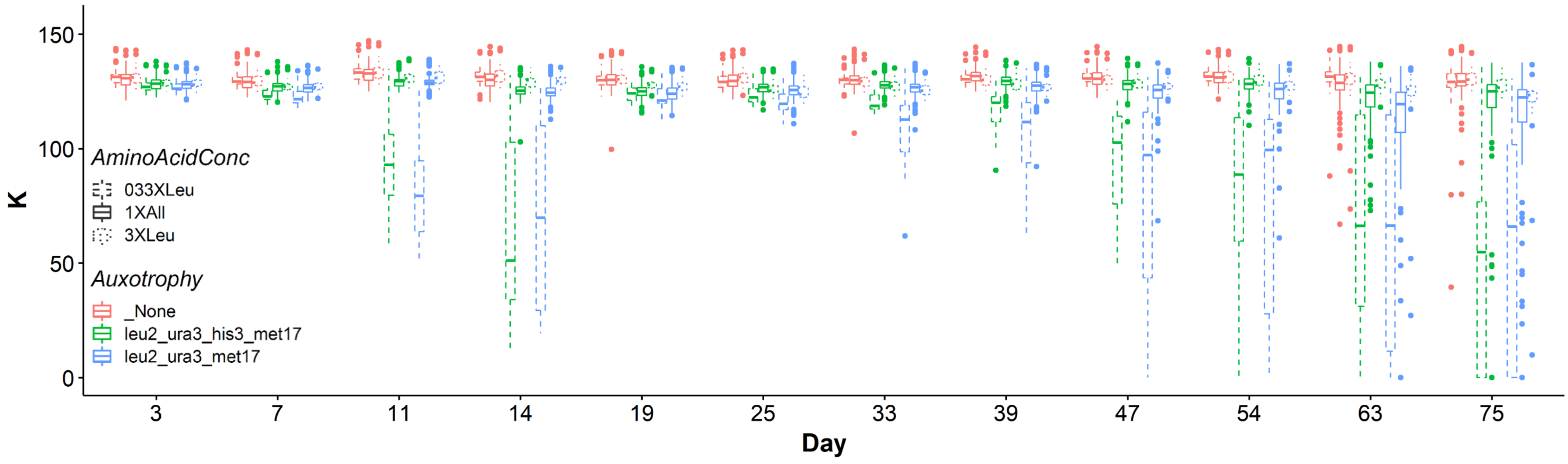

150
100
50
0
K
AminoAcidConc
033XLeu
1XAll
3XLeu
Auxotrophy
_None
leu2_ura3_his3_met17
leu2_ura3_met17
3
7
11
14
19
25
33
39
47
54
63
75
Day

Fig. S5_K

A) Leucine:

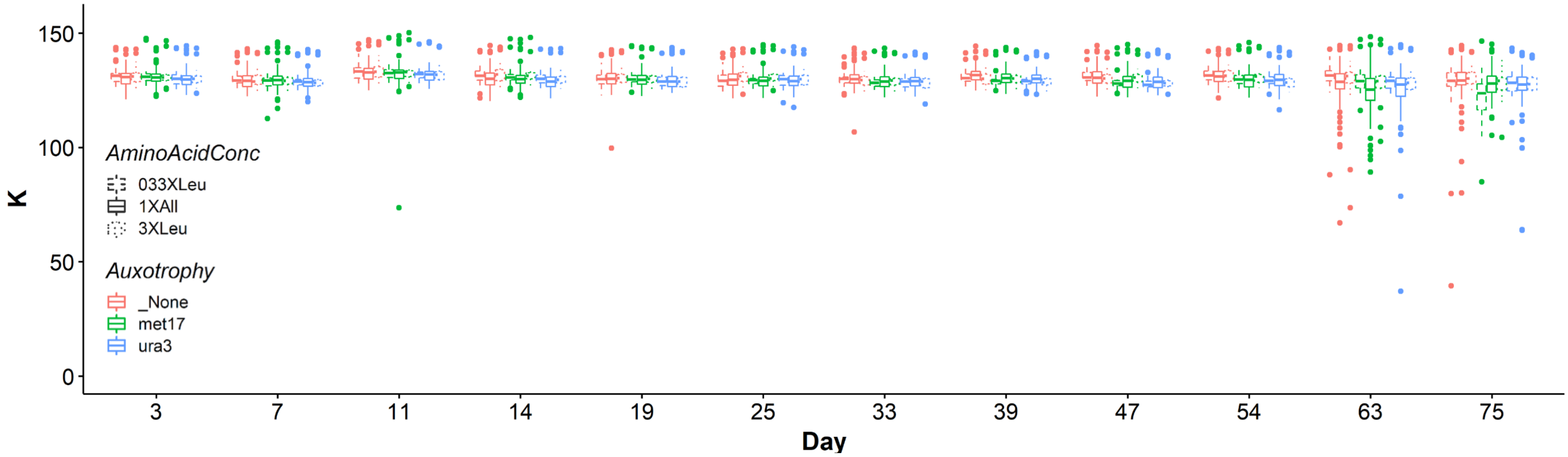


B) Lysine:

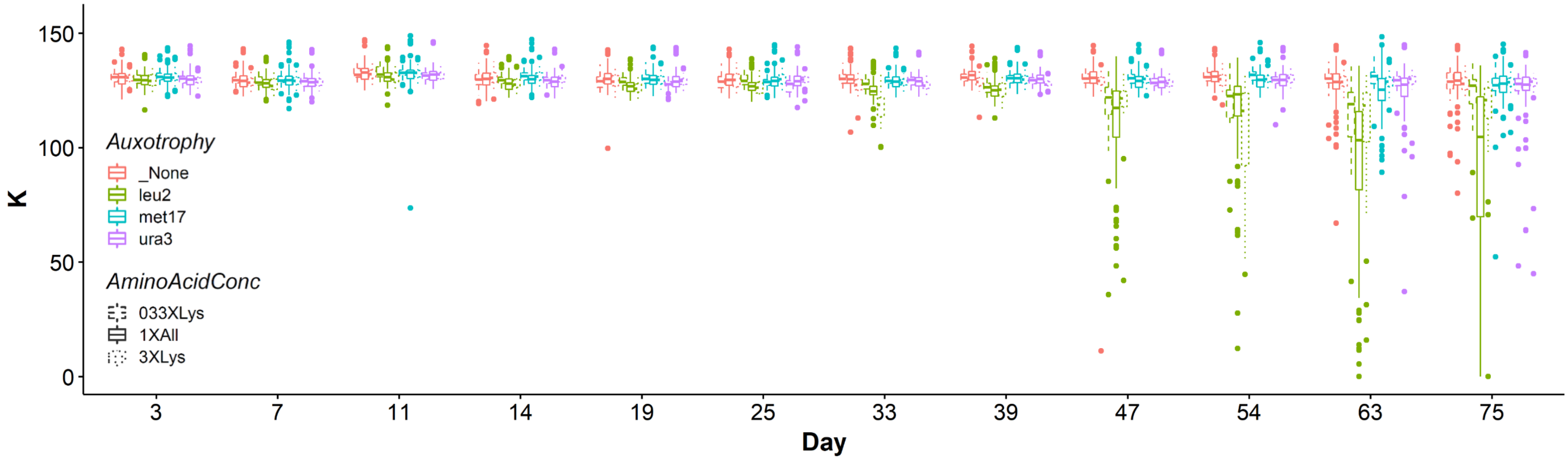


C) Uracil:

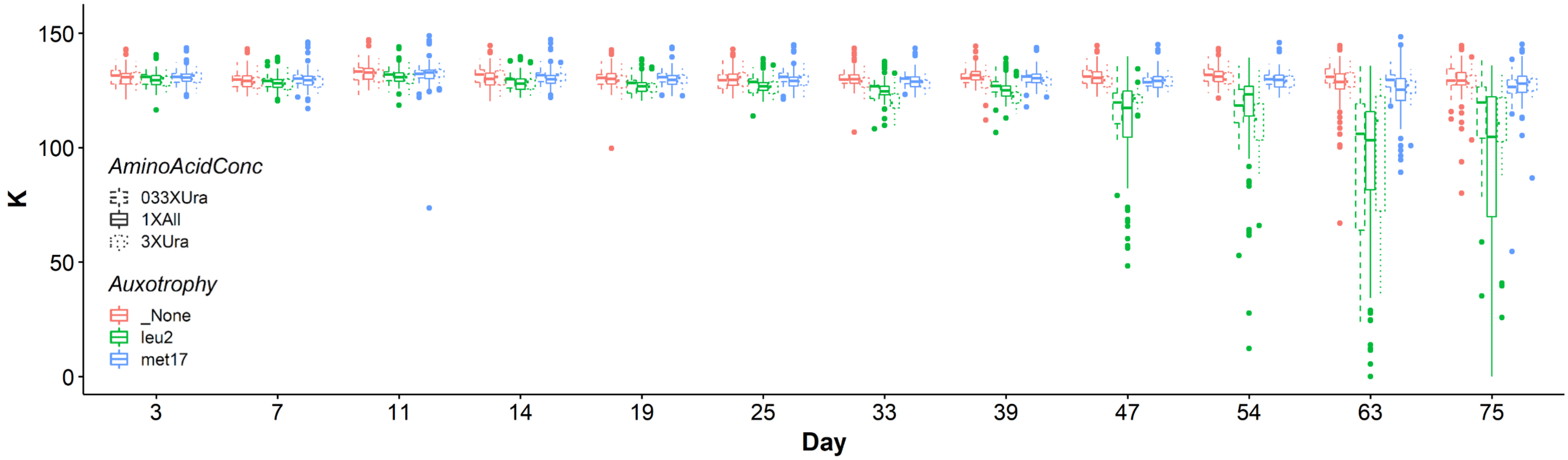


D) Methionine:

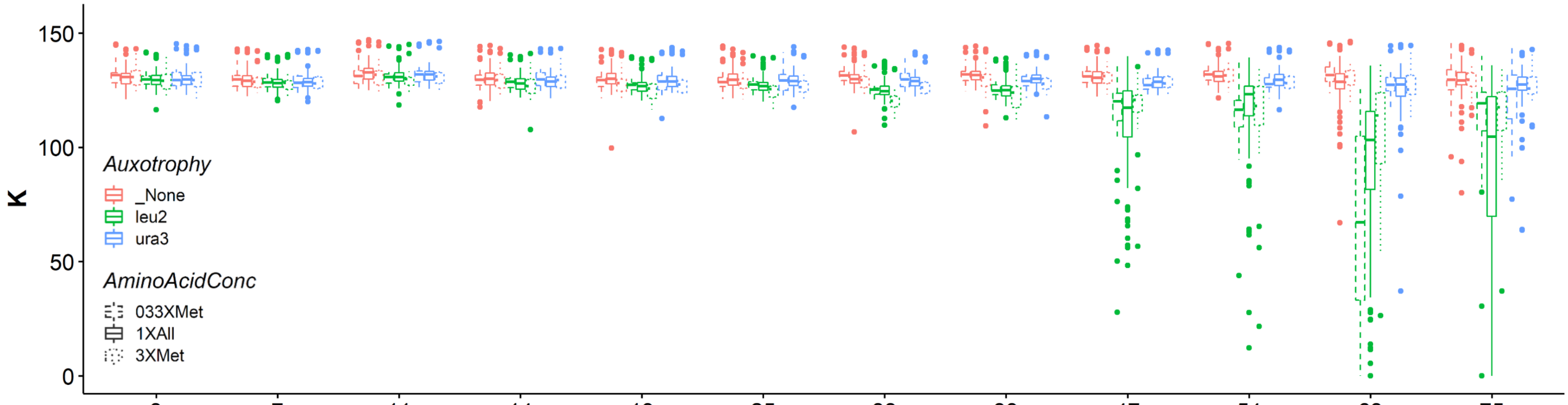

Fig. S6_K

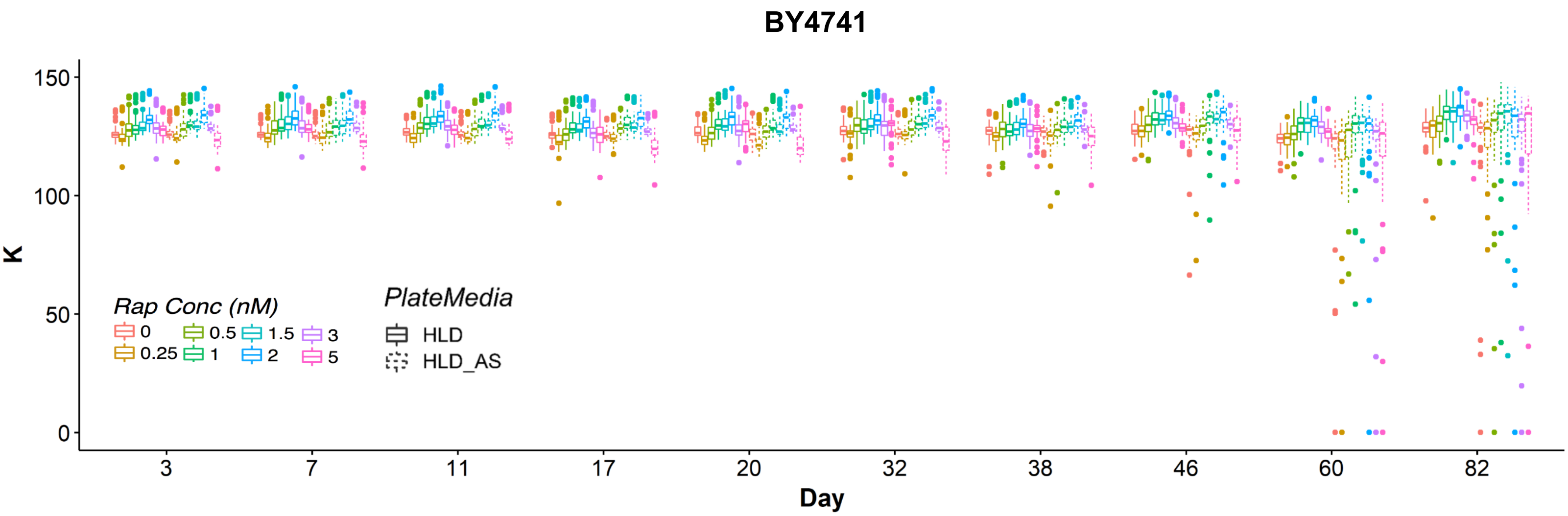

BY4741
150
100
50
0
K
Rap Conc (nM)
0
0.25
0.5
1
1.5
2
3
5
PlateMedia
HLD
HLD_AS
3
7
11
17
20
32
38
46
60
82
Day


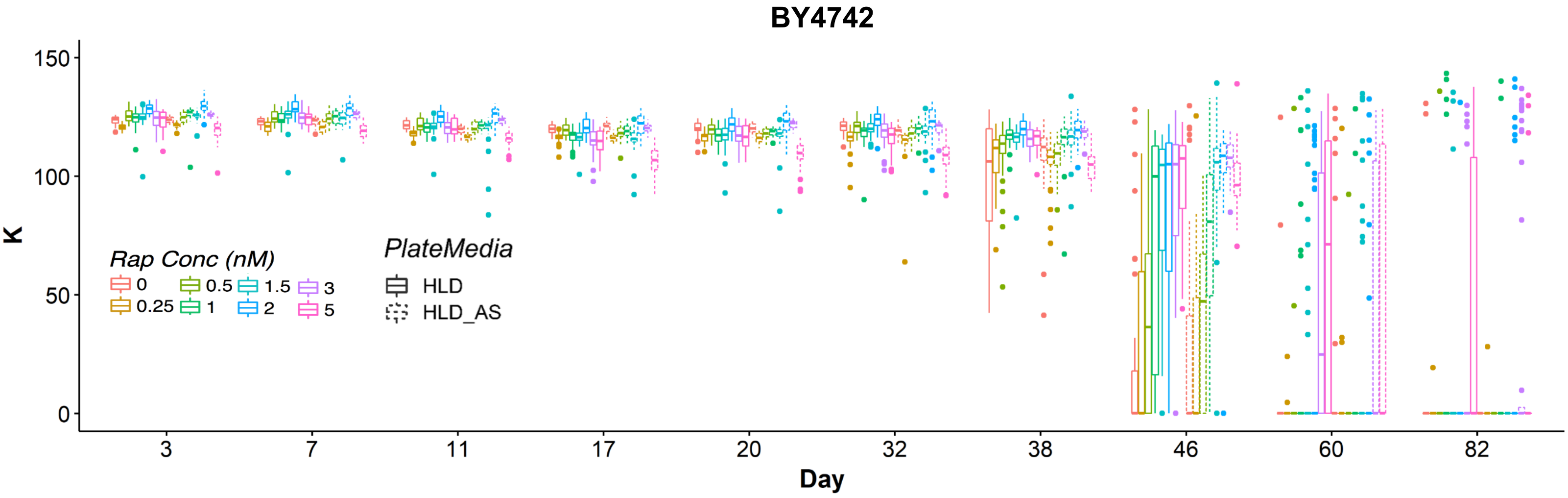

BY4742
150
100
50
0
K
Rap Conc (nM)
0
0.25
0.5
1
1.5
2
3
5
PlateMedia
HLD
HLD_AS
3
7
11
17
20
32
38
46
60
82
Day


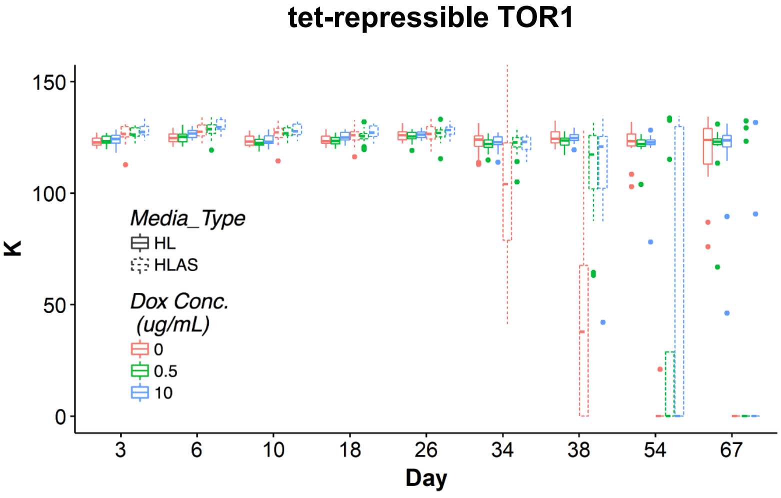

tet-repressible TOR1
150
100
50
0
K
Media_Type
HL
HLAS
Dox Conc.
(ug/mL)
0
0.5
10
3
6
10
18
26
34
38
54
67
Day

### Fig. S7_K

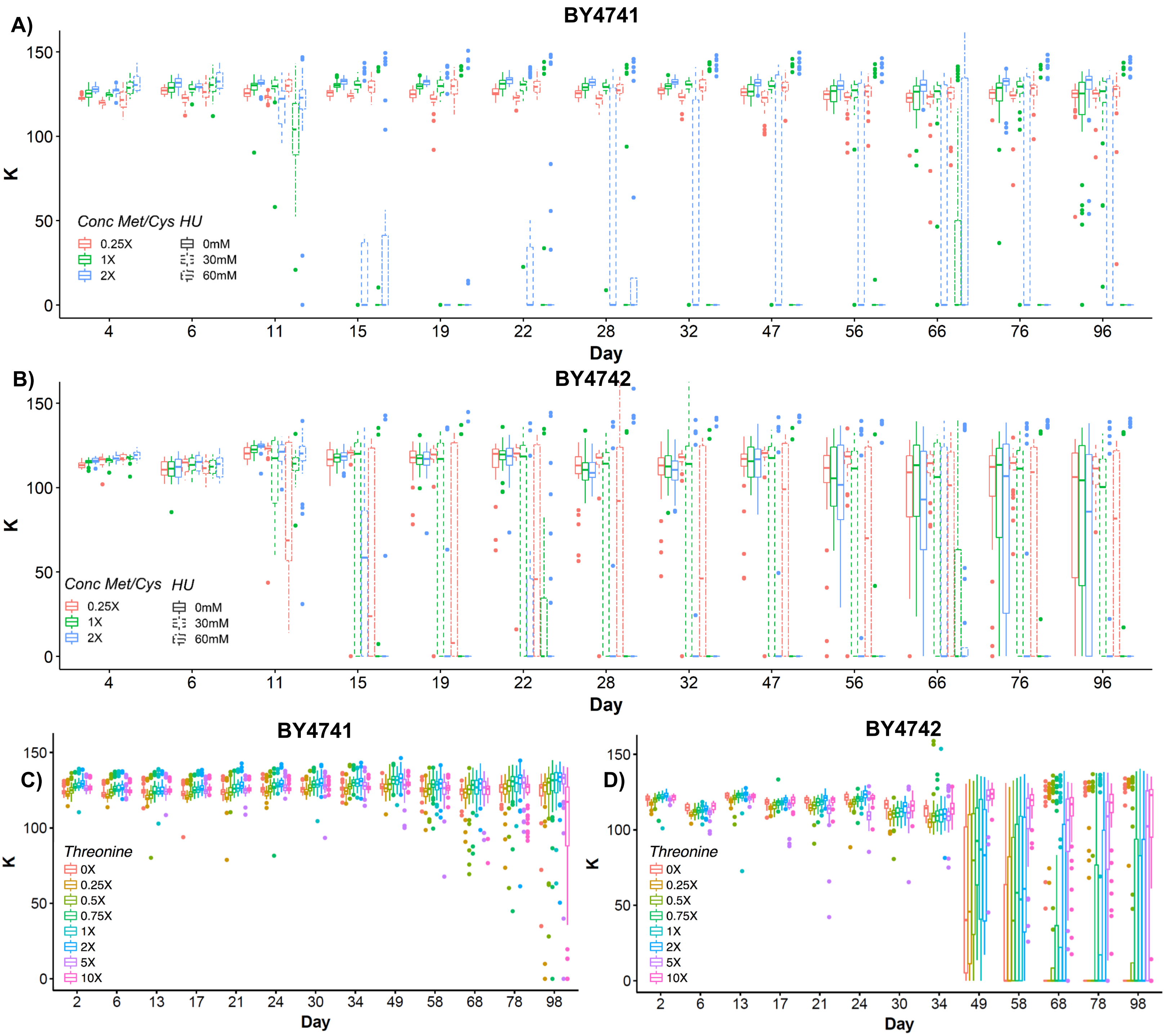

A)
BY4741
K
Day
Conc Met/Cys HU
0.25X
1X
2X
0mM
30mM
60mM
4
6
11
15
19
22
28
32
47
56
66
76
96
B)
BY4742
C)
BY4741
Threonine
0X
0.25X
0.5X
0.75X
1X
2X
5X
10X
2
6
13
17
21
24
30
34
49
58
68
78
98
D)
BY4742
