## Supplementary material for "High-resolution yeast quiescence profiling in human-like media reveals interacting influences of auxotrophy and nutrient availability": Online Resource 4

Online Resource 4 – Additional Figures – See Online Resource 1 for legends.

Fig. S1

A

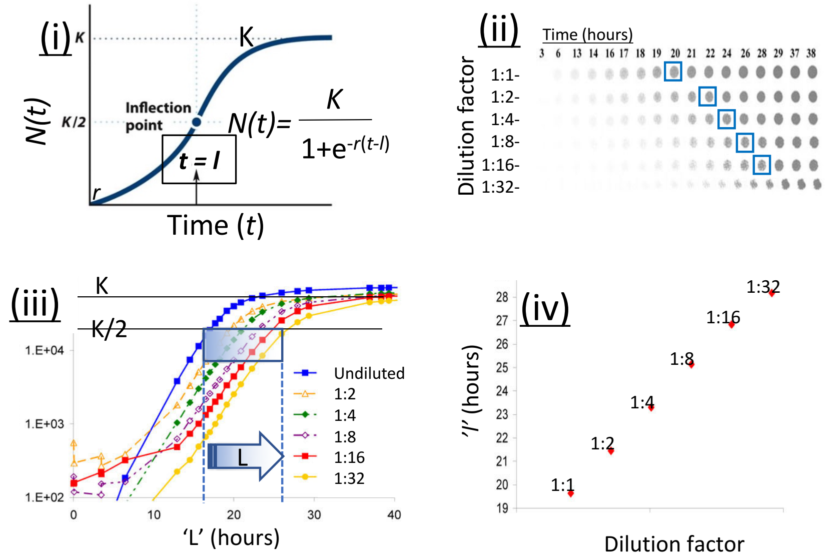


(v)

| L | %viable |
|---|---|
| 15 | 100 |
| 16.5 | 50 |
| 18 | 25 |
| 19.5 | 12.5 |
| 21 | 6.25 |
| 22.5 | 3.125 |
| 24 | 1.563 |
| 25.5 | 0.781 |
| 27 | 0.391 |
| 28.5 | 0.195 |
| 30 | 0.098 |
| 31.5 | 0.049 |
| 33 | 0.024 |

B

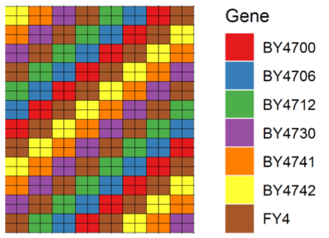


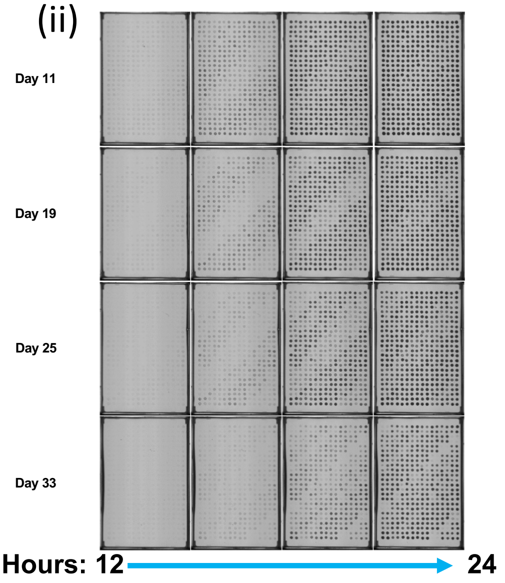


(iii)

| Strain | *MAT* | *HIS3/his3* | *LEU2/leu2* | *LYS2/lys2* | *MET17/met17* | *URA3/ura3* |
|---|---|---|---|---|---|---|
| **BY4700** | **a** | *HIS3* | *LEU2* | *LYS2* | *MET17* | *ura3* |
| **BY4706** | **a** | *HIS3* | *LEU2* | *LYS2* | *met17* | *URA3* |
| **BY4712** | **a** | *HIS3* | *leu2* | *LYS2* | *MET17* | *URA3* |
| **BY4730** | **a** | *HIS3* | *leu2* | *LYS2* | *met17* | *ura3* |
| **BY4741** | **a** | *HIS3* | *leu2* | *LYS2* | *met17* | *ura3* |
| **BY4742** | **a** | *HIS3* | *leu2* | *lys2* | *MET17* | *ura3* |
| **FY4** | **a** | *HIS3* | *LEU2* | *LYS2* | *MET17* | *URA3* |

Fig. S1, cont'd

C

(i)

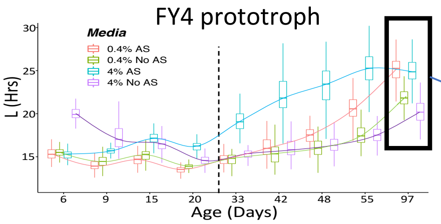


(ii)

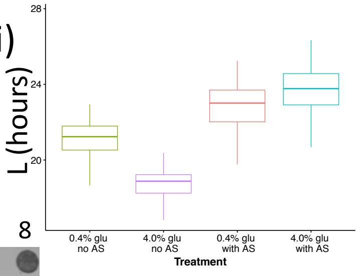


(iii)

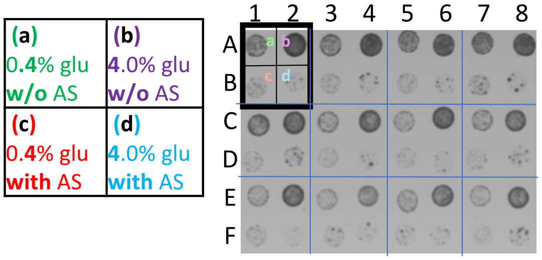


(iv)

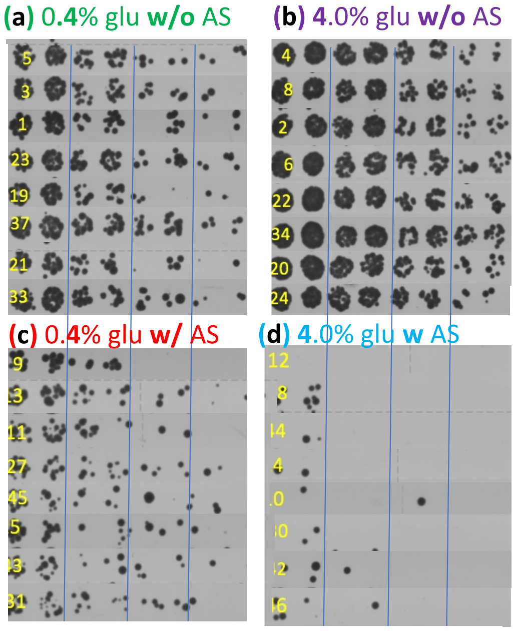


(v)

| | Q-HTCP Array pos. (panel C) | Ser.Dil Spot# | Avg. L | Stdev. L |
|---|---|---|---|---|
| (a) | A5 | 5 | **20.0** | 0.02 |
| | A3 | 3 | **20.1** | 0.41 |
| | A1 | 1 | **20.1** | 0.65 |
| | C7 | 23 | **21.0** | 0.29 |
| | C3 | 19 | **21.0** | 0.51 |
| | E5 | 37 | **21.4** | 0.14 |
| | C5 | 21 | **21.5** | 0.42 |
| | E1 | 33 | **21.9** | 0.23 |
| (b) | A4 | 4 | **17.7** | 0.20 |
| | A8 | 8 | **17.7** | 0.60 |
| | A2 | 2 | **17.8** | 0.04 |
| | A6 | 6 | **18.0** | 0.54 |
| | C6 | 22 | **18.5** | 0.34 |
| | E2 | 34 | **18.7** | 0.55 |
| | C4 | 20 | **19.0** | 0.29 |
| | C8 | 24 | **19.2** | 0.27 |
| (c) | B1 | 9 | **22.4** | 0.46 |
| | B5 | 13 | **22.7** | 0.97 |
| | B3 | 11 | **22.7** | 0.75 |
| | D3 | 27 | **22.9** | 0.77 |
| | F5 | 45 | **23.2** | 0.27 |
| | D1 | 25 | **23.3** | 1.15 |
| | F3 | 43 | **23.4** | 0.13 |
| | D7 | 31 | **23.4** | 0.81 |
| (d) | B4 | 12 | **23.6** | 0.20 |
| | D4 | 28 | **23.7** | 0.11 |
| | F4 | 44 | **23.8** | 1.27 |
| | B6 | 14 | **24.0** | 0.70 |
| | B2 | 10 | **24.1** | 0.81 |
| | D6 | 30 | **24.3** | 2.45 |
| | F2 | 42 | **24.8** | 3.51 |
| | F6 | 46 | **25.0** | 0.35 |

Fig. S2

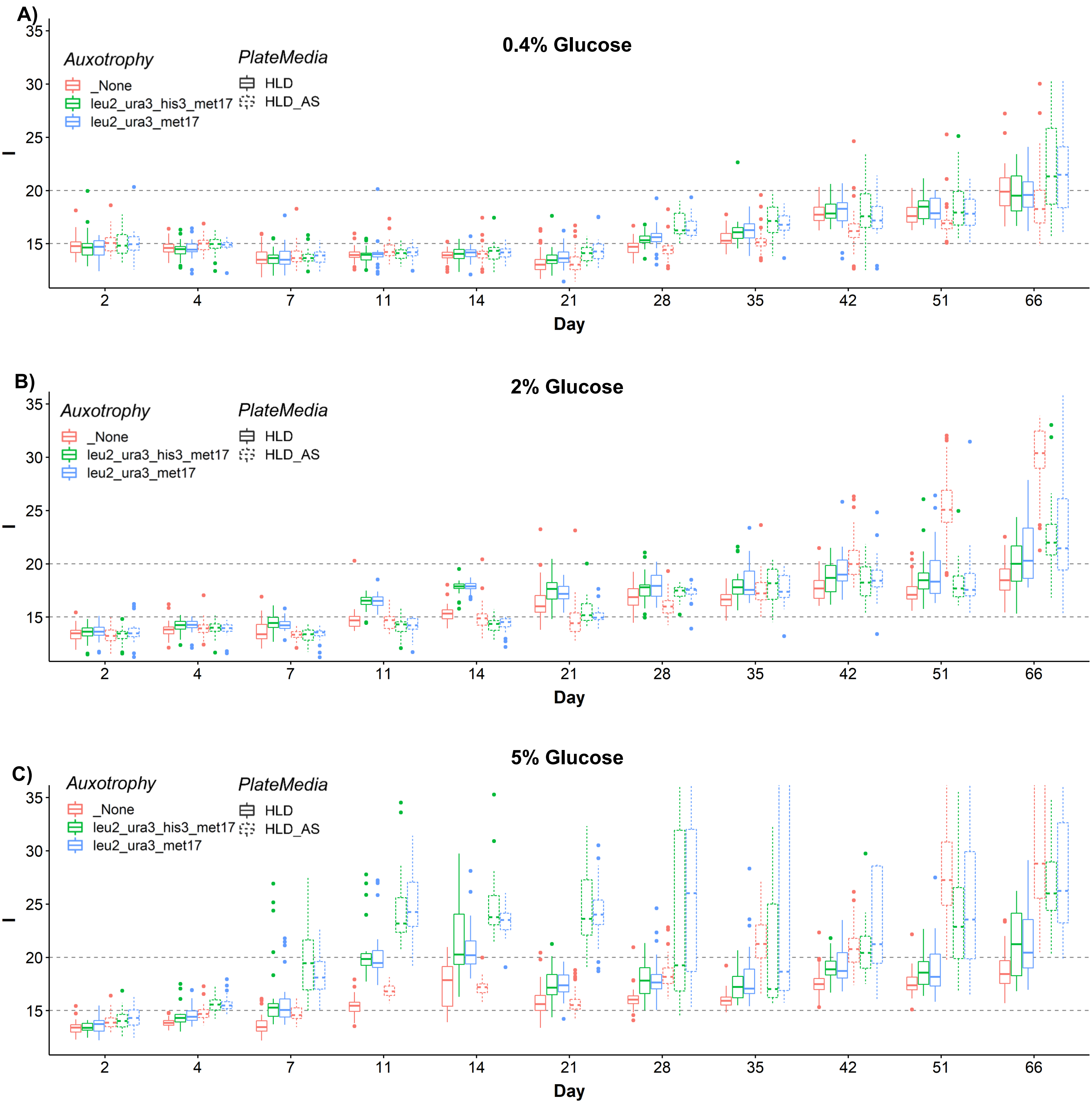
A)
0.4% Glucose
Auxotrophy
_None
leu2_ura3_his3_met17
leu2_ura3_met17
PlateMedia
HLD
HLD_AS
35
30
25
20
15
2
4
7
11
14
21
28
35
42
51
66
Day
B)
2% Glucose
Auxotrophy
_None
leu2_ura3_his3_met17
leu2_ura3_met17
PlateMedia
HLD
HLD_AS
Day
C)
5% Glucose
Auxotrophy
_None
leu2_ura3_his3_met17
leu2_ura3_met17
PlateMedia
HLD
HLD_AS
Day

Fig. S3

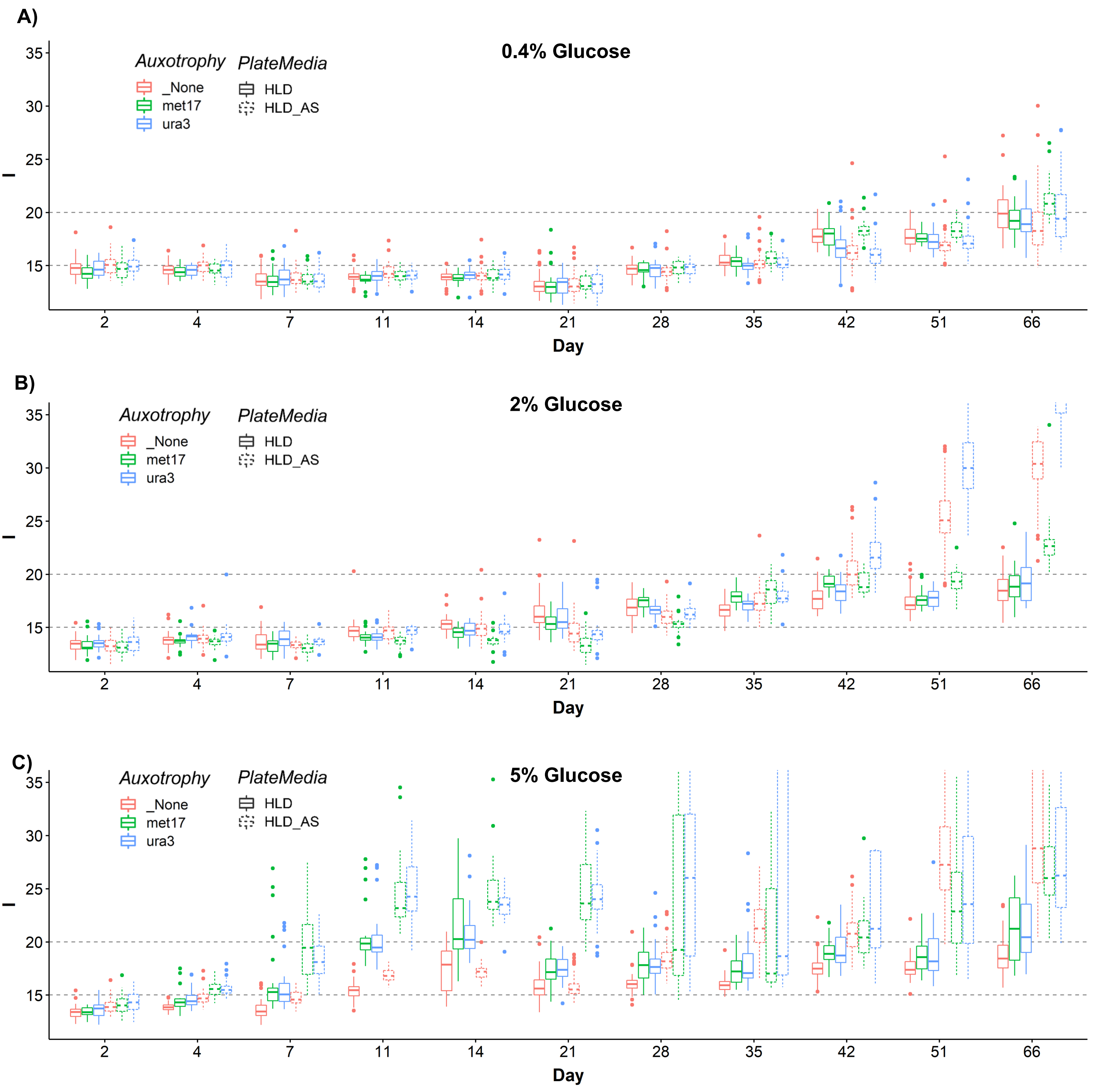
A)
0.4% Glucose
Auxotrophy
_None
met17
ura3
PlateMedia
HLD
HLD_AS
35
30
25
20
15
2
4
7
11
14
21
28
35
42
51
66
Day
B)
2% Glucose
Auxotrophy
_None
met17
ura3
PlateMedia
HLD
HLD_AS
Day
C)
5% Glucose
Auxotrophy
_None
met17
ura3
PlateMedia
HLD
HLD_AS
Day

### Fig. S4

**Leucine:**

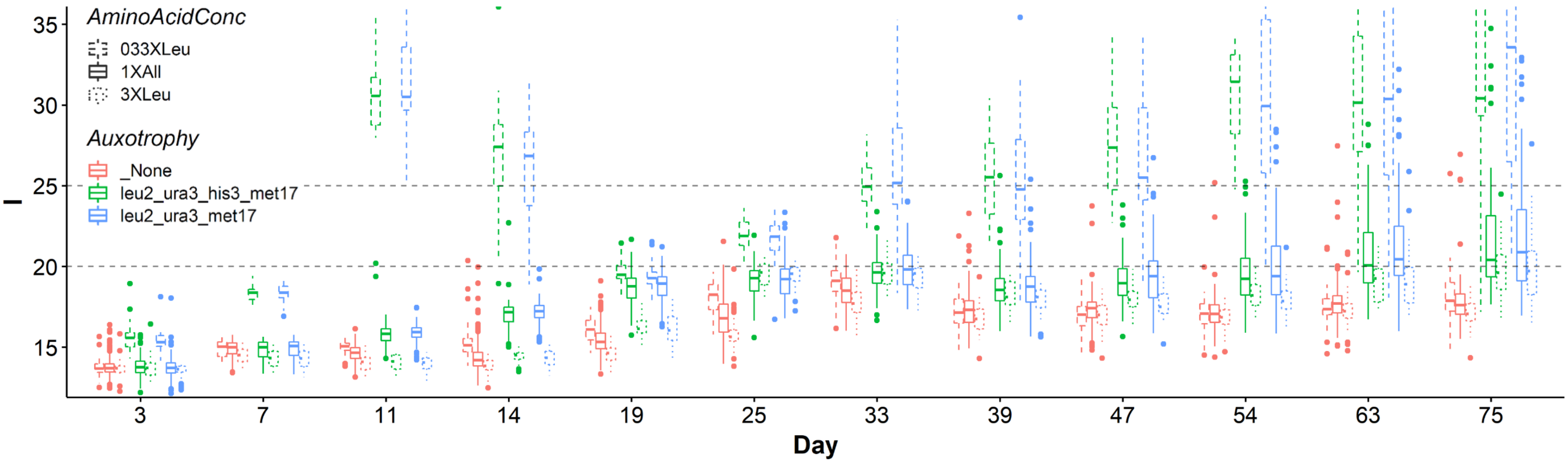

Fig. S5

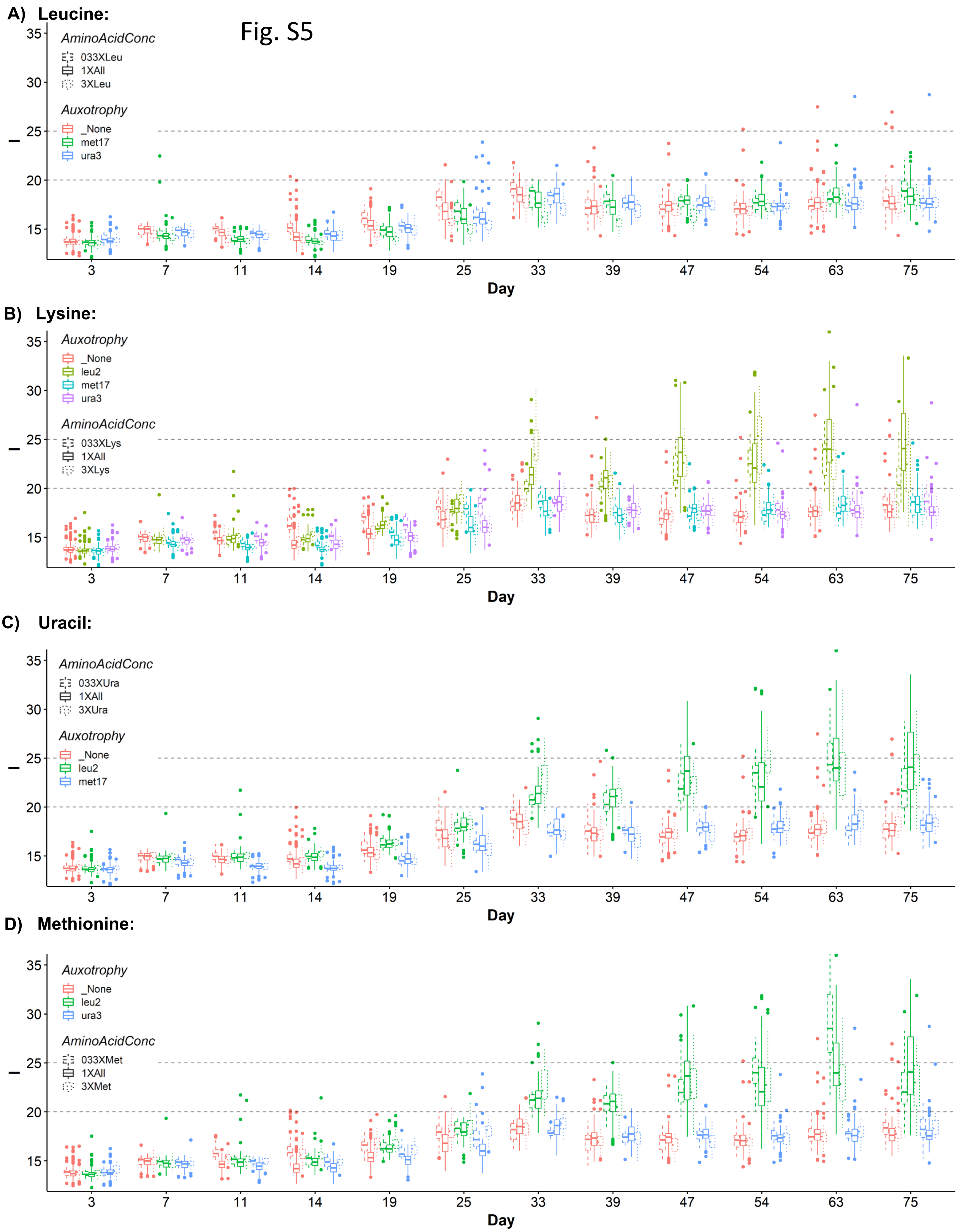
A) Leucine:
AminoAcidConc
033XLeu
1XAll
3XLeu
Auxotrophy
_None
met17
ura3
15
20
25
30
35
3
7
11
14
19
25
33
39
47
54
63
75
Day
B) Lysine:
Auxotrophy
_None
leu2
met17
ura3
AminoAcidConc
033XLys
1XAll
3XLys
Day
C) Uracil:
AminoAcidConc
033XUra
1XAll
3XUra
Auxotrophy
_None
leu2
met17
Day
D) Methionine:
Auxotrophy
_None
leu2
ura3
AminoAcidConc
033XMet
1XAll
3XMet
Day

Fig. S6

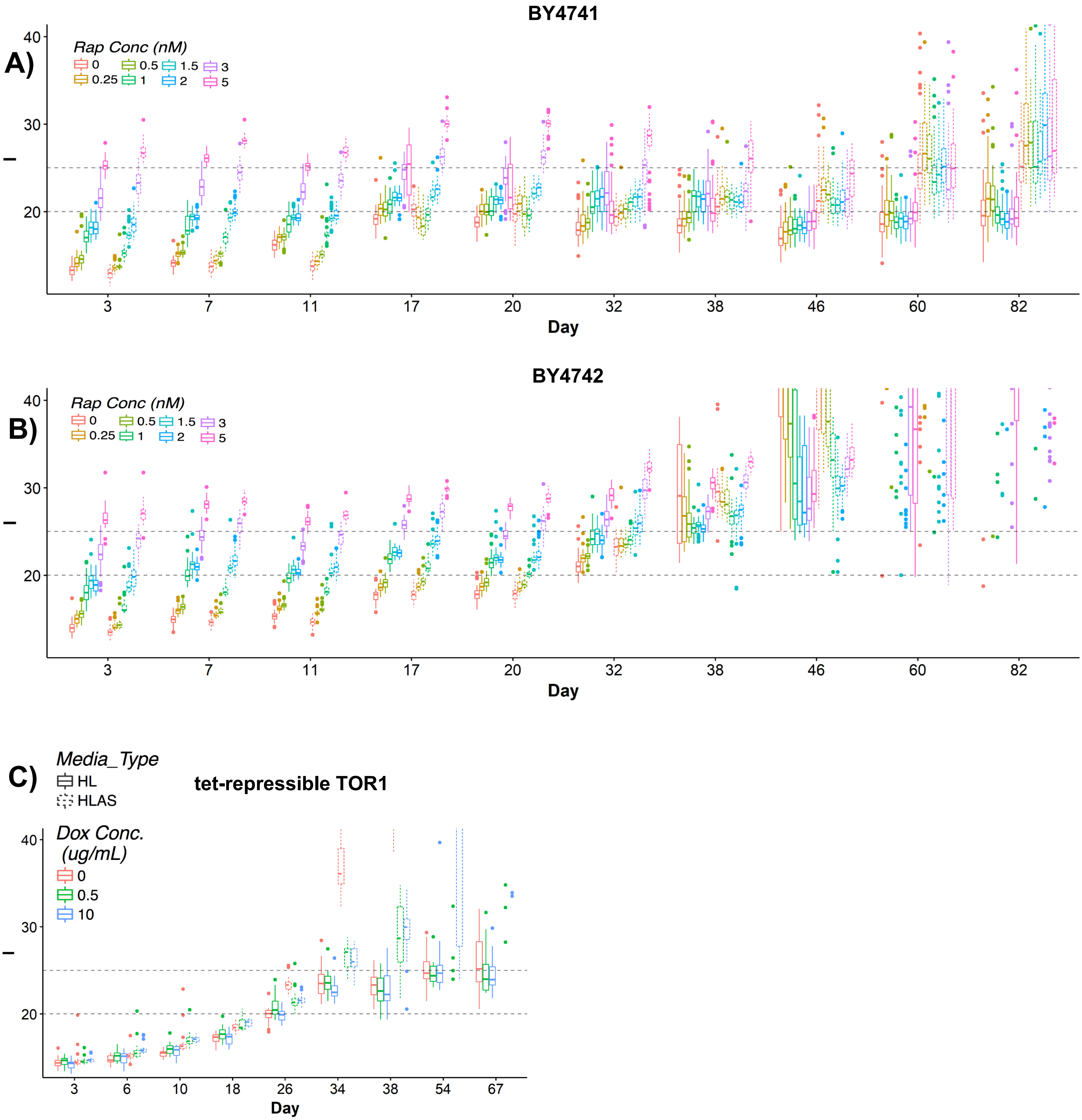
A)
BY4741
Rap Conc (nM)
0
0.25
0.5
1
1.5
2
3
5
40
30
20
3
7
11
17
20
32
38
46
60
82
Day
B)
BY4742
Rap Conc (nM)
0
0.25
0.5
1
1.5
2
3
5
40
30
20
3
7
11
17
20
32
38
46
60
82
Day
C)
Media_Type
HL
HLAS
tet-repressible TOR1
Dox Conc.
(ug/mL)
0
0.5
10
40
30
20
3
6
10
18
26
34
38
54
67
Day

Fig. S7

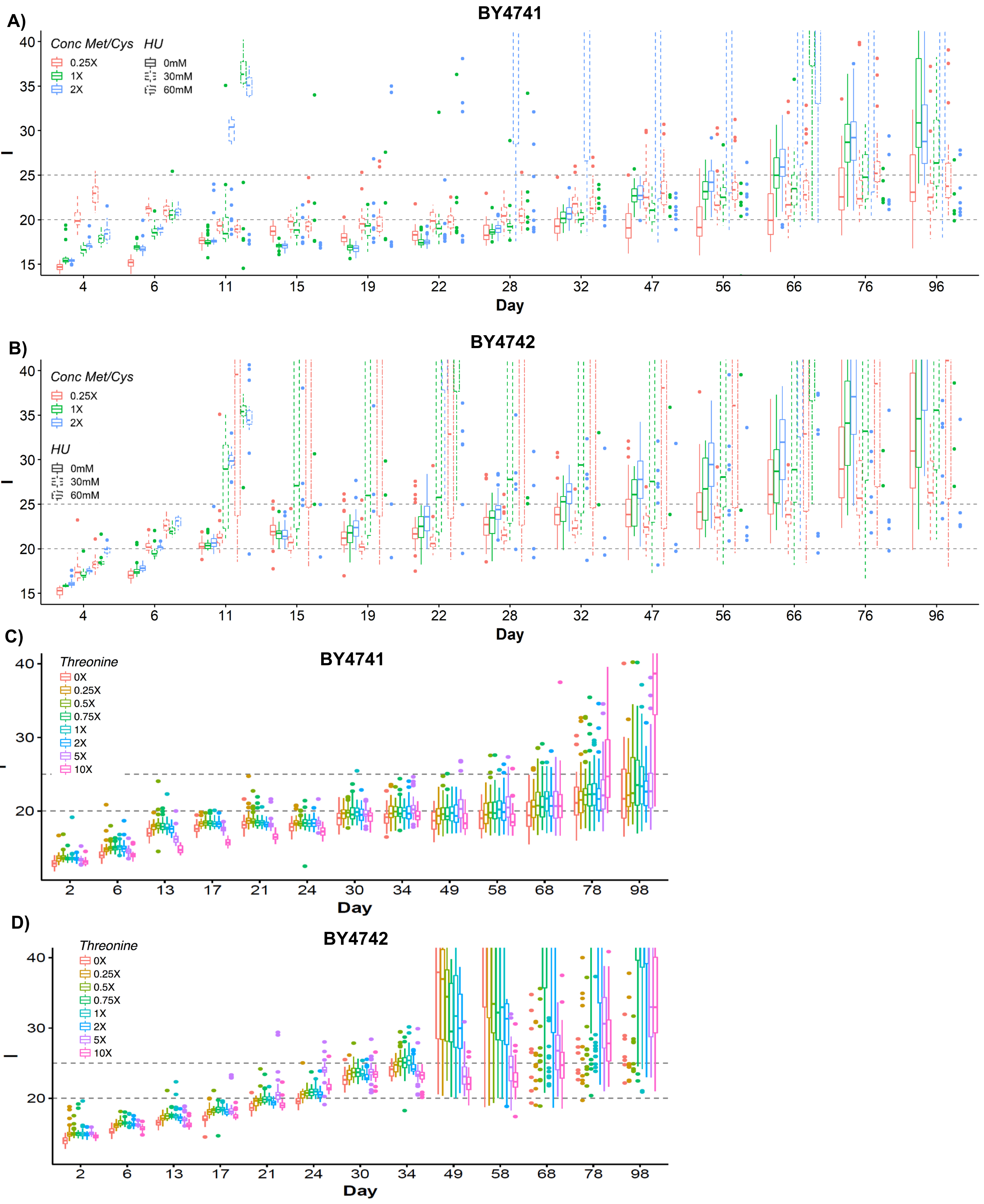
A)
BY4741
Conc Met/Cys
0.25X
1X
2X
HU
0mM
30mM
60mM
40
35
30
25
20
15
4
6
11
15
19
22
28
32
47
56
66
76
96
Day
B)
BY4742
Conc Met/Cys
0.25X
1X
2X
HU
0mM
30mM
60mM
40
35
30
25
20
15
4
6
11
15
19
22
28
32
47
56
66
76
96
Day
C)
BY4741
Threonine
0X
0.25X
0.5X
0.75X
1X
2X
5X
10X
40
30
20
2
6
13
17
21
24
30
34
49
58
68
78
98
Day
D)
BY4742
Threonine
0X
0.25X
0.5X
0.75X
1X
2X
5X
10X
40
30
20
2
6
13
17
21
24
30
34
49
58
68
78
98
Day
